# Subcellular proteomics and iPSC modeling uncover reversible mechanisms of axonal pathology in Alzheimer’s disease

**DOI:** 10.1101/2022.09.30.510408

**Authors:** Yifei Cai, Jean Kanyo, Rashaun Wilson, Shveta Bathla, Pablo Leal Cardozo, Lei Tong, Shanshan Qin, Lukas A. Fuentes, Iguaracy Pinheiro-de-Sousa, Tram Huynh, Liyuan Sun, Mohammad Shahid Mansuri, Zichen Tian, Hao-Ran Gan, Amber Braker, Hoang Kim Trinh, Anita Huttner, TuKiet T. Lam, Evangelia Petsalaki, Kristen J. Brennand, Angus C. Nairn, Jaime Grutzendler

**Affiliations:** Department of Neurology, Yale University, New Haven, CT 06511; Keck MS & Proteomics Resource, Yale University, New Haven, CT, 06511; Yale/NIDA Neuroproteomics Center, Yale University, New Haven, CT 06511; Department of Psychiatry, Yale University, New Haven, CT 06511; John A. Paulson School of Engineering and Applied Sciences, Harvard University, Cambridge, MA, USA 02134; Department of Cell Biology, Yale University School of Medicine, New Haven, CT, 06511; European Molecular Biology Laboratory, European Bioinformatics Institute, Hinxton, UK; Yale College, Department of Neuroscience, Yale University, New Haven, CT 06520; Department of Pathology, Yale University, New Haven, CT 06520; Department of Molecular Biophysics and Biochemistry, Yale University, New Haven, CT, 06511; Department of Genetics, Yale University, New Haven, CT 06520; Department of Pharmacology, Yale University, New Haven, CT 06520; Department of Neuroscience, Yale University, New Haven, CT 06511

## Abstract

Axonal spheroids (dystrophic neurites) are commonly found around amyloid deposits in Alzheimer’s disease (AD). They impair electrical conduction, disrupt neural circuits, and correlate with AD severity. Despite their significance, the mechanisms underlying spheroid formation remain unknown. To address this, we developed a proximity labeling proteomics approach to uncover the proteome of spheroids in human postmortem and mouse brains. Additionally, we established a human iPSC-derived AD model allowing mechanistic investigation of spheroid pathology and optical electrophysiology. This approach revealed the subcellular molecular architecture of spheroids and identified abnormalities in key biological processes, including protein turnover, cytoskeleton dynamics, and lipid transport. Notably, the PI3K/AKT/mTOR pathway, which regulates these processes, was activated within spheroids. Furthermore, phosphorylated mTOR levels in spheroids strongly correlated with AD severity in humans. Importantly, inhibition of mTOR in iPSC-derived neurons and in mice ameliorated spheroid pathology. Altogether, our study provides a multidisciplinary toolkit for investigating mechanisms and novel targets for axonal pathology in neurodegeneration.

## INTRODUCTION

A major pathological hallmark in Alzheimer’s disease (AD) is the accumulation of aggregated extracellular β-amyloid (Aβ) peptide deposits [1]. However, the precise mechanisms through which these deposits trigger downstream neuronal changes and contribute to cognitive deficits remain unknown. While amyloid plaques have been shown to cause overall synapse loss [2] and dendritic spine reduction in their vicinity [3], a less understood, yet potentially critical pathological hallmark are the axonal spheroids that develop around plaques. Specifically, we and others have shown that hundreds of axons but not dendrites in the vicinity of individual amyloid plaques develop enlarged spheroid-like structures (traditionally termed dystrophic neurites) [4–10]. These plaque-associated axonal spheroids (PAAS) have been shown to correlate with AD severity [4, 11] and markedly disrupt axonal electrical conduction [4, 12, 13] and neuronal network function [4], potentially leading to cognitive impairment. PAAS contain large numbers of enlarged and enzyme-deficient endolysosomal vesicles [4, 14–16], and autophagosomes [15, 17]. Spheroid enlargement may be driven by the accumulation of these vesicles [4], and by the disruption of axonal cytoskeleton and transport [16, 18–22]. Ultimately, the presence of axonal spheroids may further impair axonal trafficking, leading to downstream synaptic dysfunction and axonal degeneration [3, 23–25]. Additionally, it has been proposed that PAAS may play a role in the propagation of tau pathology through neuronal networks [26]. Therefore, PAAS could be a critical neuropathological hub implicated in circuit disruption and proteinopathy, contributing to cognitive decline in AD [3, 4, 12, 13, 23, 27].

Axonal spheroids can form as a result of a variety of insults and have been observed in many pathological processes, ranging from acute neural injury to age-related neurodegenerative conditions. These spheroids share morphological and subcellular cytoskeletal and organelle similarities, including the accumulation of certain proteins such as Amyloid Precursor Protein (APP) and Cathepsins [14, 22]. However, significant mechanistic differences exist given the diversity of pathological processes involved. For example, in models of neural injury, spheroid formation has been shown to depend on alterations in the cytoskeleton and disruption of axonal and membrane tension [22, 28]. This process is associated with phosphatidylserine exposure, leading to glial phagocytosis [22, 29]. Conversely, in Alzheimer’s disease, plaque-associated spheroids have been shown to be long-lasting and are not typically engulfed by phagocytes [4, 30–32].

Despite these observations, PAAS have not been a major focus of mechanistic investigations, and the cell biological processes involved in their formation remain poorly understood. In this study, we developed a proteomics approach to systematically investigate the molecular composition of PAAS, by employing proximity labeling to selectively isolate the subcellular proteome of PAAS in human postmortem and mouse brain. Using this approach, we uncovered the presence of protein turnover, cytoskeleton dynamics and lipid transport as key biological processes in PAAS. Furthermore, we identified hundreds of previously unknown proteins and signaling pathways that are expressed in PAAS, some of which could play important roles in PAAS formation.

To investigate the structural dynamics, functional consequences, and potential reversibility of PAAS, we established a versatile human iPSC-derived AD model that recapitulates PAAS pathology. This model enabled us to conduct longitudinal imaging of PAAS and optical electrophysiology using calcium sensors in human neurons, revealing the growth patterns and disruption in action potential conduction caused by PAAS. To further investigate the mechanisms of spheroid growth, we focused on the mTOR signaling pathway which was identified in the PAAS proteomics and confirmed to be expressed within PAAS. Both genetic and pharmacological inhibition of the mTOR pathway in human iPSC neurons and in mice led to a marked reduction in PAAS pathology. Given that mTOR is a master regulator of protein turnover, lipid metabolism and axonal cytoskeletal remodeling [33–35], these findings underscore the importance of these biological processes in PAAS formation.

Altogether, the integration of subcellular proteomics in postmortem human brain, human iPSC AD modeling, and molecular manipulation of PAAS in human neurons and mice, provides new insight into the complex cell biology and reversibility of axonal pathology in AD.

## RESULTS

### Proximity labeling of proteins within axonal spheroids in the human postmortem AD brain

Proximity labeling is a recently developed methodology used to biotinylate proteins within a specific cellular or subcellular compartment through genetic expression of localized peroxidases or biotin ligases in living cells and animals [36–40]. This method allows the selective protein isolation and subsequent identification of subcellular proteomes using liquid chromatography with tandem mass spectrometry (LC-MS/MS) technology [41, 42]. A modification of this approach was recently introduced enabling localized protein biotinylation in fixed tissues, by means of horseradish peroxidase (HRP)-conjugated antibodies targeted to a subcellular compartment [43–46]. Leveraging these methodologies, we devised and refined an antibody-based proximity labeling approach to uncover the proteome of PAAS in both human AD postmortem brains and AD-model mice (5XFAD) (**Figure 1; Figures S1-S4 and Movie S1**).

**Figure 1.**
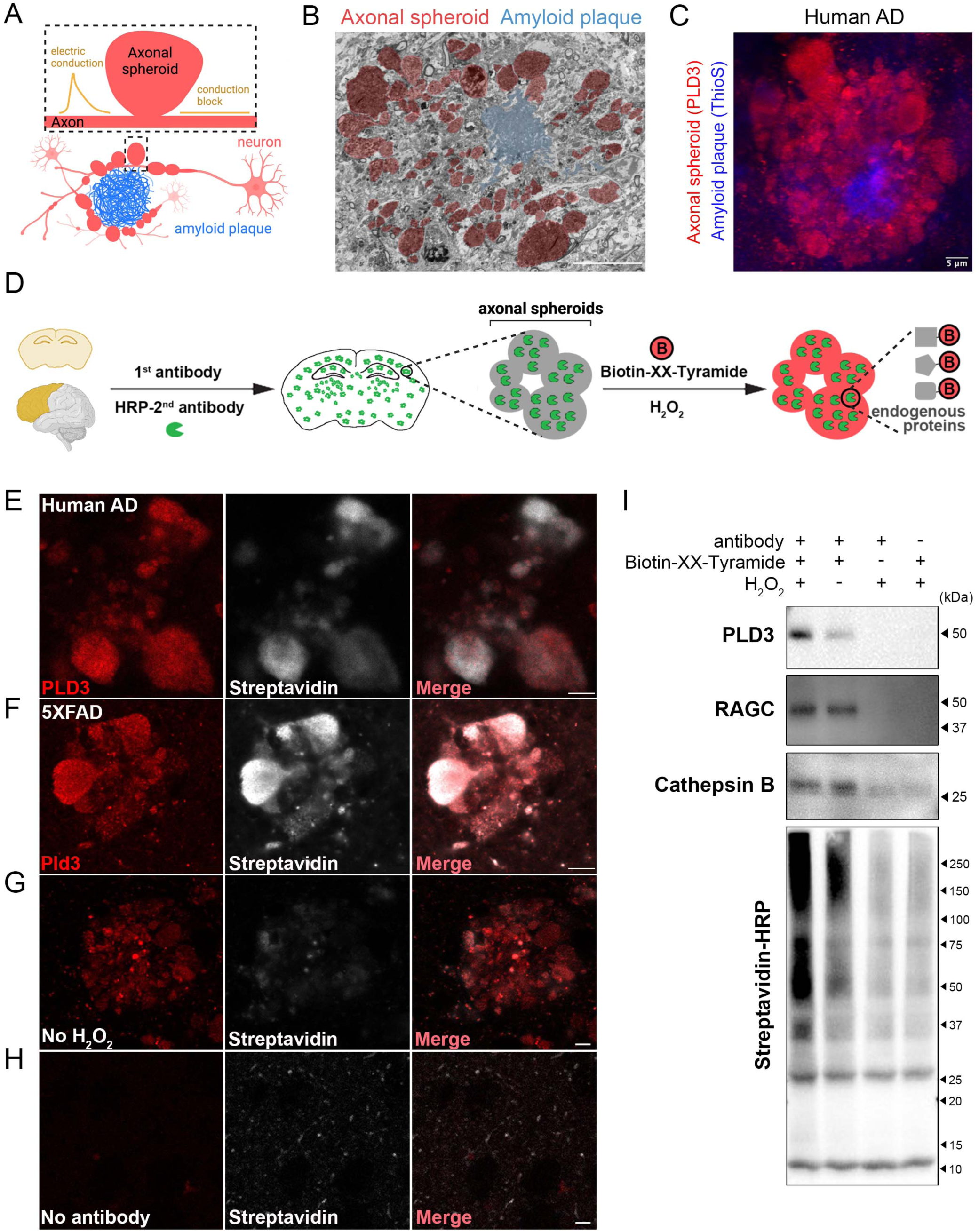
Proximity labeling of proteins within plaque-associated axonal spheroids. **A.** Schematic showing axons with spheroids (red) around an amyloid plaque (blue). Spheroids cause electric conduction delay and blockage along axons [4]. **B.** A FIB-SEM image from a 5XFAD mouse shows spheroids (red) around an amyloid plaque (blue). Scale bar = 20 μm. Related to **Movie S1**. **C.** Immunofluorescence confocal deconvolved image shows that PLD3 is highly enriched in axonal spheroids (red, PLD3) around amyloid plaques (blue, ThioflavinS) in postmortem AD human brain. **D.** A schematic showing the pipeline for proximity labeling PAAS proteomics in postmortem brains. Postmortem AD human or mouse brains were incubated with primary antibody against PLD3 and HRP-conjugated secondary antibody, followed by biotinylation reaction in the presence of Biotin-XX-Tyramide and hydrogen peroxide. **E-H.** Proximity labeling biotinylation of proteins within PAAS in **(E)** human AD and **(F)** 5XFAD mouse brains. **(E-F)** Biotinylated proteins were revealed by streptavidin-Alexa fluor 647. **(G-H)** Control conditions including **(G)** no H2O2 or **(H)** no-antibody labeling markedly reduced biotinylation. Scale bar = 5 μm. **I.** Streptavidin-HRP western blot demonstrates efficient streptavidin bead pull-down of biotinylated proteins, including the protein bait PLD3, and the known axonal spheroid proteins RAGC and Cathepsin B. See also **Figures S1-S4**.

Our approach was based on the observation that phospholipase D3 (PLD3), an endolysosomal protein, is highly abundant within PAAS (**Figures 1C and S2E-G**) [4, 47–50], and is specifically expressed in neurons [4, 47] and absent in glial cells [4]. Although low levels of PLD3 are also found in neuronal cell bodies (**Figure S2**), quantitative immunofluorescence demonstrated that the majority of PLD3 originates from PAAS (**Figure S2G**). Leveraging this finding, we used PLD3 as a protein bait for proximity labeling proteomics of PAAS. This involved the sequential incubation of postmortem AD human or mouse brains with primary antibody against PLD3 and secondary HRP-conjugated antibody, followed by a peroxidation reaction in the presence of hydrogen peroxide and Biotin-XX-Tyramide (**Figure 1D**). This resulted in robust biotinylation of proteins contained within PAAS, as confirmed by streptavidin labeling, with minimal background outside the boundaries of axonal spheroids (**Figures 1E-H**). To further validate the spatial precision of proximity labeling, we employed Stimulated Emission Depletion (STED) super-resolution imaging, demonstrating the high precision of proximity labeling of axonal spheroids in AD human brains (**Figure S3**). Additionally, to provide evidence of the subcellular specificity of proximity labeling, we conducted parallel experiments utilizing the neuronal nuclear and perinuclear cytoplasm marker NeuN as the protein bait. (**Figure S2C-D**).

Furthermore, we optimized the protein lysis method, resulting in a significant improvement in protein extraction efficiency compared to previous studies [43]. This new protocol involved increased SDS concentration to 2% in basic Tris-HCl solution (pH = 8.0), which led to significantly increased protein extraction yield by effectively decrosslinking proteins in fixed postmortem tissue (**Figure S4A** and method section). Using this approach, we performed pulldown of biotinylated proteins from the lysed tissues and subsequently detected them through streptavidin-HRP western blotting. Our analysis revealed a diverse array of proteins, including the protein baits PLD3 or NeuN, along with axonal spheroid proteins RAGC and Cathepsin B (**Figures 1I and Figures S4B-D**). Thus, the refined proximity labeling method for postmortem brain samples provides a robust strategy for tagging and isolating proteins enriched in axonal spheroids, thereby facilitating comprehensive mass-spectrometry-based proteomic profiling.

### Proteomic analysis of plaque-associated axonal spheroids

To uncover the proteome of axonal spheroids, samples from individuals with AD and unaffected controls were processed for PLD3 proximity labeling protein biotinylation (**Figures 2A and S5**). Human frontal cortex postmortem samples were obtained from 39 individuals, sourced from the Yale Alzheimer’s Research Center (Yale ADRC, n = 8 AD and 2 control samples) and the Banner Sun Health Research Institute (Banner, n = 17 AD and 12 control samples), with detailed clinical and postmortem neuropathological data (**Figure S1**). All AD subjects exhibited high levels of amyloid plaques and other pathological hallmarks of AD, while control individuals had minimal or no amyloid plaque burden. For PAAS proximity labeling proteomics, we selected 6 AD individuals (3 females and 3 males), with the highest amyloid plaque burden (**Figure S1C**), along with 8 unaffected controls (3 females and 5 males). Additionally, 25 AD individuals (13 females and 12 males) and 7 unaffected controls (2 females and 5 males) were chosen for immunofluorescence validation of protein expression.

**Figure 2.**
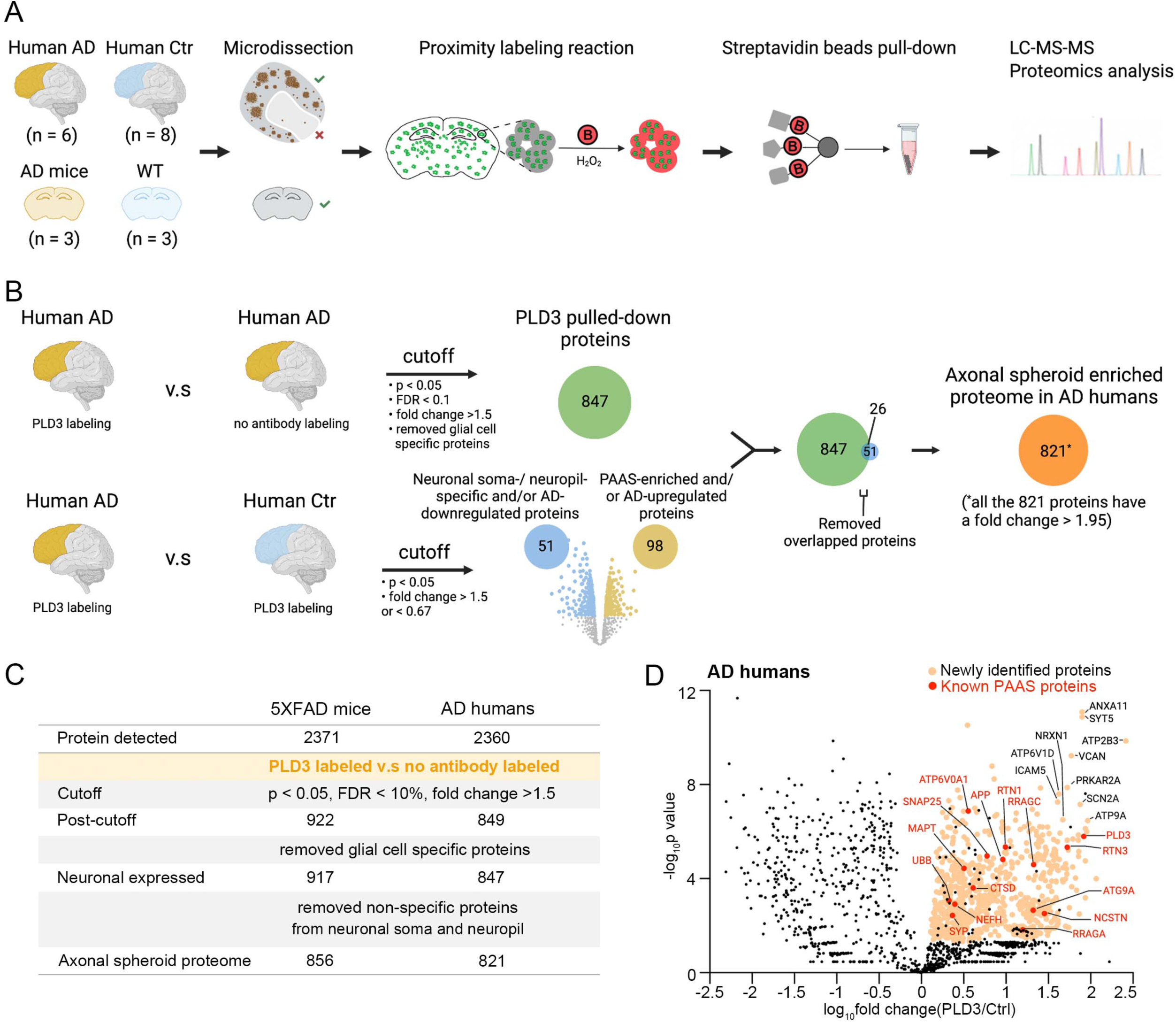
Proteomic analysis of plaque-associated axonal spheroids in AD humans and 5XFAD mice. **A.** Schematic showing the technical pipeline of PAAS proteomic analysis. The grey matter regions with high plaque load were microdissected out from human AD brain sections under guidance of a fluorescent stereomicroscope. Similarly, the grey matter regions were dissected from unaffected control brain sections. **B.** Statistical pipeline for revealing PAAS proteomes in humans (related to Figure 2C **and Figure S2E-G**). The same pipeline was applied to uncover PAAS proteomes in 5XFAD mice. **C.** Table showing the statistical cutoffs and summary of identified proteomic hits in AD humans and 5XFAD mice. The final PAAS proteome in humans contains 821 proteins (all of them have a fold change > 1.95), and the final PAAS proteome in mice contains 856 proteins (all of them have a fold change > 1.66). **D.** Volcano plot show proteins that passed the statistical cutoffs (orange dots) in AD humans. The top 10 proteomic hits with the lowest p-value and highest fold changes are indicated by their gene names in black. Selected known PAAS proteins are labeled as red dots, with their gene names in red. The black dots among the yellow ones represent proteins filtered out by the statistical pipeline (Figure 2B). (See **Table S1** for full list of proteomic hits). See also **Figures S5-S13**.

Through proteomic analysis of these human samples, we identified a total of 2,360 proteins (**Figures 2B** and **2C**). The process of identifying the PAAS proteomes involved three steps (**Figures 2B** and **2C**). First, to eliminate non-specific protein binders to beads, we compared PLD3 antibody-labeled samples to no-antibody samples, using cutoffs for p-value of < 0.05, false discovery rate (FDR) < 0.1, and fold change > 1.5. To ensure the stringency of the proteomic data, we compared normalized total precursor intensity (NTPI) and normalized total spectra count (NTSC) methods for quantification (**Figure S6**). We found that 870 proteins (NTSC) and 965 proteins (NTPI) remained after applying these statistical cutoffs, and 849 proteins were shared between the two methods. Next, we used the 849 shared proteins for downstream analysis. Second, we searched for glial cell-specific proteins and found only two (see method section), which were then excluded from the dataset, leaving 847 proteins. Finally, we aimed to exclude proteins specific to neuronal soma and neuropil, by comparing the proteomes of PLD3 antibody-labeled AD samples to unaffected controls, applying cutoffs for p-value of < 0.05 and fold change > 1.5 or < 0.67. Given that the PLD3-labeling in unaffected controls is derived from neuronal soma and neuropil given the lack of plaques, the proteins with fold change > 1.5 (98 proteins) represented those enriched in PAAS and/or broadly increased in AD, while those with a fold change < 0.67 (51 proteins) represented those specific to neuronal soma and neuropil, and/or proteins decreased in AD (**Figure 2B**). To increase stringency, we removed proteins with a fold change < 0.67 from the PAAS proteomic dataset (**Figure 2B**). As a result, 821 proteins remained, representing the PAAS proteomes in AD humans, all of which exhibited a fold change enrichment greater than 1.95 (**Figures 2B, 2C, Figures S7-S8 and Table S1**).

For comparative analysis, parallel experiments were conducted with 15-month-old 5XFAD mice (**Figure S9**). Using a similar proteomic strategy, we identified 856 PAAS proteomic hits in mice (**Figures 2A-C, Figure S9A** and **Table S1**, also see method section). All 856 proteins had a fold change greater than 1.66 (**Table S1**), among which 476 proteins were shared between AD humans and 5XFAD mice (**Figure S9B**). The proteomic analyses in both humans and mice uncovered hundreds of proteins that were not previously known to be expressed in PAAS, as well as those previously reported (**Figure 2D, Figure S9A and S10, and Table S2**).

Various controls were implemented to ensure specificity. This included additional proteomics using Lamp1 as a bait in 5XFAD mice, which detected 510 overlapping protein hits with PLD3 However, Lamp1 proximity labeling also detected many more proteins derived from glial cells (**Figure S11**), consistent with its expression in these cells. To further confirm the robustness of proteomic hits, we used anti-biotin beads for pulldown of biotinylated peptides, showing results that were highly consistent with those from the streptavidin beads pulldown (**Figure S12**). Additionally, to further examine the subcellular specificity of the antibody-based proximity labeling proteomic approach, we included NeuN, a neuronal nuclei and perinuclear cytoplasm marker [51], as a control protein bait. In contrast to the PLD3-labeled PAAS proteome, the NeuN-labeled proteome showed distinct specificity to nuclei and neuronal soma (**Figure S13 and Table S1**).

### Protein turnover and cytoskeletal dynamics are key signatures in axonal spheroids

To gain insights into the molecular mechanisms associated with axonal spheroid pathology, we conducted Gene Ontology (GO) annotation of biological process, molecular function and cellular component using the human PAAS proteomics dataset of 821 proteins, followed by pathway enrichment analysis. The results showed that the proteomic hits were mainly associated with axon, synapse, cytoskeleton, lysosome and proteosome complex (Figure **3A** **and Table S4**). These findings likely reflect the axonal origin of PAAS and the massive accumulation of endolysosomal organelles within them, as demonstrated by immunofluorescence imaging and electron microscopy (**Figures 1B-C and Movie S1**) [4, 14]. Since many synapse-related proteins are involved in vesicle fusion, such as the SNARE complex, these proteins are also actively involved in endolysosomal function. Given the fact that PAAS structures do not contain pre- and post-synaptic features (**Movie S1**), we speculate that the synapse-related signatures from the GO annotation imply the presence of vesicle fusion processes within the endolysosomal pathway.

**Figure 3.**
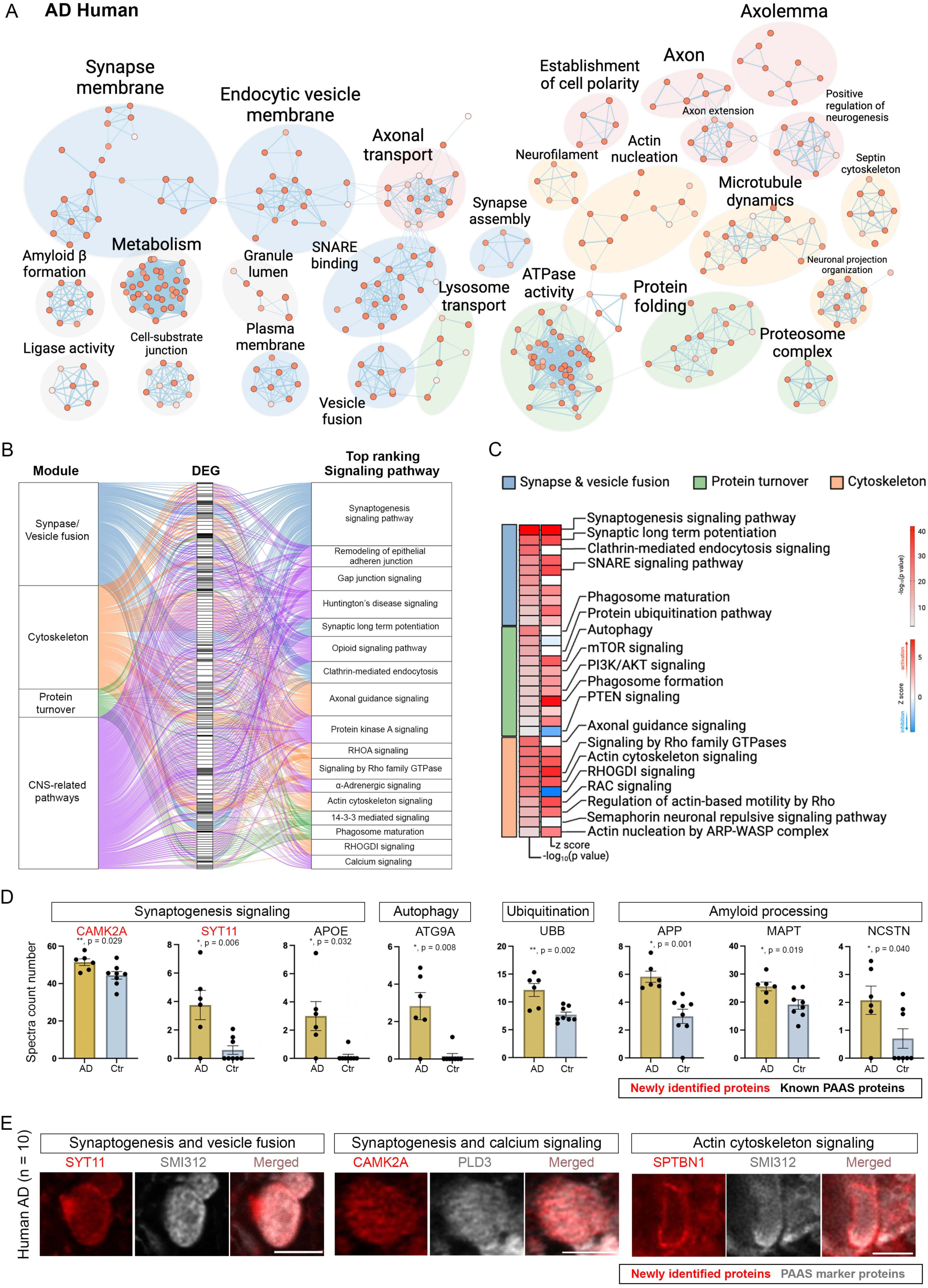
Pathway enrichment and signaling pathway analyses reveal that proteins related to protein turnover and the cytoskeleton are key components of PAAS. **A.** Pathway enrichment analysis of PAAS proteome in AD human brains. The Enrichment Map represents a network of pathways where edges connect pathways with many shared genes. Node color reflects the FDR of each pathway. The theme labels were curated based on the main pathways of each subnetwork. Subnetworks with a minimum of four pathways connected by edges are shown. **B**. IPA pathway analysis of the PAAS proteome in AD humans. Top ranking CNS-related signaling pathways are shown. The signaling pathways are summarized as four modules. The alluvium plot shows different color-coded modules connect to the differentially expressed genes (DEGs) and the DEGs connects to the pathways that they are involved. **C.** IPA pathways related to the three modules (synapse/vesicle fusion, protein turnover and cytoskeleton) with p-values less than 0.01 are listed. Heatmaps indicate either the -log10 (p-value) or the z score of each signaling pathway (pathways with a z score in red are predicted to be activated while blue ones are predicted to be inhibited). **D.** Bar chart shows representative proteomic hits from the signaling pathways in (**C**). The newly identified proteins are labeled in red, and the known PAAS proteins are labeled in black. n = 6 Human AD, and n = 8 unaffected human control brains were utilized. Error bars indicate SEM. **E.** Representative immunofluorescence confocal images of newly identified proteins (red) expressed in spheroids (grey) in AD postmortem brains. Scale bar = 5 μm. Zoom-out images are shown in **Figure S14**. Quantification was performed in n = 10 AD human brains. Protein expression quantifications can be found in **Table S2**. See also **Figures S9 and S14.**

Next, we performed signaling pathway analysis using the human PAAS proteome. We found that 10 out of the top 17 CNS-related pathways are involved in three main modules: synapse/vesicle fusion, protein turnover and cytoskeleton (**Figure 3B**), which is consistent with the findings from the GO analysis (**Figure 3A**). Specifically, among these pathways, we identified 5 pathways related to the cytoskeleton (e.g., axonal guidance), 3 related to synapse and vesicle fusion (e.g., synaptogenesis), and 2 related to protein turnover (e.g., phagosome maturation) (**Figure 3B** and **Table S5**). Additionally, beyond the top-ranked pathways, we also observed signaling pathways related to synapse and vesicle fusion, such as Clathrin-mediated endocytosis signaling, as well as pathways related to cytoskeleton growth and dynamics, such as actin cytoskeletal signaling and Rho family GTPase signaling (**Figure 3C**). Furthermore, pathways related to protein turnover, such as protein ubiquitination, autophagy and phagosome formation, were also identified. In addition, we noted the activation of the PI3K/AKT and mTOR signaling pathways, along with inhibition of the PTEN signaling (**Figure 3C**), all of which have been implicated in the regulation of protein turnover and axonal growth [52]. Importantly, subsets of proteomic hits from these identified pathways were observed to be increased in AD humans compared to controls (**Figure 3D**).

To confirm the expression of proteomic hits in PAAS, we selected various proteins from different pathways for validation using high-resolution immunofluorescence confocal microscopy. These included proteins associated with synaptogenesis, vesicle fusion, and calcium signaling (e.g., SYT11 and CAMK2A), and proteins involved in cytoskeleton dynamics (e.g., SPTBN1) (**Figure 3E, Figure S14 and Table S2**). A complete list of validated proteins in this study can be found in **Figure S10** and **Table S2**. For immunofluorescence validation, we used either SMI312 or PLD3 to reveal the structure of PAAS. SMI312 labels neurofilament and serves as a pan-axonal marker. Under normal conditions, SMI312 signals reveal the typical morphology of axons. However, in the context of AD, SMI312 is not only expressed along axons but also accumulates significantly in PAAS, indicating cytoskeletal abnormalities. SMI312 has been widely validated as a marker for PAAS due to its ability to highlight these structures in pathological conditions [15, 16]. Spheroids can be easily identified with this marker, making SMI312 very useful for colocalizations studies in conjunction with other antibodies. However, SMI312 is only expressed in a subset of spheroids [14]. Similarly, the newly validated proteins from our proteomic dataset demonstrate heterogeneity in the degree of expression. This heterogeneous protein expression pattern within spheroids suggests a potential mechanistic sequence of events at various stages during spheroid formation and growth.

Additionally, we conducted parallel PAAS proteomics analysis in 5XFAD mice. Similar to the human PAAS proteome, we observed the presence of GO terms related to axon, cytoskeleton, SNARE binding, as well as lysosome and endosome transport (**Figure S9C**). In addition, many signaling pathways related to synapse/vesicle fusion, protein turnover and cytoskeleton dynamics were captured in the PAAS proteomes from 5XFAD mice **(Figures S9D-E).**

### Lipid transport signaling is markedly upregulated in axonal spheroids

To investigate the aberrant signaling in PLD3-labeled AD brains compared to unaffected controls, we employed gene set enrichment analysis (GSEA) (**Figure 4A and Table S6**). This analysis revealed a significant upregulation of biological processes related to lipid transport (**Figures 4A** and **4B**). Among the top-ranked proteins associated with these processes were ATP8A1, C3, APOE, ATG9A, ATP8A2, TMEM30A, HEXB, and HDLBP (**Figure 4C**), all of which were identified in the PAAS proteome and increased in AD (**Figure 4D**, related to **Figure 2B, and Figure S10**). Conversely, GSEA analysis demonstrated the downregulation of biological processes related to ribosome, translation, and RNA metabolism in PLD3-labeled AD brains compared to unaffected controls (**Figure 4A**). Given that the PLD3-labeled signals in unaffected controls are derived from neuronal soma and neuropil, these downregulated signals reflect the protein functions associated with neuronal soma and neuropil identified in the unaffected control brains.

**Figure 4.**
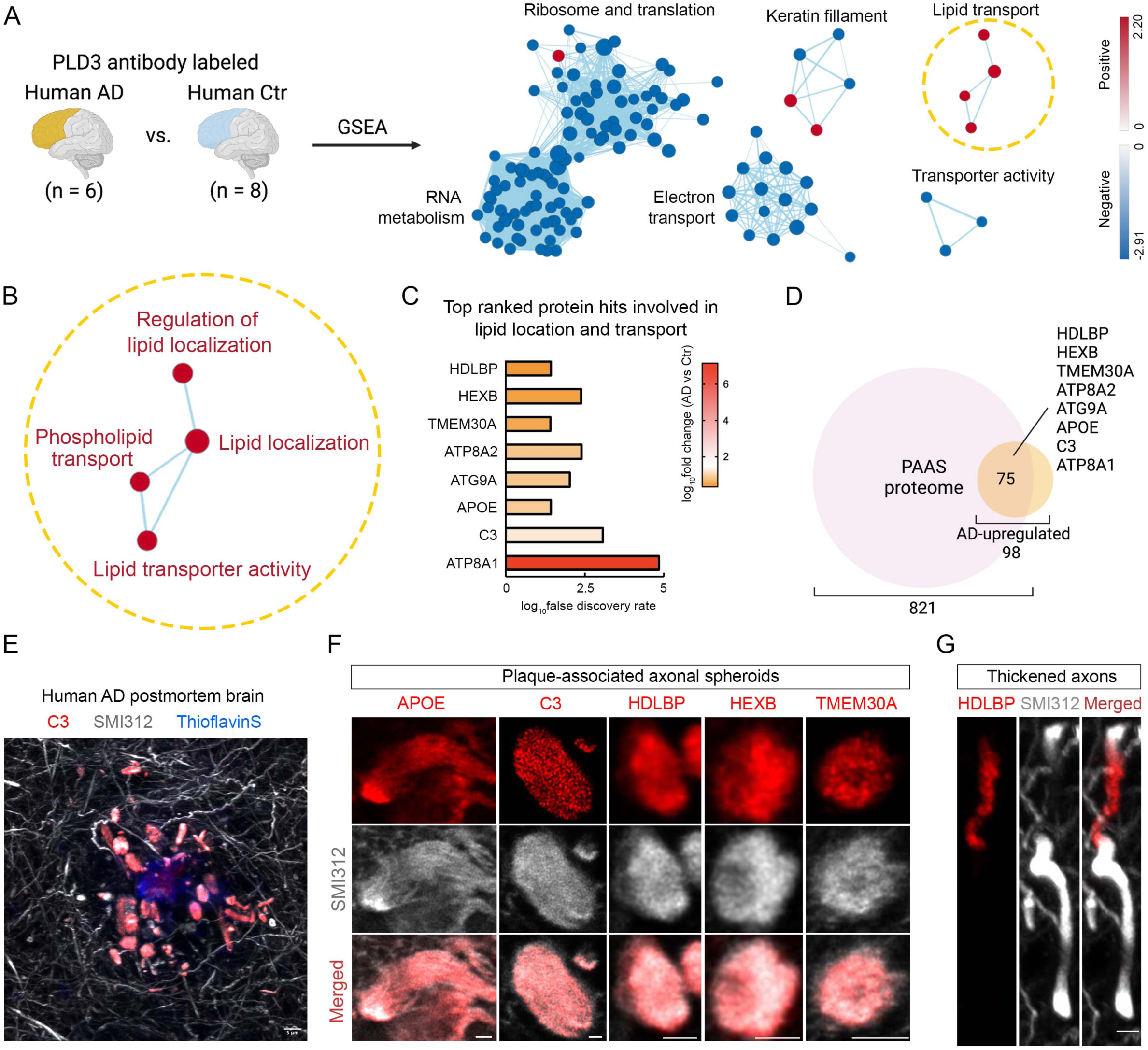
Proteins involved in lipid transport are upregulated in PAAS. **A.** Gene set enrichment analysis (GSEA) was performed to compare PLD3-labeled proteins between AD humans and unaffected controls. Pathway enrichment analysis was performed to cluster GSEA nodes. Each node represents a biological process or cellular component. The name of each cluster was curated based on the main GSEA biological processes and cellular components within each cluster. See also **Table S6**. **B.** Detailed information on the lipid transport cluster. The biological process or cellular component of each node is listed. **C.** The 8 top-ranked proteomic hits involved in the lipid transport cluster. The bar chart shows the fold change and FDR of these hits by comparing PLD3-labeled AD humans versus unaffected control humans. **D.** Venn diagram shows that the 8 top-ranked lipid transport-related proteins are shared between the human PAAS proteomes (821 proteins) and the AD-upregulated proteins (98 proteins). There are 75 proteins shared between these two datasets. **E-F.** Representative (**E**) zoomed-out and (**F**) zoomed-in immunofluorescence confocal images of the top-ranked lipid-related proteomic hits in AD human brain, including C3, APOE, HDLBP, HEXB and TMEM30A. Scale bar = 5 μm. Zoomed-out images of all the proteins are shown in **Figure S15**. Quantification was performed in n = 3 AD human brains. Protein expression quantifications can be found in **Table S2**. **G.** Representative immunofluorescence confocal images show the anti-colocalized distribution of HDLBP (red) and the pan-axonal marker SMI312 (grey) within thickened axons in the AD human postmortem brain (n = 3). Scale bar = 5 μm.

To validate the enrichment of the top-ranked lipid-related proteomic hits found in PAAS, including APOE, HDLBP, C3, HEXB, and TMEM30A, we performed immunofluorescence confocal imaging (**Figures 4E, 4F and Figure S15A**). Notably, APOE, which is the strongest genetic risk factor for AD and a lipid transporter [53], was among the top hits. We observed varying degrees of expression of these proteins in axonal spheroids in AD human brains (**Figures 4E, 4F, Figure S15A and Table S2**). Among them, Complement C3 (C3), APOE and high-density lipid binding protein (HDLBP) exhibited the highest expression in axonal spheroids and aberrant axons around amyloid plaques, but much less in distal axons away from plaques (**Figure 4E, Figure S15A and Table S2**). Additionally, these proteins were found to be expressed in cell bodies, neuropil, and amyloid plaque-related regions, consistent with their expected protein distribution patterns (**Figure S15A and Table S2**). Notably, HDLBP showed a peculiar pattern of segregation within a subset of thickened axon segments where the pan-axonal marker SMI312 was absent (**Figure 4G**), suggesting that lipid metabolism dysregulation may precede the enlargement of axonal spheroids. Regarding HEXB, known to be specifically expressed in microglia in mice [54], we found that in human brains it was expressed in PAAS and neuronal cell bodies (**Figure S15B**), consistent with human single cell transcriptional profiles [55].

### mTOR signaling is expressed in axonal spheroids and phosphorylated mTOR S2448 correlates with Alzheimer’s disease

The PAAS proteomics results drew our focus to the PI3K/AKT/mTOR axis, as it emerged as an activated signaling pathway within PAAS (**Figure 3C** and **Figure S9E**). This pathway is known to be a master regulator of mRNA translation, metabolism and protein turnover [35, 52]. In line with this, protein turnover, lipid metabolism, and axonal cytoskeleton dynamics were identified as prominent signatures in our proteomic analysis (**Figures 3, 4 and Figure S9D-E**) [33–35]. Therefore, we selected key proteins within the PI3K/AKT/mTOR axis for validation. Among them, mTOR, RAGA (RRAGA), RAGC (RRAGC), LAMTOR1 and AKT1 were detected in the PAAS proteomes in humans and/or mice, while PIK3R4, RHEB and RAPTOR were not detected (**Figure S10 and Tables S1 and S2**). Immunofluorescence staining revealed the presence of all selected proteins associated with the PI3K/AKT/mTOR axis in axonal spheroids in both postmortem AD human brains and 5XFAD mice (**Figure 5A**). The proteins that were detected in the PAAS proteomes showed moderate to high immunofluorescence signals in spheroids, while those that were not detected showed moderate to low expression (**Figure 5A and Table S2**). Notably, phosphorylated-mTOR-S2448, a marker of mTOR signaling activation, was found to be expressed in axonal spheroids in AD human brains, but not in unaffected controls (**Figures 5B-D**). This observation suggests that phosphorylated-mTOR-S2448 could potentially serve as a marker for disease progression. Overall, these findings underscore the involvement and activation of the PI3K/AKT/mTOR axis within axonal spheroids in AD.

**Figure 5.**
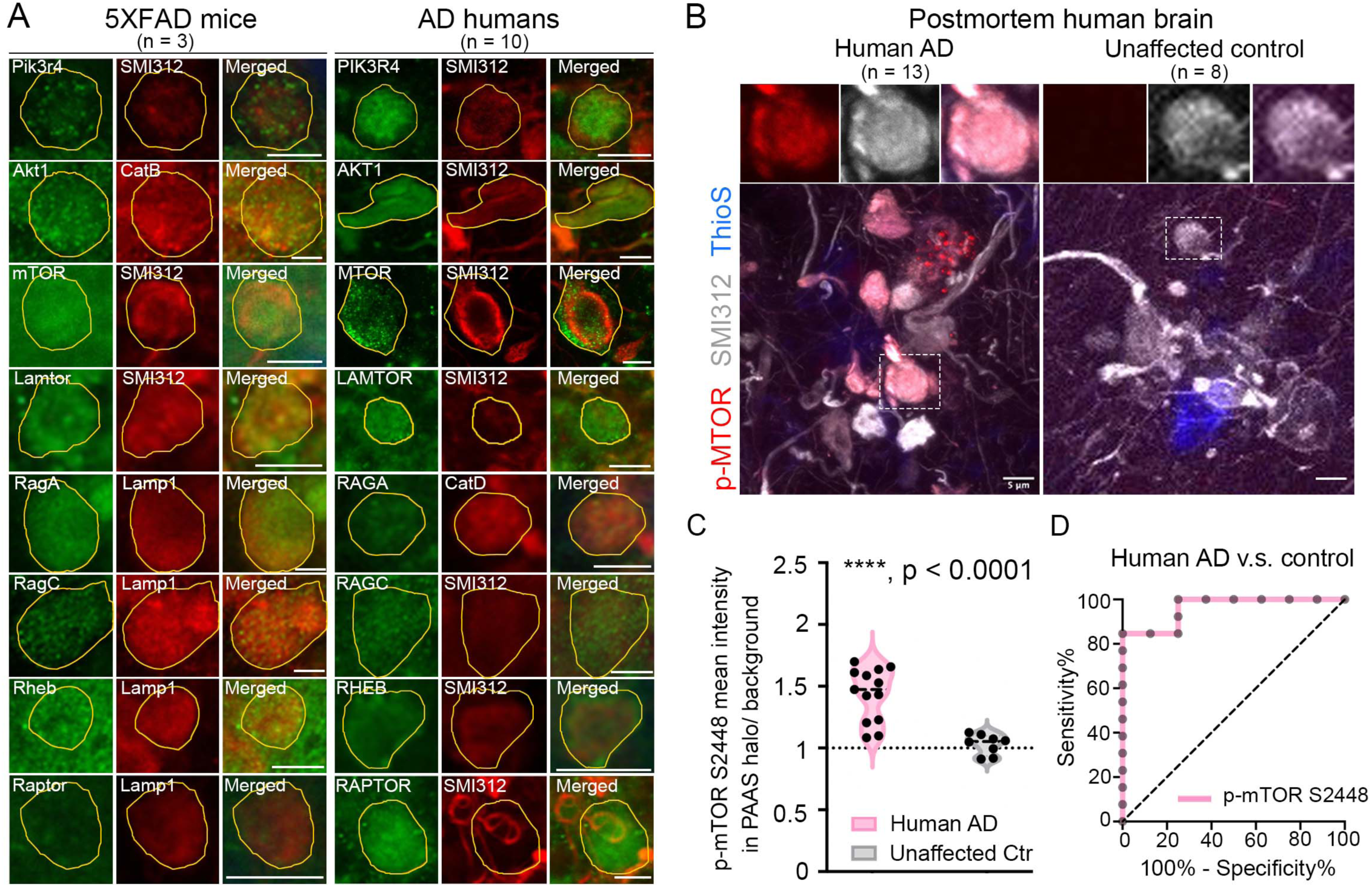
mTOR signaling is expressed in axonal spheroids and is associated with Alzheimer’s pathology. **A.** Immunofluorescence confocal imaging validation of selected proteomic hits and the related proteins in the PI3K/AKT/mTOR axis, revealing that signaling molecules of the PI3K/AKT/mTOR axis are expressed in PAAS in both AD humans and 5XFAD mice. PAAS were labeled using traditional markers including neurofilament SMI312, Cathepsin B (CatB), Cathepsin D (CatD), or Lamp1. PAAS are outlined in yellow. Scale bar = 5 μm. Protein expression quantification results can be found in **Table S2**. **B.** Phosphorylated-mTOR-S2448 (red) is highly enriched within PAAS (grey, SMI312) around amyloid plaques (blue, ThioflavinS) in severe AD. Scale bar = 5 μm. **C.** Quantification of the mean fluorescence intensity levels of p-mTOR-S2448 within axonal spheroid halos normalized to background fluorescence, comparing human AD (n = 13 brains) versus unaffected controls (n = 8 brains). Unpaired t-test (Mann Whitney test), two-tailed, ****, p < 0.0001. The Black dashed line indicates the median. **D.** A Receiver Operating Characteristic (ROC) curve shows that the p-mTOR-S2448 level in PAAS significantly separates AD brains from those of unaffected controls. The area under the ROC curve = 0.962, standard error = 0.038, 95% confidence interval = 0.888 to 1.000, p-value = 0.0005.

### Human iPSC modeling replicates cellular and functional abnormalities of axonal spheroids

To establish a comprehensive strategy for investigating selected proteomic hits and their potential roles in spheroid formation, we utilized long-term human iPSC-derived neuron [56] and astrocyte co-cultures (**Figures 6A-B and S16A-D**). A similar approach was recently shown to replicate the formation of axonal spheroids upon administration of exogenous aggregated β-amyloid 1-42 [57]. We simplified the human neuron induction protocol by using the *NGN2*-induced glutamatergic neuron approach [56] (see method section), making it more accessible for most laboratories. Additionally, we increased the overall neuronal density in the cultures to better mimic the axonal spheroids observed around amyloid plaques in the human brain. Our optimized model successfully generated Thioflavin S-positive amyloid deposits surrounded by many axonal spheroids (**Figures 6B, Figures S16E-F, S17A and S18B-C**). These spheroids accumulated lysosomes and autophagosomes (**Figures 6B-C**), and expressed phosphorylated Tau S235, S396 and S404 (**Figure S16G**), closely resembling axonal spheroids in humans [4, 58].

**Figure 6.**
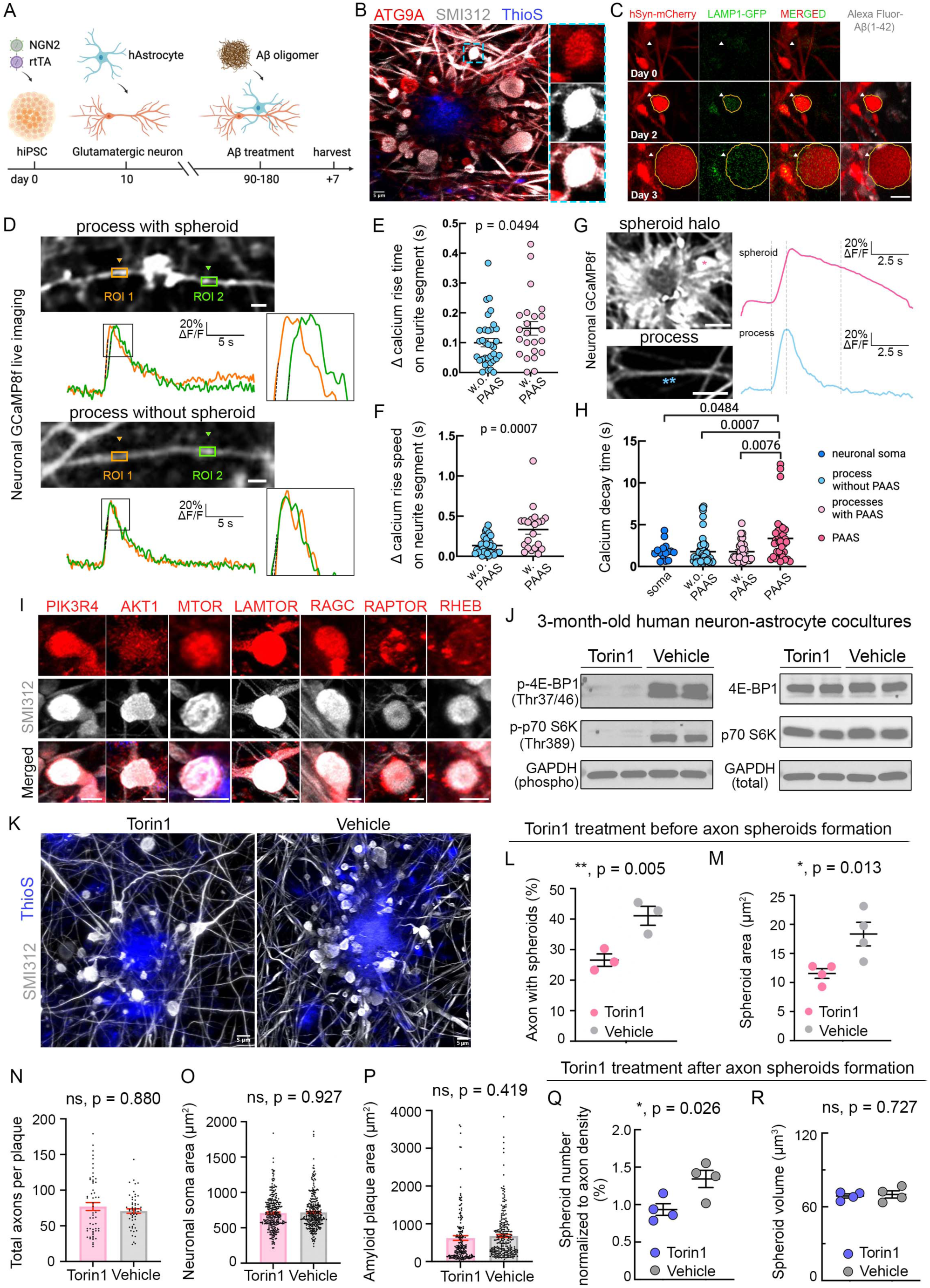
A human iPSC-derived AD model demonstrates that mTOR signaling inhibition reduces PAAS pathology. **A.** The workflow shows the generation of a human iPSC-derived neuron and astrocyte co-culture AD model to recapitulate axonal spheroid pathology around amyloid plaques. Human iPSCs (hiPSCs) are plated at day 0 to generate *NGN2*-induced glutamatergic human neurons. Human primary astrocytes (hAstrocytes) are seeded at day 10, followed by 3 to 6 months of co-culture. Oligomerized Aβ 1-42 peptides are administered for 7 days to the neuron-glia co-culture. **B.** Confocal deconvolved image shows the robust formation of axonal spheroids (grey, SMI312) in human neurons around amyloid deposits (blue, ThioflavinS). These spheroids expressed the axonal cytoskeleton marker SMI312 (grey) and autophagosome marker ATG9A (red), resembling PAAS in postmortem AD brains. **C.** Time-lapse imaging of human neurons shows the formation of a spheroid (arrowhead) from a neurite (red, AAV9-hSyn-mCherry labeled). Lysosomes (green, AAV2-CMV-LAMP1-GFP labeled) accumulate within spheroids. Spheroids (arrowhead) near Aβ deposits (grey) enlarged over time. **D-H.** Neuronal GCaMP8f imaging in human iPSC AD model. **(D)** Representative images showing CAMKII-GCaMP8f labeled neuronal processes with (upper panel) or without (lower panel) axonal spheroids. Representative traces showing calcium dynamics recorded from neuronal processes segments in the human iPSC AD model. The y axis indicates the normalized ΔF/F of calcium transients. The dotted black lines indicates the calcium rise slope. **(E)** Quantification of calcium rise time and **(F)** calcium rise speed. Each dot represents a neuronal process from three independent experiments. Unpaired two-tailed Mann Whitney t test was performed. **(G)** Representative images showing CAMKII-GCaMP8f signal from axonal spheroid halo or neuronal processes. Representative traces showing that calcium decay time is much slower in spheroids (pink asterisk) than in neuronal processes (blue asterisks). The y axis indicates the normalized ΔF/F of calcium transients. **(H)** Quantification showing calcium decay time in neuronal soma (blue), neuronal processes without spheroids (light blue), neuronal processes with spheroids (light pink) and spheroids (pink). Each dot represents a neuronal process from three independent experiments. One-way ANOVA was performed to compare groups. **I.** mTOR signaling molecules (red) are expressed in hiPSC-derived axonal spheroids (grey, SMI312). Large-field-of-view images are shown in **Figure S17**. **J.** Western blot showing that Torin1 treatment markedly inhibited mTOR signaling, as revealed by the significant reduction of mTOR downstream effectors phosphorylated 4E-BP1 and phosphorylated p70 S6K, while their total protein level remained unchanged. **K-R.** Torin1 treatment reduced axonal spheroid pathology in iPSC-derived human neurons. **(K)** Immunofluorescence confocal deconvolved images show that Torin1 reduced axonal spheroids (grey, SMI312) around Aβ deposits (blue, ThioflavinS). **(L-P)** Torin1 treatment before amyloid plaque and spheroid formation. **(L)** Quantification of the percentage of axons with spheroids relative to the total number of axons around Aβ deposits. Torin group and vehicle groups (n = 3 independent experiments for each group). Paired t-test, two-tailed, **, p = 0.005. **(L)** Quantitative analysis of axonal spheroid size. Paired t-test, two-tailed, *, p = 0.013. Torin and vehicle groups (n = 4 independent experiments for each group). (**J-K**) Each dot represents an experiment in which 20 to 30 ROIs around plaques were quantified. **(N)** Quantification of axon number around an amyloid plaque in each ROI (represented by black dots). ROIs were quantified for Torin1 and vehicle treatment groups, n = 56 and n = 55, respectively. Unpaired t-test, two-tailed, p = 0.880. **(O)** Quantification of neuronal soma size (related to **Figure S15I**). Each dot represents an individual neuronal soma. Torin1 treatment group n = 298, vehicle group n = 316. Unpaired t-test, two-tailed, p = 0.927. **(P)** Quantification of amyloid plaque size. Each dot represents an amyloid plaque. Torin1 treatment group n = 201, vehicle group n = 253. Unpaired t-test, two-tailed, p = 0.419. **(Q-R)** Torin treatment after amyloid plaque and spheroid formation, related to **Figure S18**. Quantification of (Q) spheroid number normalized to axon density and (R) spheroid size. Unpaired Mann Whitney t-test, two-tailed was performed. Torin and vehicle groups (n = 4 independent experiments for each group). **(A-R**) All experiments were validated independently at least three times using cultures derived from two independent hiPSC lines. Scale bar = 5 μm, except scale bar = 5 μm in panel G. See also **Figures S16-S18**.

This culture system enabled us to perform longitudinal structural and functional imaging, providing insights into the dynamics and consequences of spheroid pathology. By utilizing reporter AAV-viruses to label neurons and lysosomes, we were able to track the formation of spheroids following administration of Aβ 1-42. By visualizing individual axons through confocal microscopy, we observed the gradual formation of spheroids and lysosomal accumulation as early as day 1, with spheroids increasing in size over a 7-day observation period after treatment (**Figures 6C**).

We also investigated the functional repercussions of spheroid formation in this human iPSC-derived AD model. Employing calcium imaging with the reporter GCaMP8f, we measured the Ca2+ rise time in axonal segments on both sides of axonal spheroids following electrical stimulation. We observed a significant difference in the slope of the calcium rise time in process segments with spheroids compared to those without (**Figures 6D-F**). This is consistent with impaired action potential conduction across the spheroids in human neurons, similar to our earlier in vivo findings in 5XFAD mice [4]. Additionally, we noted a very prolonged calcium decay time in process segments with spheroids relative to normal processes and somata, indicating disrupted calcium homeostasis within spheroids (**Figures 6G-H**).

### mTOR signaling inhibition reduces spheroid pathology in human neurons

We further explored the role of PI3K/AKT/mTOR signaling in the development and expansion of axonal spheroids using the human iPSC model. Consistent with our observations in postmortem human brains (**Figure 5**), the human iPSC-derived AD model exhibited the expression of proteins associated with the PI3K/AKT/mTOR signaling pathway within axonal spheroids (**Figure 6I and Figure S17**).

To assess the impact of mTOR inhibition on axonal spheroid pathology, we utilized Torin1, a compound that inhibits both mTORC1 and mTORC2 [59, 60]. Treating 3-month-old iPSC-derived human neurons and astrocyte co-cultures with Torin1 for 7 days led to a significant suppression of mTOR signaling, evidenced by reduced levels of mTOR downstream effectors p-p70 S6K (Thr 389) and p-4E BP1 (Thr 37/46), without affecting their total protein levels (**Figure 6J**). To assess the effects of Torin1 on the formation and reversibility of axonal spheroids, we treated cultures with Torin1 either before or after β-amyloid exposure and spheroid formation. Pre-treatment with Torin1 prior to Aβ exposure resulted in a considerable decrease in both the number and size of individual spheroids (**Figures 6K-P**), while treatment with Torin1 after Aβ exposure reduced the number of spheroids but did not alter their size (**Figures 6Q-R and Figures S18A-C**). Importantly, none of these outcomes were attributable to axonal loss (**Figure 6N and Figure S18D**), changes in neuronal density or amyloid plaque size (**Figures 6O, 6P, Figures S16H-I and S18E**). Thus, our findings indicate that mTOR signaling plays an important role in amyloid-induced spheroid formation and suggest that targeting mTOR could be a promising approach for both preventing and reversing, as well as initiation and maintenance of spheroid pathology.

#### Amelioration of spheroid pathology in 5XFAD mice through genetic manipulation of mTOR

To investigate the effect of mTOR signaling on axonal spheroids in vivo, we employed a viral-mediated Cre/lox-based *Mtor* knockout approach in 5XFAD mice (**Figures 7A-C and S19A-D**). Using heterozygous *Mtor*-floxed-5XFAD mice [61] and infecting them with AAV9-hsyn-cre-2a-tdTomato, we achieved partial loss of mTOR expression in neurons. Sparse neuronal infection allowed us to clearly visualize individual spheroids (**Figures 7D-E**), revealing a significant decrease in axonal spheroid size following heterozygous *Mtor* knockout (**Figures 7F-G**).

**Figure 7.**
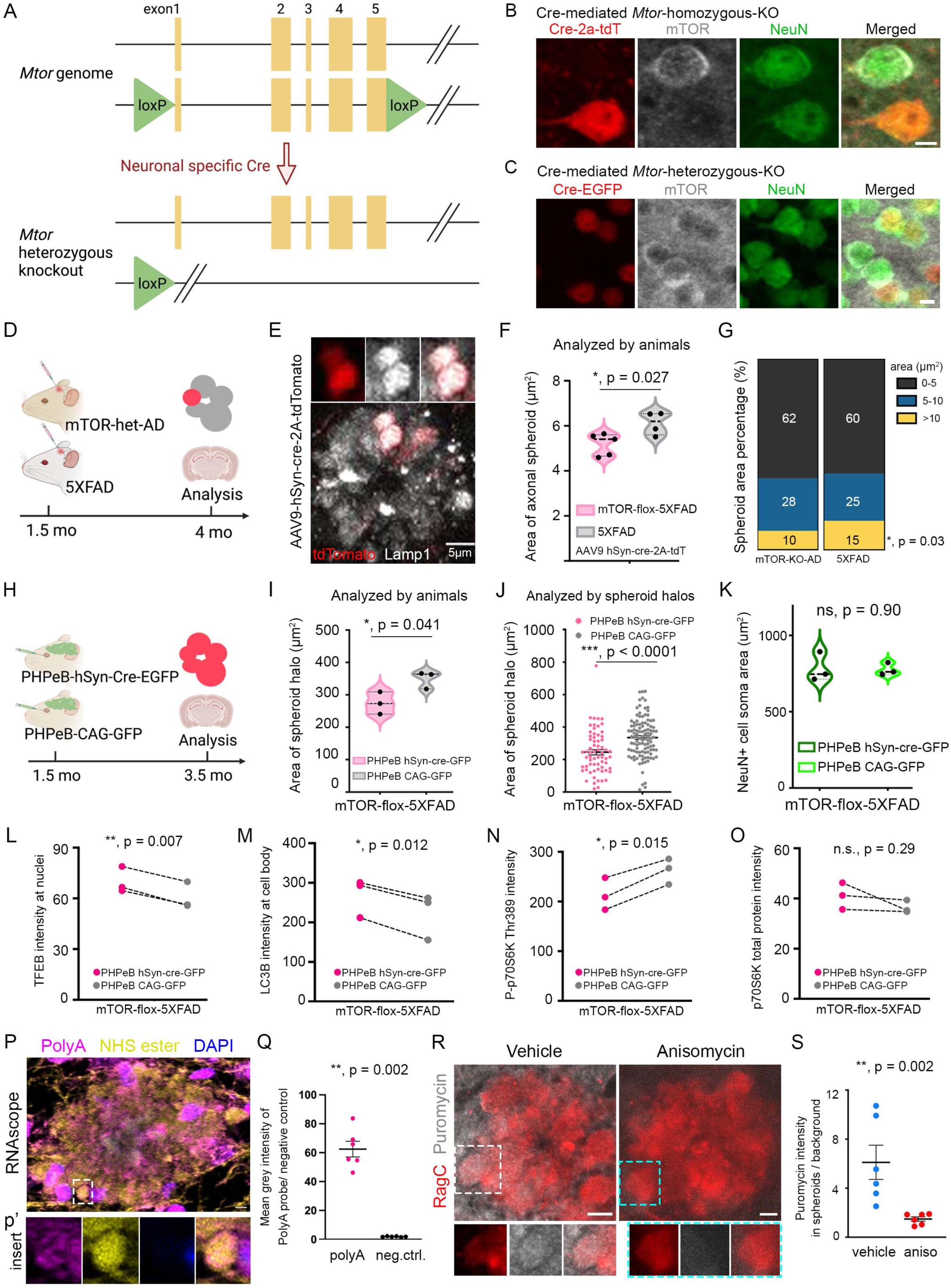
mTOR signaling inhibition reduces axonal spheroid pathology *in vivo*. **A.** Schematic of neuronal-specific conditional knockout of *Mtor* in the heterozygous floxed mouse. **B-C.** Immunofluorescence confocal images showing neuronal specific Cre-mediated *Mtor* homozygous knockout **(B)**, and heterozygous knockout **(C)** in *Mtor*-floxed mice using AAV9-hSyn-Cre-2A-tdTomato, or AAV PHPeB-hSyn-Cre-EGFP, respectively. (**B**) mTOR expression (grey) was absent in Cre-expressing neurons (red), compared to the adjacent neuron (green, NeuN) without Cre expression. (**C**) mTOR expression (red) was reduced in Cre-expressing neurons (green), compared to other neurons (grey, NeuN) without Cre expression. Scale bar 5 μm. **D.** Experimental design of virally mediated genetic manipulation to study the impact of *Mtor* heterozygous knockout on individual axonal spheroids in heterozygous *Mtor*-floxed-5XFAD mice**. E.** Immunofluorescence confocal images showing AAV9-hSyn-Cre-2A-tdTomato sparsely labeling individual axonal spheroids (red) within a spheroid halo (Lamp, grey). **F.** Quantification showing that AAV9-mediated *Mtor* heterozygous knockout in heterozygous *Mtor*-flox-5XFAD mice significantly reduced the size of individual axonal spheroids. Each dot represents an animal (n = 5 in each group). Unpaired t-test, two-tailed, *, p = 0.027. **G.** Using the same results in (**F**), quantification analysis shows that *Mtor* heterozygous knockout significantly reduced the percentage of larger axonal spheroids (area greater than 10 μm2). Unpaired t-test, two-tailed, *, p = 0.03. **H.** Experimental design of virally mediated genetic manipulation to study the impact of *Mtor* heterozygous knockout on the size of the axonal spheroid halo in heterozygous *Mtor*-floxed-5XFAD mice. **I.** Quantification showing that AAV.PHPeB-mediated *Mtor* heterozygous knockout in *Mtor*-flox-5XFAD mice significantly reduced the size of axonal spheroid halos. Each dot represents an animal (n = 3 in each group). Unpaired t-test, two-tailed, *, p = 0.041. **J.** Using the same results as in (**I**) and quantified by the axonal spheroid halo. Each dot represents an axonal spheroid halo. Knockout group n = 66 and control group n = 109 halos. Unpaired t-test, two-tailed, ****, p < 0.0001. **K.** AAV PHPeB-hSyn-Cre-EGFP mediated *Mtor* heterozygous knockout did not alter neuronal soma size. Each dot represents an animal (n = 3 in each group). Unpaired t-test, two-tailed, p = 0.90. **L-O.** Investigation of mTOR heterozygous knockout downstream signaling effectors in mTOR-floxed-5XFAD mice, comparing mice injected with PHPeB-hSyn-cre-GFP viruses, with those injected with control viruses PHPeB-CAG-GFP. Immunofluorescence intensity of (**L**) TFEB was measured in neuronal nuclei, and (**M**) LC3B, (**N**) P-p70S6K Thr389, (**O**) p70S6K were measured in neuronal soma in an automated fashion. The littermates and sex were paired as indicated by the paired points in (L-O), paired T-test was performed. Each dot represents an animal (n = 3 in each group). **P.** RNAscope in 5XFAD mice cortices showing that mRNA species (PolyA probe labeled, magenta) are expressed in axonal spheroids (NHS ester labeled, yellow and DAPI negative). NHS ester (yellow) labels the axonal spheroid halo, as well as the extracellular amyloid plaques. Nuclei are labeled with DAPI (blue). Scale bar = 5 µm. **Q.** Quantification showing fluorescence intensity of PolyA probe compared to that of negative control probe in axonal spheroids in 5XFAD mice cortices. Each dot represents a large field of view (FOV), two FOVs are shown in each animal (n = 3 in each group). Unpaired t-test, two-tailed, p = 0.002. **R.** Puromycylation in 5XFAD mice cortices showing nascent peptides (puromycin labeled, grey) in axonal spheroids (RagC labeled, red). Puromycin labeling was significantly reduced with the treatment of translation inhibitor anisomycin. Scale bar = 5 µm. **S.** Quantification of puromycin fluorescence intensity in axonal spheroids, comparing vehicle versus anisomycin control. Each dot represents a large field of view (FOV), two FOVs are shown in each animal (n = 3 in each group). Unpaired t-test, two-tailed, p = 0.002. See also **Figures S19-S21**.

To assess the overall impact on spheroid size and number, we measured the axonal spheroid halo size around individual amyloid plaques. For this, we infected *Mtor-*floxed-5XFAD mice with AAV-PHP.eB-hSyn-Cre-GFP virus, enabling dense neuronal infection, thereby inducing widespread *Mtor* heterozygous knockout in neurons (**Figures 7H and Figures S19B, D and E**). This manipulation significantly reduced the axonal spheroid halo size around plaques without affecting amyloid plaque size (**Figures 7I-J and Figures S19E-H**). Importantly, despite mTOR’s known role in cell growth and maturation [35, 59], the heterozygous *Mtor* knockout did not alter the size of neuronal cell bodies (**Figure 7K and Figure S19F**). These in vivo findings in mice, paralleling those in vitro with human iPSC-derived neurons, highlight the potential of PI3K/AKT/mTOR signaling as a therapeutic target for mitigating axonal spheroid pathology in AD.

Signaling pathways associated with lysosome biogenesis and autophagy are likely to play important roles in the accumulation of aberrant endolysosomes in spheroids. In addition, local mRNA translation has been shown to be important for axonal outgrowth, which could play a role in spheroid formation [62, 63]. Considering the known effects of mTOR on lysosome biogenesis, autophagy and local mRNA translation [35, 52], we explored related downstream molecules. Using AAV-PHPeB-hSyn-Cre to induce extensive neuronal infection in mTOR-heterozygous-floxed-5XFAD mice, we achieved widespread mTOR heterozygous knockout specifically in neurons (**Figures 7A-C and 7H**). Immunofluorescence confocal imaging was used to assess the expression levels of TFEB (lysosomal biogenesis transcription factor), p-p70S6K (regulator of mRNA local synthesis), and LC3B (a marker of autophagy) in neuronal somata, with automated quantitative analysis comparing expression between mTOR heterozygous knockout mice and controls (**Figures 7L-O**). Results indicated increased expression of TFEB and LC3B (**Figures 7L-M**), suggesting enhanced lysosomal biogenesis and autophagy. Additionally, the levels of p-p70S6K were decreased (**Figures 7N-O**), which may be associated with reduction in mRNA local translation [52].

We further examined whether mRNA local translation occurs at axonal spheroids. Initially, we probed the presence of mRNA species in axonal spheroids using RNAscope in 5XFAD mice (**Figure 7P and Figure S20A**). For this we used a probe against mRNA polyA tails and compared labeling patterns to a scrambled control probe. We found that the polyA but not the control probe signal was present in axonal spheroids, indicating the localization of mRNA in these structures (**Figure 7Q and Figures S20B-D**) (see methods for spheroid detection during RNAscope). Next, we examined the occurrence of mRNA local translation at axonal spheroids by performing an *in vivo* puromycylation assay in 5XFAD mice (**Figure 7R and Figure S21A**). We detected nascent proteins labeled by puromycin within axonal spheroids, and treatment with the protein translation inhibitor anisomycin reduced the amount of labeling (**Figure 7S and Figure S21A-C**), indicating that local mRNA translation may occur within axonal spheroids. To examine whether mTOR inhibition plays a role in regulating nascent protein production in spheroids, we performed both pharmacological and genetic manipulations *in vivo.* We first used Torin1 to inhibit mTOR in 5XFAD mice and observed that the puromycin signal remained unchanged. Additionally, we used mTOR-floxed mice and a PHPeB-hSyn-Cre virus to genetically knock out mTOR in neurons. However, neither the homozygous nor heterozygous mTOR knockouts had a significant effect on the puromycin signal (**Figures 7A-C, Figures S19A-D, and Figures S21D-F**). These results indicate that under our experimental conditions (**Figures S21D-F**), mTOR likely does not control local protein translation within axonal spheroids.

Altogether, these experiments indicate that the reduction in axonal spheroid size and number observed after mTOR inhibition is likely due to its enhancement of lysosomal biogenesis and autophagy, rather than modulation of local protein translation in spheroids.

## DISCUSSION

Plaque-associated axonal spheroids, also known as dystrophic neurites, have been recognized for over a century [10] and are one of the most prevalent neuropathological hallmarks in AD [11, 14, 64, 65]. However, the molecular composition and cellular biology underlying the progression of this pathology have not been systematically studied, primarily because of a lack of appropriate methodologies. In this study, we implemented a comprehensive approach to investigate the molecular and cellular mechanisms of PAAS formation. We developed a subcellular proximity labeling proteomics approach in postmortem human and mouse brains [43], which allowed us to uncover the molecular architecture of PAAS and other subcellular compartments. Through bioinformatics analysis and high-resolution confocal imaging validation, we identified hundreds of proteins and signaling pathways previously not known to be associated with PAAS, revealing lipid transport, protein turnover and cytoskeletal dynamics as signature biological processes associated with PAAS. Additionally, we found activation of the PI3K/AKT/mTOR signaling within PAAS, a pathway known to be a master regulator of these biological processes. Overall, the human PAAS proteomics provided a comprehensive view of the protein composition and potential signaling pathways operating within axonal spheroids (**Figure 8, Figure S10 and Table S2**). To investigate the functional implications of these findings on axonal spheroids pathogenesis in humans, we utilized an optimized human iPSC-derived AD model [57] that faithfully replicates amyloid plaque and PAAS formation. Both pharmacological and genetic inhibition of mTOR signaling in this model and in AD-like mice resulted in a significant reduction in PAAS pathology. This study, therefore, uncovers the molecular architecture and functional abnormalities of PAAS in human neurons and implicates the PI3K/AKT/mTOR axis as a key signaling pathway in PAAS formation and growth. Furthermore, it suggests a new potential strategy for mitigating axonal pathology, independent of amyloid removal.

**Figure 8.**
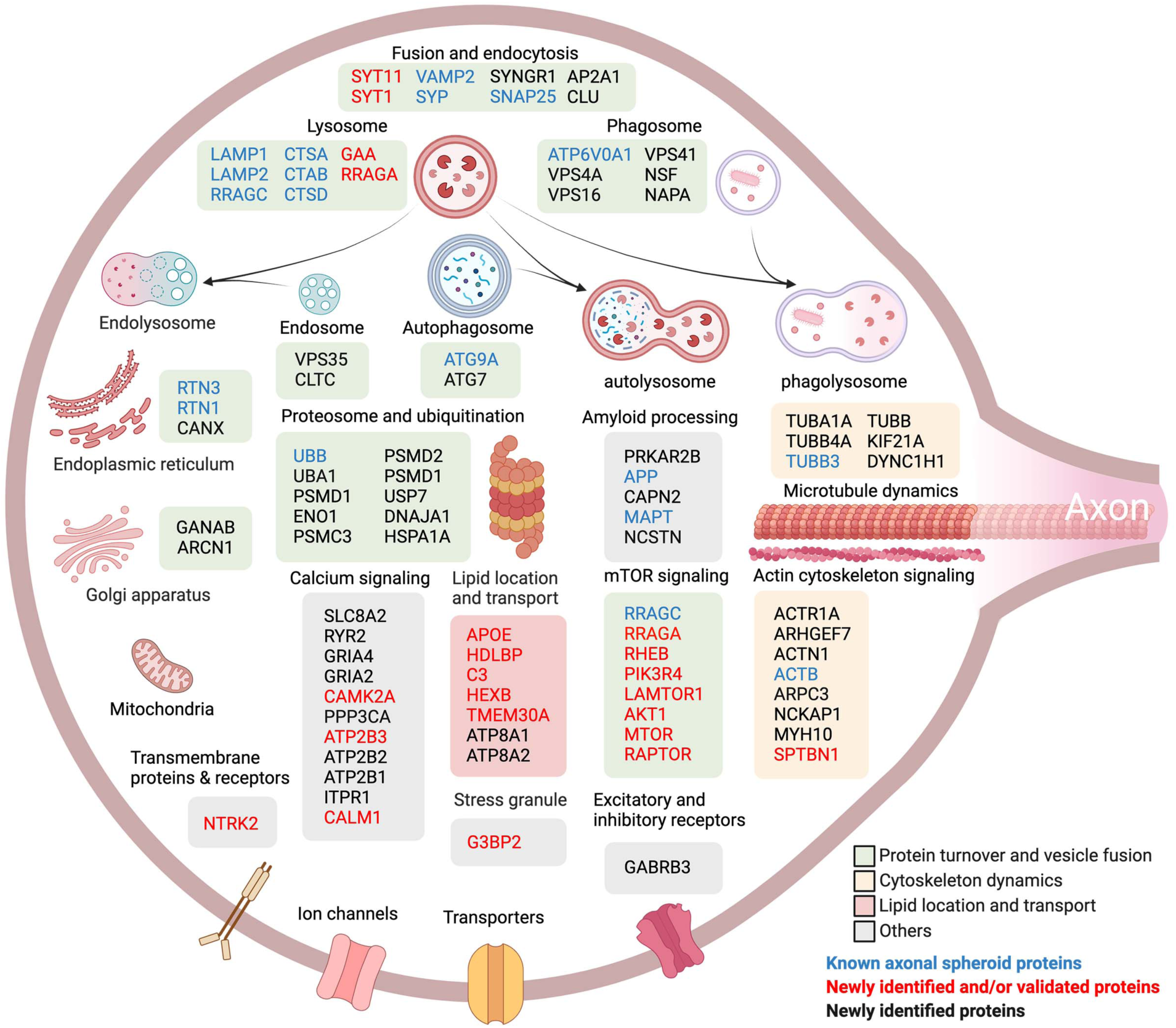
A schematic of the molecular architecture of plaque-associated axonal spheroids. Proximity-labeling proteomics reveals proteins preferentially involved in a variety of subcellular organelles, the ubiquitin-proteosome system and cytoskeleton. These proteins and their signaling pathways relate to biological functions including protein turnover and vesicle fusion (green box); cytoskeletal dynamics (yellow box); lipid localization and transport (red box) and others (grey box). Here we depict selected newly identified and validated proteins as well as those previously known to be enriched in PAAS (lysosomal proteins LAMP1 [4], Cathepsin B and D [14] , RAGC [14], PLD3 [47, 48]; autophagosome protein ATG9A [91] and endoplasmic reticulum protein RTN3 [91] and RTN1 [92]; cytoskeletal neurofilament protein [93], microtubule TUBB3 [20]; synaptic proteins Synaptophysin [5] and VAMP2 [14], APP [31], Tau (MAPT) [26] and Ubiquitin [71, 91]) (**Table S2**).

To uncover the PAAS proteome, we employed proximity labeling proteomics with subcellular resolution, a technique that overcomes the lack of cellular and subcellular specificity of conventional tissue proteomics, including micro-dissected amyloid plaques [66–68]. Using an antibody-based biotinylation method [43, 44] to tag proteins in axonal spheroids, without requiring overexpression of an exogenous peroxidase or biotin ligase [36, 37, 41], allowed us to obtain comparative proteomes from widely available postmortem tissues. We selected PLD3 as an antibody bait because this protein is highly enriched in PAAS of both humans and mice [47, 48] and is almost exclusively expressed in neurons [4, 47], thus minimizing the potential for contamination from other cell types. Indeed, comparison between PLD3- and Lamp1-labeled proteomes in 5XFAD mice showed that the PLD3-labeled proteome is much more specific for neuronal and axonal structures (**Figure S11**).

Utilizing our proteomic strategy, we discovered a range of proteins previously unknown to be enriched in PAAS, along with those already identified in these structures (**Figure 8**, **Figure S10, Tables S1 and S2**). Our proteomics analysis in AD humans highlighted the activation of three key signature biological processes that are likely involved in the formation and growth of PAAS (**Figures 3** and **4**). These key signatures include: 1) Proteolysis Dysfunction: This was indicated by the accumulation of proteins associated with endocytosis, phagosome, proteosome, ubiquitin-mediated proteolysis, and lysosome acidification (**Figures 3, 5, 8 and Table S2**). Prior studies have shown that protease-deficient lysosomes and autophagosomes within PAAS [14], and the enlargement of these vesicles, mediates the overall increase in spheroid size [4], highlighting the critical role of aberrant lysosomal function in spheroid formation and growth. 2) Cytoskeletal Dysregulation: we found that PAAS were enriched with activated signaling pathways, such as actin cytoskeletal signaling, RAC signaling and actin nucleation by the ARP-WASP complex. Conversely, pathways like RHOGDI signaling, which has been shown to regulate Rho family GTPase, were inhibited (**Figures 3C and S9E**) [69, 70]. This indicates that ongoing cytoskeletal reorganization and plasticity within PAAS could be important events in the initial formation and enlargement of these structures. Moreover, such cytoskeletal changes could impact both retrograde and anterograde axonal cargo transport, leading to further accumulation of endolysosomal vesicles [18, 28, 71], and the subsequent expansion of spheroids. 3) Lipid Transport and Metabolism: our results indicate that lipid transport is highly activated in PAAS (**Figures 4A-D**), and proteins involved in lipid transport and metabolisms, such as APOE, HDLBP and C3, are highly expressed within axonal spheroids and aberrant axons around amyloid plaques (**Figures 4B-G and Figure S15A**). Among these proteins, APOE is the most significant risk gene in AD and is a lipid transporter [53]. TMEM30A, ATP8A1 and ATP8A2 (**Figures 4C** and **4E**) compose the P4-ATPase complex, which controls the asymmetric membrane lipid distribution, maintains membrane stability and regulates vesicle-mediated protein transport [72]. The enrichment of these proteins likely relates to the massive accumulation of endolysosomal vesicles within PAAS, which must require active local lipid transport, synthesis and metabolism. This is in line with previous findings that lipids participate in mediating axonal lysosome delivery and spheroid formation [21]. Interestingly, complement C3, a crucial protein for activating the complement system [73] that is also implicated in lipid metabolism [74, 75], exhibited the highest expression in axonal spheroids and aberrant axons (**Figures 4E, 4F and Figure S15A**). This aligns with prior research indicating the expression of complement proteins in dystrophic neurites around compact amyloid plaques [76–78], and could suggest a connection between complement pathway activation in the formation of axonal spheroids, potentially independent of neuroimmune interactions. Importantly, based on our previous work showing that microglia do not polarize toward spheroids and that spheroids are very long lasting [4, 31, 32], we suggest that in this case, C3 is not likely to be playing a role in glial engulfment. The precise role of C3 within spheroids will require additional investigation in the future.

The PI3K/AKT/mTOR axis emerged as a key activated signaling pathway within PAAS (**Figures 3C and S9E**). Additionally, the activation of this pathway in human postmortem brains showed a strong correlation with AD (**Figures 5B-C**). The PI3K/AKT/mTOR pathway is known for its inhibitory effects on cell autophagy, endosome, autophagosome maturation [79], lysosomal biogenesis and proteasome assembly [52], while simultaneously promoting axonal outgrowth [62, 63, 80] and lipid synthesis [52, 81]. Interestingly, certain lipids like phosphatidic acid and cholesterol have been shown to activate the mTORC1 complex [82, 83]. This suggests that lipids present within PAAS could mediate the activation of mTOR signaling, which in turn may play a role as a modulator of PAAS formation by regulating the three key biological processes that we identified within PAAS (lipid transport, protein turnover and cytoskeletal dynamics) (**Figures 3** and **4**).

The pharmacological and genetic inhibition of mTOR signaling in our human iPSC-derived AD model led to a marked reduction in PAAS pathology (**Figures 6** and **7**). Moreover, by using hSyn promoter driven Cre viruses in vivo, we achieved neuron-specific mTOR heterozygous knockout in 5XFAD mice, thereby avoiding potential confounding effects of mTOR manipulations in glial cells. Investigation of downstream mechanisms revealed that mTOR knockdown enhanced autophagy, which could operate at the whole cell level and locally at axonal spheroids, considering the extensive accumulation of endolysosmal vesicles in these structures. Furthermore, we noted the presence of mRNA and nascent proteins within axonal spheroids, indicating a potential for local translation at these sites. Although mTOR signaling is known to modulate local translation in neurons in various contexts [62, 63, 84], our study did not show changes in local translation levels following mTOR knockout or pharmacological inhibition (**Figure S21D-F**).

In addition to the PI3K/AKT/mTOR, we observed the activation of other signaling pathways within PAAS, including calcium signaling and amyloid processing (**Figures 3C** and **8**). Regarding calcium homeostasis, proteins involved in calcium signaling like CAMK2 and calmodulin were identified in PAAS (**Figures 3E and Table S2**), suggesting a significant role of local calcium signaling dysregulation in PAAS pathogenesis. This is supported by our longitudinal calcium imaging in human neurons, which demonstrated that axonal spheroids disrupt calcium rise and decay times following electrical stimulation, indicating impaired action potential conduction and calcium homeostasis within PAAS. Recent studies have shown that abnormal local calcium efflux from deacidified late endosomes and amphisomes can affect axonal transport of these vesicles [85], indicating a complex interaction between various signaling pathways during PAAS formation and growth.

Our human iPSC-derived AD model successfully replicated PAAS pathology, characterized by abundant spheroid formation around thioflavin S positive amyloid plaques and significant accumulation of endolysosomal vesicles, cytoskeletal elements, and phosphorylated tau within PAAS. The administration of exogenous β-amyloid in this model (**Figure 6A-D**) supports the theory that spheroids are formed in response to extracellular amyloid deposition rather than being the source of these deposits [18, 86]. Longitudinal imaging of individual axons in human neurons showed rapid spheroid formation and lysosome accumulation within days after Aβ administration. Additionally, axons showed active growth during spheroid formation, as opposed to a dying-back pattern (**Figure 6C**), consistent with our previous *in vivo* observations in mice [4]. Despite widespread exposure to diffuse Aβ in this model, abundant axonal spheroids were only observed next to compact thioflavin S positive deposits (**Figure 6B, Figures S16E-F and S18B-C**). This is consistent with observations in humans showing that diffuse amyloid deposits (thioflavin S negative) are not typically surrounded by PAAS, whereas thioflavin S positive deposits usually have extensive PAAS pathology [31]. This strongly suggests that specific changes in β amyloid conformation are critical for inducing PAAS formation. Applying Torin1 before amyloid plaque treatment can answer whether mTOR inhibition has an effect on axonal spheroid initiation, whereas applying Torin1 after amyloid plaque treatment and axonal spheroid formation can answer whether mTOR inhibition has an effect on axonal spheroid maintenance, although spheroids likely continue to form in a gradual way following amyloid administration.

There are several potential limitations of our study. First, while our STED super-resolution imaging indicated that the radius of antibody-based proximity labeling in brain tissue is less than 50 nm, it is possible that some proteins that are distal to PLD3 were not identified in the PAAS proteome. However, the high enrichment of PLD3 in axonal spheroids increases the likelihood of protein biotinylation, as evidenced by our ability to capture most proteins previously described in these structures, including not only endolysosomal-related proteins, but also cytoskeletal proteins, such as SPTBN1, and proteins expressed on the cell surface, such as receptor NTRK2 (**Figures 2D, 8, Figure S9A, S10 and Table S2**). Second, although PLD3 is found in spheroids of both AD humans and mice, some labeling does occur at neuronal cell bodies and neuropil (∼27% of PLD3 signal) (**Figures S2E-G**). We addressed this by eliminating proteins potentially derived from neuronal cell bodies (**Figure 2B**, also see method section), but we cannot completely rule out the presence of some neuronal cell body-specific proteins in the PAAS proteome. For example, in theory if a protein is up-regulated in AD and expressed only in the neuronal soma but not in PAAS, that could constitute a potential false positive in this dataset, although we have not found such situation thus far. Third, the low number of plaques and axonal spheroids in early-stage AD limited our ability to compare the levels of the various PAAS proteins at different stages of disease progression due to limited protein yield. Future studies could employ emerging techniques for multiplexed high-resolution quantitative imaging [87] for comparisons at different disease stages. Fourth, the human iPSC-derived AD model recapitulates the formation of spheroids around amyloid plaques, but it does so through a rapid process that may not fully reflect the *in vivo* situation. Future models may need refining for a more chronic buildup of amyloid deposits and potentially include co-culturing with other cell types like microglia to more closely mimic the *in vivo* plaque microenvironment.

Altogether, the proteomics resources and methodologies we developed for analyzing axonal pathology in human postmortem brains, iPSC-derived human neurons and *in vivo* open opportunities for elucidating mechanisms controlling the initiation and progression of axonal pathology, and for testing new therapeutic targets. While this study focused on axonal spheroids associated with amyloid plaques, spheroids can be found in other neurodegenerative disorders [21, 88–90]. Therefore, the multidisciplinary strategies established here will facilitate future studies into the diverse cell biological processes governing axonal spheroid pathology in AD and other neurodegenerative disorders. Ultimately, this will provide crucial insights into the role of axonal pathology in neurodegeneration and neural circuit disruption, with significant therapeutic implications.

## ONLINE METHODS

### Human postmortem brain tissue

Snap-frozen postmortem human brain specimens of frontal cortices from AD patients and age-matched controls were obtained from the Yale Alzheimer’s Disease Research Center and the Banner Sun Health Research Institute. Detailed demographic and clinical information can be found in **Figure S1**. For proximity labeling proteomics, 6 AD cases with intermediate to high AD level [64] and 8 age-matched unaffected controls were used. To reduce inter-sample variability and maximize signal-to-noise by avoiding brains with low-density amyloid deposition, we carefully inspected ∼ 40 individual postmortem brains using microscopy and selected for proteomic analysis 6 AD brains with the highest density of amyloid plaques and axonal spheroids within the frontal cortex. The grey matter regions with high plaque load were microdissected out from AD brain sections under visual guidance using a fluorescence stereomicroscope(Leica). Similarly, the grey matter regions were dissected from unaffected control brain sections. For immunofluorescence proteomic validations, 25 severe AD and 14 unaffected control cases were used (**Figure S1**).

### Human iPSC line and human primary astrocytes

Two fully characterized, de-identified control human iPSC lines NSB3182-3 (female) and NSB2607 (male) were used in all experiments [94]. *NGN2*-induced glutamatergic neurons [56] were generated and co-cultured with human primary astrocytes (ThermoFisher #N7805200, or ScienCell #1800) for all experiments [57].

### Mice

All animal procedures were approved by the Institutional Animal Care and Use Committee at Yale University. WT (C57BL/6J), 5XFAD (Tg6799) mice [95] and mTOR-flox (JAX #011009) mice [61] were obtained from Jackson Laboratory. 5XFAD and WT mice, used for proximity labeling proteomics, were euthanized at 15-month-old, followed by transcardial perfusion. Three male mice per genotype (WT and 5XFAD) were used. Animals used for immunofluorescence proteomic validation were euthanized at 2-3 or 12-15 months of age, with 3 biological replicates per experiment. mTOR-flox mice were cross-bred with 5XFAD mice to create an mTOR-flox-5XFAD line. For AAV-mediated mTOR heterozygous knockout experiments, mTOR-flox-5XFAD mice were injected with AAVs at 6 weeks of age. Five biological replicates (combining male and female mice) in each group were used for AAV9-hSyn-cre-2a-tdT experiment, and 3 male mice in each group were used for AAV-PHPeB experiment.

### Antibodies and reagents

Full list of primary antibodies for newly validated and known PAAS proteins can be found in **Table S2**, including catalog number, RRID, dilution factors and brief staining instructions. Briefly, Anti-PLD3 antibody, anti-SMI312 and anti-Cathepsin D were used to label PAAS in both mice and humans. Anti-Lamp1 and anti-Cathepsin B antibodies were used to label PAAS in mice. For proteomic hits validation, anti-GAA, anti-GBA, anti-TPP1, anti-ATP6V0A1, anti-SYT11, anti-G3BP1, anti-G3BP2, anti-ITM2B, anti-SPTBN1, anti-SV2A, anti-ATP2B3, anti-CAMK2A, anti-Calmodulin, anti-SYT1, anti-CACNA2D1, anti-CACNA1B, anti-NTRK2, anti-mTOR, anti-p-mTOR S2448, anti-PIK3R4, anti-AKT1, anti-LAMTOR, anti-RAGA, anti-RAGC, anti-RHEB, anti-RAPTOR, anti-HDLBP, anti-APOE, anti-C3, anti-HEXB and anti-TMEM30A were used for validating newly identified PAAS proteins. Anti-RAGC, anti-ATG9A, anti-Ubiquitin, anti-RTN3, anti-PKC, anti-synaptophysin, anti-SNAP25, anti-VAMP2 and anti-beta Tubulin III were used for immunostaining of known PAAS proteins. To reveal GFP and tdTomato protein expression, anti-GFP (1:500, RRID:AB_10000240) and anti-RFP (1:200, RRID:AB_2209751) were used respectively. For staining neuronal and glial markers in iPSC-derived human neurons, anti-neurofilament H (1:1000, RRID:AB_2149761), anti-NeuN (1:1000, RRID:AB_10711040), anti-NeuN (1:200, RRID:AB_2532109), anti-Synapsin1/2 (1:500, RRID:AB_2622240), anti-PSD95 (1:200, RRID:AB_10807979), anti-S100b (1:500, RRID:AB_2814881), anti-IBA1 (1:100, RRID:AB_2891289) were used. To stain amyloid beta deposits, anti-6e10 (1:200, RRID:AB_2565328) was used. To stain phosphorylated Tau, anti-phospho-Tau S235 (1:1000, Thermo-Fisher), anti-phospho-Tau S396 (1: 200) and anti-phospho-Tau S404 (1:200) were used (RRID see **Table S2**). Dendritic marker MAP2 (1:200, RRID:AB_776174) was used. For puromycylation, anti-puromycin-647 (1:1000, RRID:AB_2736876) was used. Thioflavin S (Sigma-Aldrich, T1892, 2% w/v stock solution, 1:10,000 staining) was used to label amyloid plaques. Alexa dye-conjugated secondary antibodies were used (1:600, ThermoFisher Scientific).

### Tissue fixation

We have compared the impact of different tissue fixation approaches on the proximity labeling efficacy and protein extraction efficiency. For human brain samples, snap-frozen postmortem human brains coupled with fresh fixation in 4% PFA at 4 °C for ∼24 hours worked the best. For mice, freshly perfused mouse brains and fixed in 4% PFA at 4 °C for ∼24 hours performed the best in both proximity labeling efficacy and protein extraction efficiency. We found that long-term fixation and storage with the FFPE method markedly reduced both proximity labeling and protein extraction efficiencies.

### Proximity labeling in brain tissue

Proximity labeling in human and mouse tissue was performed based on [43] with optimizations. Detailed procedures are described below. Axonal spheroids were proximity labeled by using anti-PLD3 or anti-Lamp1 antibodies in mice and humans. Neuronal soma were proximity labeled using anti-NeuN antibody. Briefly, frozen postmortem human brain specimens were fixed by submerging into ice cold 4% paraformaldehyde, and put onto a shaker at 4 °C for ∼ 24 hours. For mice, after transcardial perfusion, brains were fixed in 4% paraformaldehyde at 4 °C for ∼ 24 hours while shaking. Human and mouse brains were vibratome sectioned at 50 μm thickness. Ten sections (around 1cm x 0.8cm each) for human or mouse brain were used in each reaction/per biological replicate. Human sections contained mostly grey matter. Six to eight human biological replicates were used in each group, and three biological replicates were used in each mouse group. Sections were permeabilized by PBS with 0.5% Triton X-100 for 7 min, followed by rinsing with PBST (0.1% Tween-20 in PBS). To quench the endogenous peroxidase activity, sections were incubated in 0.1% H2O2 for 10 min, followed by rinsing with PBST twice. Primary antibody diluted in blocking buffer (0.1% Tween-20 with 1% BSA in PBS) was incubated overnight at 4 °C on a shaker, followed by PBST washes for 3 times 20 min/per wash. Secondary antibody conjugated with HRP was incubated in blocking buffer for 1 hour at room temperature, followed by PBST washes for 3 times ∼40 min/per wash. Proximity labeling was performed by using Biotin-XX-Tyramide dissolved in 50 mM Tris-HCl buffer (pH = 7.4), with H2O2 for 5 min, according to the user’s manual (ThermoFisher Scientific B40921). Specifically, every 1 mL of reaction solution was made of 10 μL 1X Biotin-XX-Tyramide and 10 μL 1X H2O2 in 50 mM Tris-HCl buffer. Biotinylation reactions were terminated by rinsing sections with freshly made 500 mM sodium ascorbate for 3 times, followed by PBST washes for 3 times.

### Enrichment of biotinylated proteins using streptavidin beads

We performed proximity labeling proteomics on fixed brain specimens, which may reduce the protein extraction yield compared to other proximity labeling methods using fresh tissue. Thus, we optimized the protein extraction protocol and largely increased the protein extraction yield compared to previously published methods (**Figure S4A**)[43]. Specifically, brain sections from proximity labeling experiments were lysed and de-crosslinked in 100 mM Tris-HCl buffer (pH = 8.0) with 2% SDS and protease inhibitor (Roche) at 95 °C for 45 min with constant shaking. For every 10 brain sections, 500uL lysis buffer was used. Protein lysate was sonicated using Sonic Dismembrator Model 500 (Fisher Scientific) for 3 times, 3 sec/per time at 4°C. Protein lysate was centrifuged at 12k rcf for 5 min. Then, 450 µL protein lysate supernatant was collected from each sample, incubated with 550 µL of PBST containing 200 µL prewashed streptavidin magnetic beads (ThermoFisher Scientific #88817), protease inhibitor and phosphatase inhibitor, to meet a final 1mL volume. Samples were then incubated on a 360° rotator at 4 °C overnight. The rest of protein lysates were used for protein concentration measurement by BCA (ThermoFisher Scientific). After incubation, beads were sequentially washed once with PBST, twice with PBST with 1 M NaCl, twice with PBS. Biotinylated proteins were eluted in elution buffer (20 µL of 20 mM DTT and 2 mM Biotin in 1X NuPAGE LDS lysis buffer (ThermoFisher) with protease inhibitor and phosphatase inhibitor) at 95 °C for 5min. Supernatant was collected and centrifuged at 12k rcf for 1min, followed by running into a 4%–20% Tris-Glycine gel (Invitrogen) at constant 150 V until all the proteins had run into the gel (approximately 10min). Gel was rinsed once in ultrapure water (AmericanBio) and incubated in ∼50 mL of Coomassie blue R-250 staining solution (Bio-Rad) for 1 hour incubation. Gel was de-stained with Coomassie blue R-250 destaining solution (Bio-Rad) for 2 hours with 3 times buffer changes. Gel was rinsed with ultrapure water for three times. Gel containing protein samples was visualized, cut with clean blades, and kept at -20 °C.

### Enrichment of biotinylated peptides using Anti-biotin antibody

The labeled tissue was lysed using 100 mM Tris-HCl buffer (pH = 8.0) with 2% SDS and protease inhibitor (Roche). The lysates were sonicated and then centrifuged at 16,500 g for 10 min at 4°C. The proteins were precipitated using acetone and the pellet was dissolved in 8M urea, 50 mM ammonium bicarbonate (ABC), then sonicated for 30 sec to re-solubilize the proteins. A Bradford Assay was performed to determine protein concentration and 2 mg of protein was used to processed further. Proteins were reduced with 5 mM DTT for 45 min at RT and subsequently carbamidomethylated with 10 mM iodoacetamide for 30 min at RT in the dark. Prior to digestion, the urea concentration was reduced to 2 M with 50 mM ABC) and digested with trypsin at an enzyme:substrate ratio of 1:50 overnight at 37 °C. Following digestion, samples were acidified with 10% formic acid and de-salted using Nest Group C18 macro-spin columns (HMMS18V) as per manufacturer’s instructions. Biotinylated peptides were enriched using anti-biotin antibody-based immunoprecipitation (IP). The peptides were dissolved in IAP buffer containing 50 mM MOPS, 10 mM HNa2PO4, 50 mM NaCl at pH 7.5. Anti-biotin beads (Immune Chem Pharmaceuticals) were washed twice in IAP buffer before the samples were added to the beads for incubation on a rotator for 2 hrs at 4°C. The beads were washed twice with IAP buffer and twice with H2O (HPLC-grade). The biotinylated peptides were eluted from the beads using 80% ACN and 0.15% TFA with vortexing followed by a 10 min incubation at room temperature. The elution was repeated twice, and the supernatants were collected and vacuum dried.

### Western bhlotting

For western blot, 4%–20% Tris-Glycine gels (Invitrogen) were used for protein electrophoresis following the manufacturer’s protocol. Proteins were transferred to nitrocellulose membranes (BioRad) at constant 350 mA for about 50 min. After blocking with 5% bovine serum albumin (BSA) in TBST (Tris-buffered saline with 0.1% Tween 20) for 1 hour, membranes were incubated with primary antibodies (anti-PLD3 1:250; anti-CatB 1:1000, anti-RAGC 1:1000, anti-NeuN 1:1000; anti-GAPDH 1:1000) diluted in 5% BSA in TBST on a shaker at 4 °C overnight, followed by three times of 15min washes with TBST. Membranes were then incubated with horseradish peroxidase (HRP) conjugated secondary antibodies diluted in 5% BSA in TBST for 1 hour at room temperature, followed by three times of 15min washes with TBST. To blot biotinylated proteins on the same membrane, stripping buffer (ThermoFisher #46430) was used to cover the whole membrane and incubated at room temperature with shaking for 10-12 min, followed by rinsing with PBST for 3 times. HRP-conjugated streptavidin (1:1000) was diluted in blocking buffer for 1-2 hours at room temperature or 4 °C overnight. Clarity Western ECL blotting substrate (Bio-Rad) and ChemiDoc MP imaging system (Bio-Rad) were used for chemiluminescence development and detection.

### In-Gel Digestion

Gel slices were cut into small pieces and washed with 600 µL of water on a tilt-table for 10 min followed by 20 min wash with 600 µL 50% acetonitrile (ACN)/100 mM NH4HCO3 (ammonium bicarbonate, ABC). The samples were reduced by the addition of 100 µL 4.5 mM dithiothreitol (DTT) in 100 mM ABC with incubation at 37°C for 20 min. The DTT solution was removed and the samples were cooled to room temperature. The samples were alkylated by the addition of 100 µL 10mM iodoacetamide (IAN) in 100 mM ABC with incubation at room temperature in the dark for 20 min. The IAN solution was removed, and the gels were washed for 20 min with 600 µL 50% ACN/100 mM ABC, then washed for 20 min with 600 µL 50% ACN/25 mM ABC. The gels were briefly dried by SpeedVac, then resuspended in 100 µL of 25 mM ABC containing 500 ng of digestion grade trypsin (Promega, V5111), and incubated at 37°C for 16 hrs. The supernatants containing the tryptic peptides were transferred to new Eppendorf tubes. Residual peptides in the gel bands were extracted with 250 µL 80% ACN/0.1% trifluoroacetic acid (TFA) for 15 min, then combined with the original digests and dried in a SpeedVac. Peptides were dissolved in 24 µL MS loading buffer (2% acetonitrile, 0.2% trifluoroacetic acid), with 5 µL injected for LC-MS/MS analysis.

### LC-MS/MS on the Thermo Scientific Q Exactive Plus

LC-MS/MS analysis was performed on a Thermo Scientific Q Exactive Plus equipped with a Waters nanoAcquity UPLC system utilizing a binary solvent system (A: 100% water, 0.1% formic acid; B: 100% acetonitrile, 0.1% formic acid). Trapping was performed at 5 µL/min, 99.5% Buffer A for 3 min using an ACQUITY UPLC M-Class Symmetry C18 Trap Column (100Å, 5 µm, 180 µm x 20 mm, 2G, V/M; Waters, #186007496). Peptides were separated at 37 °C using an ACQUITY UPLC M-Class Peptide BEH C18 Column (130Å, 1.7 µm, 75 µm X 250 mm; Waters, #186007484) and eluted at 300 nL/min with the following gradient: 3% buffer B at initial conditions; 5% B at 2 min; 25% B at 140 min; 40% B at 165 min; 90% B at 170 min; 90% B at 180 min; return to initial conditions at 182 min. MS was acquired in profile mode over the 300-1,700 m/z range using 1 microscan, 70,000 resolution, AGC target of 3E6, and a maximum injection time of 45 ms. Data dependent MS/MS were acquired in centroid mode on the top 20 precursors per MS scan using 1 microscan, 17,500 resolution, AGC target of 1E5, maximum injection time of 100 ms, and an isolation window of 1.7 m/z. Precursors were fragmented by HCD activation with a collision energy of 28%. MS/MS were collected on species with an intensity threshold of 1E4, charge states 2-6, and peptide match preferred. Dynamic exclusion was set to 30 sec.

### Peptide Identification

Data was analyzed using Proteome Discoverer software v2.2 (Thermo Scientific). Data searching was performed using the Mascot algorithm (version 2.6.1) (Matrix Science) against the SwissProtein database with taxonomy restricted to human (20,368 sequences) or mouse (17,034 sequences) as well as a streptavidin sequence. The search parameters included tryptic digestion with up to 2 missed cleavages, 10 ppm precursor mass tolerance and 0.02 Da fragment mass tolerance, and variable (dynamic) modifications of methionine oxidation and carbamidomethyl cysteine. Normal and decoy database searches were run, with the confidence level set to 95% (p<0.05). Scaffold v5.1.2 (Proteome Software Inc., Portland, OR) was used to validate MS/MS-based peptide and protein identifications. Peptide identifications were accepted if they could be established at greater than 95.0% probability by the Scaffold Local FDR algorithm. Protein identifications were accepted if they could be established at greater than 99.0% probability and contained at least 2 identified peptides (one uniquely assignable to the protein). Proteins that contained similar peptides and could not be differentiated based on MS/MS analysis alone were grouped to satisfy the principles of parsimony. Proteins sharing significant peptide evidence were grouped into clusters. Label-free quantification was performed with Scaffold software. Spectral intensity values were used for protein quantification between groups.

To search for biotinylation sites in the anti-biotin antibody pulldown samples, the variable modifications of biotin-xx-tyramide were configured to account for marker ions resulting from fragmentation of biotinylated peptides. These marker ions have the following m/z values and elemental composition losses from the fully modified amino acid: dehyrdrobiotin (m/z-227.08), Biotin-x ion (m/z-340.25), Biotin-xx ion (m/z-453.25), immonium of tyrosine-Bxxp with loss of ammonia (m/z-706.38), immonium of tyrosine-Bxxp (m/z-723.38).

### Proteomic data analysis

Prior to the data analysis, missing values were removed. For example, if a protein A had 0 spectra count detected in all the samples, including tests and controls, then protein A was removed from the list. Normalized total spectra count (NTSC) and normalized total precursor intensity (NTPI) are two common methods for proteomic quantification. For PLD3-labeled PAAS proteomes in humans, we compared NTSC and NTPI methods, and used the shared proteomic hits for downstream analysis. For PLD3-labeled PAAS proteomes in mice, Lamp1-labeled proteomes in mice and NeuN-labeled neuronal nuclei and perinuclear cytoplasm proteomes in mice, we used NTSC for quantification. To obtain the PAAS proteome, differentially expressed proteins were analyzed by comparing proteomic hits obtained from PLD3-labeled samples versus those from control samples using no antibody. This allowed the filtering of endogenously biotinylated proteins and non-specific binders to streptavidin beads. To obtain the optimal cutoff values for the statistical analysis we tested different degrees of stringency for FDR (0.1, 0.05 and 0.01) and fold change (1 and 1.5). An optimum cutoff p < 0.05, FDR < 0.1 and fold change > 1.5 was used for these datasets, as it captures the maximum numbers of known PAAS proteins while excluding the maximum numbers of potential contaminants. Post-cutoff proteomic lists were scrutinized for possible glial contaminations by cross validations using single cell RNAseq (scRNAseq) transcriptomics in mice [55] and humans [96] and Tissue Atlas in The Human Protein Atlas [97]. When a gene had an FPKM < 10 in neurons and an FPKM > 10 in other cell types in the mice scRNAseq dataset [55], or the mean expression level was less than 0.03 in neurons but greater than 0.03 in glia in the AD pathology human scRNAseq dataset [96], and protein expression was not detected in neurons from the Tissue Atlas [97], this gene was excluded from the proteomic results. Two such genes (EPHX1 and PSAT1) were excluded from the PAAS proteome in AD humans, and five such genes (Gsn, Ephx2, Gfap, Myh9 and Anxa2) were excluded from the PAAS proteome in AD mice. Proteomic hits that passed these thresholds were considered the final PAAS proteomes in AD humans or mice. Lists of raw and filtered proteomic hits of PAAS and neuronal soma proteomes in AD humans and mice can be found in **Table S1**.

For gene ontology analysis (Figure 3A**, Figures S9C and S13B**), we uploaded the final proteomes to the GeneOntology.org [98, 99], the g:profiler [100] or the ToppGene Suite search portal [101] and plotted the top ten or top twenty retrieved terms on cellular compartment or biological process with the lowest false discovery rates. Pathway enrichment analysis was performed by retrieving Gene Ontology Biological Process, Cellular Component and Molecular function terms from g:profiler [100] using terms size 5-200. The enrichment map was visualized in Cytoscape (v3.9.1) [102]. For IPA pathway analysis (Figures 3B-C **and Figures S9D-E**), the human or mice PAAS proteome was imported into Ingenuity Pathway Analysis (IPA) software (QIAGEN, 2022 release version)[103] for canonical pathway analysis. The top IPA pathways with the lowest false discovery rates and potential relevance to PAAS pathology were listed. For gene set enrichment analysis (GSEA) (Figure 4A), PLD3 labeled proteomes of AD humans and unaffected controls were uploaded into the Broad Institute GSEA software 4.3.2 [104] to perform gene ontology analysis using default values, except the size was set to 200 to remove the larger sets from analysis. GSEA results were loaded into Cytoscape (v3.9.1) for pathway enrichment analysis using EnrichmentMap [105] and AutoAnnotate [106] plugins with default values. Principal component analysis was performed using Qlucore Omics Explorer v3.6 (Qlucore AB, Lund, Sweden).

#### Immunofluorescence of fixed specimens and human iPSC-derived co-culture

The complete list of antibodies, dilution factors, and immunofluorescence instructions for immunofluorescence staining of fixed specimens of humans and mice, can be found in **Table S2**. Briefly, for mice and human brains, fixation and vibratome sectioning was the same as described in the proximity labeling section. Heat-induced sodium citrate antigen retrieval was performed when necessary (see IF instructions in **Table S2**). Immunofluorescence staining was then performed with the following protocol: tissue was boiled in 50 mM sodium citrate with 0.05% tween-20 at 95 °C for 45 min, followed by 30 min cool down at RT and rinse with PBS for three times. Primary antibodies incubation was 12 hours to 3 days at 4 °C in blocking buffer (PBS with 1% BSA and 0.1% Tween-20) and secondary antibodies were incubated in blocking buffer at 4 °C overnight. Thioflavin S (Sigma-Aldrich, T1892, 2% w/v stock solution, 1:10,000 staining) was used for labeling amyloid deposits. Three times washes with PBST were performed before mounting tissues on slides with PermaFluor (Thermo Scientific, TA-030-FM).

For immunofluorescence of human iPSC-derived neuron-astrocyte co-culture, cells were washed 3 times with pre-warmed PBS before fixing with 4% PFA (ice-cold) at RT for 20 min. Cells were washed with PBS 15 min x 3 and were blocked with blocking buffer (1% BSA in PBS, plus 0.1% tween-20) for 1 h. Primary antibodies were diluted in blocking buffer and incubated with cells at 4 °C overnight. Cells were washed with PBST 15 min x 3. Secondary antibodies were diluted in blocking buffer and incubated with cells at 4 °C overnight. ThioflavinS (2% w/v stock solution, 1:10,000 staining) was diluted in PBS and incubated with cells at RT for 5 min. Cells were washed with PBST 15 min x 3, before imaging.

### Fixed tissue and live cell time-lapse confocal microscopy imaging

An upright or an inverted Leica SP8 confocal microscope was used to generate all images. Laser and detector settings (GaAsP hybrid detection system, photon counting mode) were maintained constant. For all analyses, single z stack images or tiled images were obtained in the somatosensory cortex in mice. All images were obtained using a 63x oil immersion objective (N.A. 1.40), 40x water immersion objective (N.A. 1.10) or 25x water immersion objective (N.A. 0.95) at 1,024 x 1,024-pixel resolution, z-step size of 1 μm, as we previously described [31]. When indicated, deconvolution was performed using the default setting in the Leica SP8 LAS X software. For time-lapse imaging of spheroids growth in human neurons, 96-well plates were imaged in the Leica SP8 incubator, with the temperature set at 37°C and supplied with CO2. Tilling images were obtained using a 63x oil immersion objective (N.A. 1.40) at 512 x 512-pixel resolution, zoom factor at 3 and z-step size of 1 or 1.5 μm at every day for 7 days.

### Lentivirus plasmid purification, lentivirus production and concentration

*Escherichia coli* stocks for pMDLg/pRRE (MDL), pRSV-Rev (Rev), pCMV-VSV-G (VSVG), FUW-M2rtTA and pLV-TetO-hNGN2-eGFP-Puro were purchased from Addgene (12251, 12253, 8454, 20342 and 79823, respectively). Bacteria was grown in 500 mL of LB Broth (Fisher Scientific, DF0446-07-5) with 100 µg/mL Ampicillin (Sigma, A9518) overnight at 37 °C and 200 rpm. The following day, bacterial cells were pelleted by centrifugation at 4000 g for 10 min at 4 °C. The supernatant was discarded, and plasmids purified using the PureLink™ HiPure Plasmid Filter Maxiprep Kit (Invitrogen, K210017), following the manufacturer instructions.

Third generation lentiviral vectors were produced as we described [107]. Briefly, HEK 293T cells (Invitrogen, R700-07) were grown in 15-cm plates (Falcon, 353025) in DMEM (Gibco, 12430054) supplemented with 10% FBS (Gibco, 10438026) until reaching 70-80% confluency. For cell transfection, the following solution was prepared in 500 µL of pre-warmed Opti-MEM (Gibco, 31985062) per plate: 12.2 µg of transfer plasmid, 8.1 µg of MDL, 3.1 µg of Rev, 4.1 µg of VSVG and 110 µL of Polyethylenimine (1 µg/µL; Polysciences, 23966-2). This solution was incubated for 10 min at room temperature, vortexed gently and added dropwise onto HEK 293T cells for transfection. Medium was changed after 6 h of incubation and harvested at 48 h and 72 h post transfection. Media containing viral particles was sterile-filtered and concentrated using the Lenti-X™ Concentrator (Takara Bio, 631231), according to the manufacturer instructions.

### Human neuron-astrocyte co-culture AD model

Human iPSC control lines 3182-3 and 2607 were used in this study, as we first described this line in [94]. Maintenance and passaging of human iPSC were performed as we previously described [108]. Human primary astrocytes (ThermoFisher #N7805200 or ScienCell #1800) were maintained as described in the user manual. iPSC-derived NGN2-induced glutamatergic neuron generation was performed as we previously described [108]. Briefly, human iPSCs maintained in 6-well-plates were harvested by incubating in Accutase (Innovative Cell Technologies AT104) 1 mL/per well plus 10 µM ROCK inhibitor THX (RI) (Tocris #1254) at 37 °C for 20 min. Dissociated iPSCs were then collected in a 50 mL falcon tube and mixed well with DMEM (ThermoFisher #11966025) (preferably 1:3 Accutase : DMEM). iPSCs were centrifuged for 4 min at RT at 1000 g. iPSCs were resuspended in 1-2 mL of Stemflex (ThermoFisher) with 10 µM RI and counted, diluted in Stemflex with THX to a cell suspension concentration of 1e6 cells/mL. Lentiviruses NGN2-Puro and rtTA (titer 4.40 x 10^10 gc/mL) at 50 uL per 10e6 cells of suspension were added. Cells were mixed and dispensed at 120k cells/per well (6 well size) coated with 1x Geltrex (ThermoFisher). Cells were incubated at 37 °C incubator overnight. DIV1: media was replaced with induction media (DMEM F12 with Glutamax and Sodium Pyruvate (Thermofisher, #10565018), 1% N-2 (Thermofisher, #17502048), 2% B-27-RA (Thermofisher, #12587010), Doxycycline for a final concentration of 1ug/mL). DIV2 and 3: replaced media with induction media containing puromycin (2 ug/mL) on each day. DIV4: Neurons were dissociated with Accutase plus 10 µM RI for 20 min, washed off with 1:3 DMEM, resuspended and centrifuged at 1000 g for 5 min. Resuspend pellet at a concentration of 1e6 cells/ml in neuron media (Brainphys (STEMCELL, # 05790), 1% N-2, 2% B-27-RA, 1 μg/mL Natural Mouse Laminin (Thermofisher, # 23017015), 20 ng/mL BDNF (R&D, #248), 20 ng/mL GDNF (R&D, #212), 250 μg/mL Dibutyryl cyclic-AMP (Sigma, #D0627), 200 μM L-ascorbic acid (Sigma, # A4403) and 1x Anti-Anti (ThermoFisher)) with Dox 1μg/mL, puromycin 2 μg/mL, 4 µM AraC (Sigma #C6645) and 10 µM RI. Neurons were replated onto 2x Geltrex coated 96-well-plates (PerkinElmer #6055302) and seeded at 10e5 per well (96-well plate size). DIV5: media was replaced with neuron medium containing puromycin 2 μg/mL and AraC 4 μM. DIV6: media was replaced with neuron medium with 4 μM AraC. DIV8: media was replaced with neuron medium with 2 μM AraC. DIV10: human primary astrocytes were dissociated with TrypLE (ThermoFisher), washed with DMEM, centrifuged at 400 rcf for 5 min, counted and plated into NGN2 neurons culture at 20k cells/per well (96-well-plate size). Human iPSC-derived neuron-astrocyte co-culture was maintained as previously described [57] with modifications. Briefly, human neuron-astrocyte co-cultures were maintained in neuron media plus 1.5% FBS for 1 week, then reduced FBS to 0.5% for another week. Media was half-changed every other day. After that, human neuron-astrocyte co-cultures were maintained in neuron maintenance medium (1X BrainPhys Basal (StemCell Technology), 1X B27 with Vitamin A (ThermoFisher), 1X N2 (ThermoFisher), 5 μg/ml Cholesterol (Sigma-Aldrich), 1 mM Creatine (Sigma-Aldrich), 10 nM β-estradiol, 200 nM Ascorbic Acid, 1 mM cAMP (Sigma-Aldrich), 20 ng/ml BDNF (Peprotech), 20 ng/ml GDNF (Peprotech), 1 μg/ml Laminin, 0.5 mM Glutamax (ThermoFisher), 1 ng/ml TGF-β1 (Peprotech), 1X Normocin (InvivoGen), 50 U/ml Penicillin-Streptomycin (ThermoFisher)) with half-changed of media every other day until harvest, or other assays. For AD modeling, amyloid beta 1-42 peptide (AnaSpec AS-72216) was oligomerized to prepare soluble amyloid beta species as previously described [57, 109]. Briefly, soluble amyloid beta species were added to the neuron maintenance medium at a final concentration of 5 μM and applied to the human neuron-astrocyte co-culture for 7 days, with half-change of media every other day as previously described [57].

For mTOR signaling inhibition, Torin1 (TOCRIS 4247) or vehicle DMSO was added to the neuron maintenance medium at a final concentration of 250 nM [59], and applied to the human neuron-astrocyte co-culture 3 days before amyloid beta treatment. Then, Torin1 or vehicle DMSO was added to the neuron maintenance medium at 250 nM concentration, along with soluble amyloid beta species at 5 μM concentration, and applied to the human neuron-astrocyte co-culture for 7 days, with a half-change of medium every other day. For Torin1 treatment after axon spheroids formation, soluble amyloid beta species at 5 μM concentration, and applied to the human neuron-astrocyte co-culture for 7 days, then Torin1 or vehicle DMSO was added to the neuron maintenance medium at 250 nM concentration for another 7 days.

For time-lapse imaging of spheroid growth in human neurons, AAV9-hSyn-mCherry (Addgene #114472) and AAV2-CAG-LAMP1-GFP (home produced as we previously described [4]) were co-transduced at 2 x 10^9 vg/100 μL medium in 4-month-old co-cultures. Human neuron-glia co-cultures were treated with amyloid at day 150. Time lapse confocal imaging was performed several hours before the treatment, and imaged every day post-treatments.

### Calcium imaging in iPSC-derived human neurons

AAV1-CAMKII-GCamp8f (Addgene #176750, 7 x 10^12 vg/mL) was transduced in 2-month-old iPSC-derived human neurons, along with AAV9-CB7-mCherry (Addgene # 105544, 1 x 10^13 vg/mL) at 1 μL per 0.2 mio cells. Transduced cells were maintained with regular media changes. Calcium imaging was performed one month after transduction. Calcium imaging was performed using an Andor Dragonfly spinning disk confocal on Nikon Ti2-E microscope. A 20×/0.75 NA air objective and Zyla scientific complementary metaloxide semiconductor (sCMOS) camera were used. Imaging was performed in culture medium, a stage-top incubator and objective heater (Oko-Lab) maintained the sample temperature at 37°C. For stimulated calcium imaging, a pair of tinfoil wire electrodes separated around 1.5 mm apart was placed in the in the bottom of the well using a micromanipulator and utilized for electrical stimulation. Electric stimulation was performed using the trigger out function in the Andor software. GCaMP8f-labeled neurons were imaged through excitation at 488 nm wavelength with the acquisition speed of 20 Hz, and 512 x 512 resolution. Stimulation trains of 20 ms pulses were delivered to the electrodes at 50 Hz (20 ms interval) with 300 to 700 μA currents for 1s. Electric stimulation was triggered automatically at the 100-frame time point, and image for 300 or 500 frames in total. The calcium responses within the imaging window were monitored upon stimulation.

Calcium imaging raw data were de-noised using DeepCAD-RT [110]. The denoising model was trained with the different datasets acquired with the same imaging conditions. The denoised GCaMP8f fluorescence intensity was normalized to ΔF/F for analysis. Several regions of interest (ROIs) were selected on each axon. The average ΔF before stimulation was used as the baseline measurement. The raw fluorescence intensity over time (F(t)) contained background noise. To isolate the stimulus-evoked signal, we first calculated the average background fluorescence intensity (F0) using data from before stimulus presentation. We then normalized the signal by subtracting F0 at each time point:

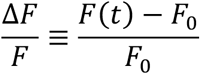

This normalized signal was smoothed using a Savitzky-Golay filter with a window length of 21 data points (1 second) and a polynomial order of 3. This filtering reduced noise while preserving the overall shape of the signal. To quantify the rising slope of the calcium response, we identified the peak of the smoothed signal. The time point halfway to this peak (𝑡_1/2_) was then determined. A linear regression was performed using the 9 data points surrounding 𝑡_1/2_. The slope of this regression line provided an estimate of the rising slope of the calcium signal in response to the sensory stimulus.

To determine the onset time of the response for each Region of Interest (ROI) following electrical stimulation, we employed an exponential growth model to fit the rising phase of the signal. Prior to reaching its peak, the normalized calcium trace can be effectively approximated using the following piecewise function:

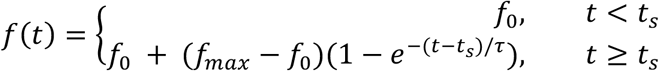

The curve fitting procedure was performed using the curve_fit function available in the scipy.optimize package. In this context, 𝑓_0_ and 𝑓_max_ represent the baseline and maximum normalized calcium signals, respectively. The parameter 𝜏 characterizes the timescale of the rising phase, while 𝑡_*s*_ denotes the response time—the specific time point at which the calcium signal initiates its ascent.

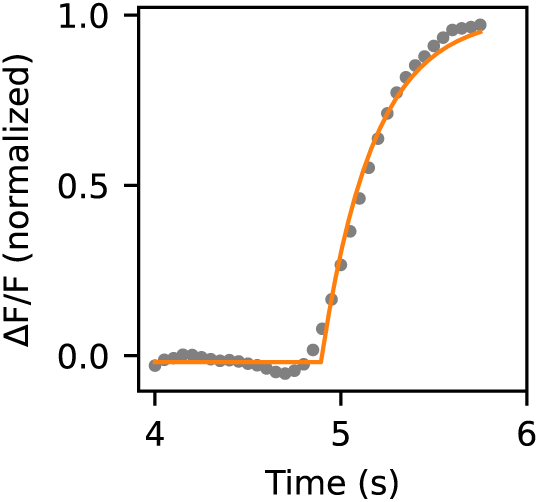

The calcium decay time constant (τ) was estimated by fitting 2.5 s calcium trace beginning from the peak to an exponential equation: Y = a*exp(-x/τ).

### Viral-mediated *Mtor* heterozygous knockout in *Mtor*-floxed-5XFAD mice

mTOR is known to be a regulator of cell growth [35, 59], previous studies showed cell size was indistinguishable between the *Mtor* heterozygous knockout and wild-type cells [111], which suggests that the effects of *Mtor* knockout on cell size is *Mtor* gene-copy dose dependent. Thus, we obtained *Mtor* heterozygous floxed AD mice to achieve conditional *Mtor* heterozygous knockout, which had no effect on cell body size (Figures 6O **and S19D**). To achieve global neuronal *Mtor* heterozygous knockout in *Mtor*-floxed-5XFAD mice, 10 μL of AAV-PHPeB-hsyn-cre-eGFP (Addgene#105540, titer ∼1x10^13vg/mL) or AAV-PHPeB-CAG-GFP (Addgene#37825, titer ∼1x10^13vg/mL) was retro-orbitally injected into 6-week-old *Mtor*-floxed-5XFAD mice. To achieve sparse neuronal *Mtor* heterozygous knockout, 1 μL of AAV9-hsyn-cre-2a-tdT (Addgene#107738, titer ∼1x10^13vg/mL) was diluted with 3 μL of PBS and was injected into the subarachnoid space of one hemisphere at the level of somatosensory cortex in *Mtor*-floxed-5XFAD mice, as we previously described [31]. Mouse brains were collected 2 months after virus injection in the AAV-PHPeB experiment or at 2.5 months for the AAV9-hsyn-cre-2a-tdT injected mice. Brains were sliced, stained and imaged with confocal microscopy as abovementioned and as we previously described [31]. For sparsely PHP.eB-hSyn-Cre-GFP viruses labeling in mouse brains, 3 μL of AAV-PHPeB-hsyn-cre-eGFP (Addgene#105540, titer ∼1x10^13vg/mL) was diluted in 30 μL PBS and retro-orbitally injected.

### RNAscope in 5XFAD mice brain slices

5XFAD mice brains were freshly dissected and immediately froze in Tissue-TEK O.C.T. compound (SAKURA) and kept in dry ice with 70% ethanol for 5∼10 min, then transferred to -80 °C for long-term storage. Frozen brain blocks were sectioned at 10 μm thickness by cryostat (Leica). RNAscope was performed using RNAscope multiplex fluorescent kit V2 (ACD #323270) according to the fresh frozen tissue protocol. PolyA probe (ACD # 318631) or negative control probe (ACD #320871) was used during probe incubation. Before counterstain with DAPI and mounting media, Atto 657 NHS ester (Millipore #07376) was diluted in 1:100 and stained brain section for 10min, followed by 3-time PBS washes, 5min/per wash. Negative control samples were used to set the baseline parameters for confocal imaging, before imaging the PolyA probe samples.

NHS ester was used to label the spheroid halo because when we attempted to perform RNAscope and co-stain with axonal spheroid markers, such as PLD3, SMI312, APP, RAGC, ATP9A, and ATP6V0A1, we found these antibodies were not compatible with the RNAscope protocol, a well-known limitation of this technique. Therefore, we leveraged the knowledge from pan-expansion microscopy [112] that NHS ester can be used to non-specifically label tissues and reveal subcellular structures using confocal imaging. We applied NHS ester after RNAscope, ensuring that the NHS ester staining did not interfere with the polyA probe signal from RNAscope. Interestingly, in the cortex of 5XFAD mice, NHS ester highlighted the neuritic plaque nicely, allowing us to use it as a marker for spheroids, especially at the periphery of the halo.

### Puromycylation in live mice brains

Puromycylation was performed as previously described [113] with a few modifications. Briefly, 25 mg puromycin (Sigma #P7255) was diluted in 278 μL of DMSO to make the puromycin stock solution. Puromycin working solution includes 10 μL of puromycin stock solution, 10 μL DMSO and 80 μL PEG400 (puromycin final concentration at 9 mg/mL). Anisomycin (Sigma #9789) stock solution was made as follow: 20 μL DMSO was added to 5 mg anisomycin, and centrifuged at 200g for 3 min. Heat at 42 °C water bath for 2-3 min, until pipet completely dissolved. Then 40 μL of PEG400 was added dropwise with agitation. Solution should keep clear with no precipitation. Anisomycin working solution was made as follow: 30 μL anisomycin stock solution, 10 μL puromycin stock solution (or DMSO vehicle), and 60 μL PEG400. To make Torin1 stock solution, 10 mg Torin1 was added to 3.292 mL DMSO and 3.292 mL PEG400 (Torin1 final concentration at 2.5 mM). Torin1 working solution was made as follow: 30 μL Torin1 stock, 10 μL puromycin stock (or DMSO vehicle), and 60 μL PEG400. All the reagents were freshly made and checked there were no precipitation during the experiment. Mice craniotomy at the somatosensory cortex was done as we previously described [4]. After removing dura, vehicle or anisomycin, or Torin1 working solution was topically applied to cranial window. For anisomycin assay, solution was applied for 30 min, then quickly removed, and replaced with anisomycin plus puromycin solution for 10 min. After quick PBS washes, mice were transcardially perfused with PBS and 4% PFA. For Torin1 treatment, drugs were incubated for 2 hrs, with reapplication 3 times to avoid drying. Brains were vibratome sectioned at 50 μm thickness, sections were incubated for 20 min with coextraction buffer (50 mM Tris-HCl, pH 7.5, 5 mM MgCl2 ,25mM KCl, protease inhibitor (Roche), and 0.015% digitonin (Wako Chemicals #043-21376)). After three rinses with PBS, sections were incubated in blocking buffer (0.05% saponin, 10 mM glycine, and 5% fetal bovine serum in PBS) for 30min. Then, sections were incubated in blocking buffer with puromycin-647 antibody (Millipore MAB E343-AF647,1:1000), NeuN antibody (ab177487, 1:200) or Lamp1 (DSHB, #1D4B, 1:200) for 72 hours at 4°C. Sections were washed with PBS 3X 10 min, then incubated with secondary antibodies along with puromycin-647 at 4°C overnight, followed by PBS washes 3X 15 min.

### STED imaging

STED imaging was performed as we previously described [114]. Briefly, human postmortem brain tissues underwent processing utilizing immunofluorescence staining and proximity labeling techniques outlined in the "Proximity labeling in brain tissues of human and mice" and "Immunofluorescence of fixed specimens" sections until the stage involving secondary antibodies and streptavidin labeling. The axonal spheroids and biotinylated proteins were labeled using STED-compatible secondary antibody Atto594 (Sigma-Aldrich) (1:100 dilution) and Atto647N-streptavidin (Sigma-Aldrich) (1:100 dilution). Subsequently, the tissues were mounted with Prolong Gold (ThermoFisher) following the user manual, and left in the dark at room temperature for 24 to 72 hours before imaging. STED imaging was carried out utilizing a Leica SP8 STED 3× equipped with a pulsed white light laser (SuperK Extreme EXW-12; NKT Photonics) for excitation and a 775 nm pulsed laser for depletion (Onefive Katana-08HP). The alignment of excitation and STED beams was accomplished using 200 nm Crimson Fluospheres (ThermoFisher; F8782). Sample imaging was performed using the following parameters: Sequence 1: 594 nm laser was at 1.15 uW. Sequence 2: 646 nm laser was at 6.4 uW. Sequence 3: 646 nm laser was at 34 uW. Sequence 4: 594 nm laser was 14 uW. Sequences 3 and 4: 775 nm STED laser was at 33 mW.

### Samples for Focus Ion Beam/Scanning Electron Microscopy (FIBSEM)

Mice brains or fixed postmortem human brains were vibratome-sectioned into 50 μm thickness as described in the ‘proximity labeling in brain tissue’ section [4]. Trimmed tissue samples were fixed in 2.5% glutaraldehyde and 2% paraformaldehyde in 0.1M sodium cacodylate buffer pH 7.4 containing 2 mM calcium chloride for 1 hour, then rinsed in buffer and post fixed in 1% osmium tetroxide and 1.5% potassium ferrocyanide for another hour. After rinsing well, the samples were immersed in aqueous 1% thiocarbohydrazide for 30 minutes and then well rinsed. They were then placed in 1% osmium tetroxide in water for 1 hour at room temperature followed by rinsing in distilled water. An overnight en-bloc stain in aqueous 1% uranyl acetate was followed by rinsing in distilled water and then placed into warm lead aspartate and kept at 600 °C for 1 hour. After 1 hour of rinsing in distilled water, the samples were dehydrated through an ethanol series to 100%, followed by 100% propylene oxide. The samples were infiltrated with Durcupan (Electron Microscopy Sciences) resin over 2 days, then placed in silicone molds and baked at 600 °C for at least 48 hours. The resin block was trimmed to a rough area of interest and the surface was cleanly cut. The entire pyramid was carefully removed with a fine blade and mounted on an aluminum stub using conductive carbon adhesive and silver paint (Electron Microscopy Sciences, Hatfield, PA, U.S.A,) then sputtered with approximately 15 nm Platinum/Palladium (80/20) using a Cressington HR sputter coating equipment (Ted Pella Inc. Redding CA) to reduce charging effects.

A dual beam FIBSEM (Zeiss CrossBeam 550) using a Gallium ion source was used to mill and SE2 secondary electron detector was used to image the samples. Smart SEM (Zeiss, Jenna Germany) was used to set up initial parameters and to find the regions of interest by SEM images at 10 kV 45 μm width and 30 μm height. The actual depth was 22 μm with 10nm/pixel and 10 nm per slice. A platinum protective layer was deposited at the ROI with the FIB (30 kV, 3 nA) to protect the structure and reduce charging. Milling and highlighting were done at 30 kV, 50 pA, with a carbon deposit (30kV 3nA). A course trench was milled (30 kv 30 nA) followed by fine milling (30 kV 3nA) and for final acquisition, a cuboid the area of interest was milled at 30 kV and 300 pA. After milling each slice an image was taken by detecting backscattered electrons of a primary electron beam (acceleration voltage of 1.5 kV, imaging current of 2 nA, and aperture diameter of 100 µm) with a pixel dwell time of 3µs Atlas5 (Zeiss) was used for preliminary SEM stack alignment and FIB/SEM image stacks were saved as Tiff and MRC files. The images were imported into Dragonfly software (ORS, Montreal Canada) for further alignment, segmentation and 3D video.

### FIBSEM image annotation and segmentation

Images from the Z-stack at different Z-locations were imported into APEER (ZEISS). Spheroids and plaques within the images were then manually annotated. After annotating 37 images, models were trained to predict each object. Then, individual spheroid and plaque segmentation models were downloaded and imported into Vision4D 4.1.0 (Arivis) for object segmentation. The entire Z-stack was then segmented in the analysis pipeline via the Deep Learning Segmenter operation with all the Z-stacks within the dataset. Next, non-specific and overlapping annotations were manually corrected using the “Draw Objects Tool.” Object color and opacity were modified using the “Set Object Style” tab to show raw data. The Z-stack was then played using the Movie Player and recorded using Capture.

### Image analysis and quantification

All analyses were processed with FIJI (ImageJ) software, unless otherwise described.

1) Quantification of PLD3 raw integrated fluorescence intensity in human brain: following PLD3 immunolabeling, we compared the number and intensity of pixels corresponding to the PAAS halo versus the rest of the field of view, representing mostly neuronal soma and neuropil. For this, we used three AD postmortem brains which had been used for PAAS proteomics. Immunofluorescence staining of PLD3 and thioflavin S staining was performed followed by confocal imaging. Background autofluorescence signals were measured in unstained brain slices and subtracted from all subsequent image quantifications to better reflect true PLD3 signal. PAAS halos were manually circled, and the raw integrated intensity (RawInt) was measured. The RawInt was also measured for the whole field of view. The sum of RawInt derived from PAAS halos was subtracted from the RawInt from the whole field of view. By subtracting the halo RawInt from the total RawInt, we obtained a measurement of the PLD3 labeling outside of plaques which mainly consists of neuronal cell bodies, neuropil and any minimal background fluorescence.
2) p-mTOR S2448 fluorescence intensity measurement: in human AD or no/mild AD postmortem brains, each zoom 1 z-stack image, containing SMI312 positive axonal spheroids around amyloid plaques, were maximum projected. Then, SMI312 positive spheroid halos were circled, and mean fluorescence intensity of the p-mTOR S2448 channel was measured within the selected circle. In the same field of view, regions without spheroids were considered as background. Three such background regions were selected, circled and p-mTOR mean intensity was measured. The three “background” p-mTOR mean intensity from each field of view was averaged. Mean intensity (p-mTOR dystrophy halo) / mean intensity (p-mTOR background) was calculated to represent the p-mTOR expression level in each axonal spheroid halo. Three fields of view were quantified in each severe AD patient, and 1-3 fields of view were quantified in each no/mild AD patient. The averaged p-mTOR expression level in each patient was used for statistical analysis.
3) Measurement of the size of individual axon spheroids in human iPSC-derived neurons in the Torin1 treatment experiment: SMI312 was used to label spheroids, while ThioflavinS was used to label amyloid plaques. Tiling images were taken in each well from a 96-well plate and were maximum projected. For Torin1 treatment before amyloid plaque and spheroid formation, individual axonal spheroids were circled using the freehand tool on ImageJ, and the circle area was measured. The total number of axonal spheroid halos analyzed ranged from 50-150 within each tilling image. Two technical replicate wells were analyzed in each experiment and three batches of experiments were performed for quantification. For Torin1 treatment after amyloid plaque and spheroid formation, machine learning based image analysis software Aivia v12 (Leica) was used to quantify the size and number of spheroids and amyloid plaques, as well as axon density in an automated fashion. The pixel classifier tool was used to classify each channel.
4) Measurement of the percentage of axonal segments with spheroids in human iPSC-derived neurons in the Torin1 treatment experiment: The SMI312 staining revealed both axonal spheroids (the spheroid shape) and axons (the linear shape), therefore allowed us to recognize individual axon segments. To quantify axons with spheroids, we traced individual axon segments with linear and continuous signal across the z-stacks within the field of view. An axon with one or more spheroids formed was defined as axon with spheroid, whereas axon segments without spheroids observed within the z-stack, were defined as axons without spheroids. In Figure 6L, the percentage of axons with spheroids was calculated by comparing the number of axonal segment with spheroids over the total number of axonal segments that were observed within the field of view.
5) Measurement of human neuron cell body size with or without Torin1 treatment: iPSC co-cultures were infected by AAV2-CB7-GFP (Addgene#105542) at titer ∼7x10^9 vg/mL one week before Torin1 and amyloid beta treatment. Cells were stained with NeuN and GFP antibodies. Large field tilling imaging was performed in each well. Tilling images were maximum projected and neurons with both NeuN and GFP positive signals were measured by size, using the freehand tool in ImageJ. Three technical replicate wells were analyzed in each experiment and two batches of experiments were done for quantification.
6) Individual axonal spheroid size measurement in the AAV9-hSyn-cre-2a-tdT infected *Mtor*-flox-5XFAD mice: anti-Lamp1 antibody was used for immunofluorescence staining to showcase the spheroid halo around amyloid plaques, ThioflavinS was used to label amyloid plaques, and anti-RFP antibody was used to reveal the tdTomato expression in infected neurons. Tilling images were taken at the somatosensory cortex of each animal. Three serial brain sections were used for tilling imaging for each mouse. An individual spheroid was counted when it was both tdTomato and Lamp1 positive. The individual spheroid size was measured using the freehand tool in ImageJ to circle the outline of spheroids, where it had the maximum diameter, followed by measurement of the circled area. The total number of axonal spheroid halo analyzed ranged from 50-150 in each tilling image.
7) Measurement of spheroid halo size in the AAV-PHPeB infected *Mtor*-flox-5XFAD experiment: anti-Lamp1 antibody was used for immunofluorescence staining to showcase the spheroid halo around amyloid plaque, ThioflavinS was used to label amyloid plaques. Tilling images were taken at the somatosensory cortex of each animal. Three serial brain sections were used for tilling imaging for each mouse. A customized ImageJ macro was used to segment individual Lamp1 positive spheroid halos and amyloid plaques. Using a customized MATLAB program, the segmented images were processed to measure the area of individual spheroid halos and amyloid plaque area, automatically. The spheroid halo area was excluding the area in the center occupied by the amyloid plaque. The number of axonal spheroid halos and amyloid plaques analyzed ranged from 30 to 70 in each tilling image from the *Mtor*-flox-5XFAD experiment.
8) Measurement of RNA signals within axonal spheroids: ImageJ free hand tool was used to circle axonal spheroid structures labeled by Atto-647 NHS ester, and fluorescence intensity from PolyA probe channel, or negative control probe channel was measured. For measuring puromycin signals within axonal spheroids or neuronal cell bodies, similarly, ImageJ free hand tool was used to circle axonal spheroid or cell bodies labeled by Lamp1 or NeuN, respectively, and fluorescence intensity from puromycin channel was measured.
9) Measurement of NeuN positive neuronal cell soma area in AAV-PHPeB infected *Mtor*-flox-5XFAD mice: A customized CellProfiler program was used for automated measurement. Briefly, tilling images were taken from one hemisphere of the somatosensory cortex. Tilling images were maximum projected before importing them into CellProfiler (Broad Institute). The NeuN channel was set as the primary object and was used as the marker for size measurement. Three tilling images were taken from three brain sections of each animal. Three animals were used in each group.
10) Investigation of mTOR heterozygous knockout downstream signaling effectors in mTOR-floxed-5XFAD mice: Similarly, a customized CellProfiler program was used for automated measurement. Tilling images were maximum projected before analysis. To analyze fluorescence intensity in the nuclei, we used DAPI staining as the primary object and measured signals from the NeuN positive cells. To measure fluorescence intensity from the cell bodies, the NeuN channel was set as the primary object and was used as the marker for fluorescence intensity measurement. We compared mice injected with PHPeB-hSyn-cre-GFP viruses, with those injected with control viruses PHPeB-CAG-GFP. The littermates and sex were paired for comparison. Immunofluorescence intensity of TFEB was measured in neuronal nuclei, while LC3B, P-p70S6K Thr389 and p70S6K were measured in neuronal soma in an automated fashion.

### Statistical analysis

No statistical power analysis was used to determine sample sizes, but our sample sizes were similar to those generally employed in the field. All samples were included for analysis. Excel (Microsoft), Prism (GraphPad), Qlucore Omics Explorer v3.6 (Qlucore AB, Lund, Sweden), CellProfiler v4.2.1 (Broad Institute), MATLAB and RStudio (4.0.2) were used for data analysis and plotting. Statistical methods used were described in the figure legend of each relevant panel.

## DATA AND CODE AVAILABILITY

Raw proteomics data is provided in **Table S1**. The mass spectrometry proteomics data have been deposited to the ProteomeXchange Consortium via the PRIDE partner repository with the dataset identifier xxx. Reviewer account details: Username: xxx, Password: xxx). The Lamp1 dataset is under the identifier xxx, username: xxx, password: xxx. For sample information see **Table S3**.

Custom codes for FIJI and MATLAB were deposited at GitHub (https://github.com/PaulYJ/Axon-spheroid) as we previously described [4].

The Python code for the analysis of calcium imaging data can be accessed at the following location: https://github.com/ShawnQin/calcium_trace.

The code for the analysis of STED imaging data can be accessed at the following location: https://github.com/bewersdorflab/Yifei-Lukas-Collab.

## Acknowledgments

This project was supported by National Institute of Health grants: RF1AG058257, R01NS115544, R01NS111961 (J.G.), a Cure Alzheimer’s Fund Research Grant (J.G.), a Yale/NIDA Neuroproteomics Center Pilot Project Grant 2019 (Y.C.), a BrightFocus Foundation Postdoctoral Fellowship Program in Alzheimer’s disease Research (A2021003F) (Y.C.), a Yale ADRC Research Scholar Award (Y.C.), an Alzheimer’s Association Research Fellowship (23AARF-1020552) (Y.C.), a Yale ADRC grant P30 AG066508 (A.C.N), and the following grants R01AG068030, RF1AG065926, R56AG071291 to K.J.B . We thank Drs. Kenneth Williams, Shannon Leslie, Bobby Mathew and the Yale/NIDA Neuroproteomics Center (P30 DA018343), for providing experimental design advice, technical support and funding opportunities. We thank staff from the Keck MS & Proteomics Resource at the Yale School of Medicine for processing the LC MS/MS experiments. We also thank the Keck MS & Proteomics Resource at the Yale School of Medicine for providing the necessary mass spectrometers and the accompany biotechnology tools funded in part by the Yale School of Medicine and by the Office of The Director, National Institutes of Health (S10OD02365101A1, S10OD019967, and S10OD018034). The funders had no role in study design, data collection and analysis, decision to publish, or preparation of the manuscript. I.P.S and E.P. were supported by EMBL-EBI Core funding. We thank Dr. Anita Huttner and the Yale ADRC (P30 AG066508) for providing human postmortem AD and control brain tissues. Additional MCI and AD brain tissue was provided by the Banner Sun Health Research Institute Brain and Body Donation Program of Sun City, Arizona, supported by the NINDS (U24 NS072026), the NIA (P30 AG19610), the Arizona Department of Health Services (contract 211002), and the Arizona Biomedical Research Commission (contracts 4001, 0011, 05-901 and 1001). We thank Drs. Nur-Taz Rahman and Rolando Garcia Milian from the Yale Medical Library Bioinformatics Support Program for providing advice and technical support for bioinformatics analysis. We thank Dr. Xinran Liu, Ms. Morven Graham and the Center for Cellular and Molecular Imaging Electron Microscopy Facility at Yale Medical School for assistance with the electron microscopy experiments (NSF Major Instrument grant 1725480). We thank Drs. Peng Yuan, Lei Tong and Mengyang Zhang for providing MATLAB and FIJI imageJ coding for plaque and axonal spheroid quantification. We thank Dr. Mengyang Zhang for generating the AAV2-CMV-LAMP-GFP viruses. We thank Drs. Jelena Platisa, Peter O’Brien, Vincent Pieribone, Ilona Kondratiuk, Dhrubajyoti Chowdhury, Thomas Biederer, Shengnan Qiao, Yao Xue and Jimmy Zhou for their generous help and critical advice on the calcium imaging experiments. Calcium imaging was performed using dragonfly spinning disk confocal from the Biederer lab, supported by grant NIH R01 DA018928 (to T.B.). We thank Drs. Teresa Spano and Erin Schuman for their generous help and critical advice on the *in vivo* puromycylation experiments. We thank Dr. Philip Coish for the critical reading of this manuscript. We created schematic figures with BioRender.com.

## Author contributions

Y.C. and J.G. conceived and designed the study. Y.C. performed proximity labeling experiments, Y.C., R.W., M.S.M., T.T.L., S. B. and A.C.N. designed protein extraction, enrichment and digestion protocol, Y.C. performed protein extraction, enrichment and western blotting, J.K., R.W., S.B. and T.T.L. performed LC-MS-MS, protein mapping and searching, Y.C. performed proteomics data analysis, Y.C., I.P.S and E.P performed bioinformatics analysis, A.C.N. supervised the proteomics experiments and analysis; L.A.F. and Y.C. performed STED imaging, L.A.F., T.H. and Y.C. performed STED imaging analysis; Y.C., L.T., and L.S. performed calcium imaging in iPSC model, L.T., S.Q. performed calcium imaging data analysis; A.H. performed pathological evaluation of human brain specimens and provided the tissues; Y.C. and T.H. performed immunofluorescence and confocal imaging; T.H. performed immunofluorescence quantification of newly identified proteomic hits and automated tilling images analysis; Y.C., P.L.C. and K.J.B. established the human iPSC-derived neuron-astrocyte co-culture protocol, P.L.C. performed lentivirus packaging, Y.C. performed AD human neuron-astrocyte co-culture, K.J.B. supervised the human neuron-astrocyte co-culture experiment; Y.C. performed mTOR inhibition in transgenic mice, Y.C., T.H., Z.T., A.B. and K.T. performed quantifications, Y.C. and T.H. performed statistics analysis; H. G. annotated FIBSEM video; Y.C. and J.G. prepared the manuscript. J.G. supervised the study.

## SUPPLEMENTARY DATA

**Figure S1.**
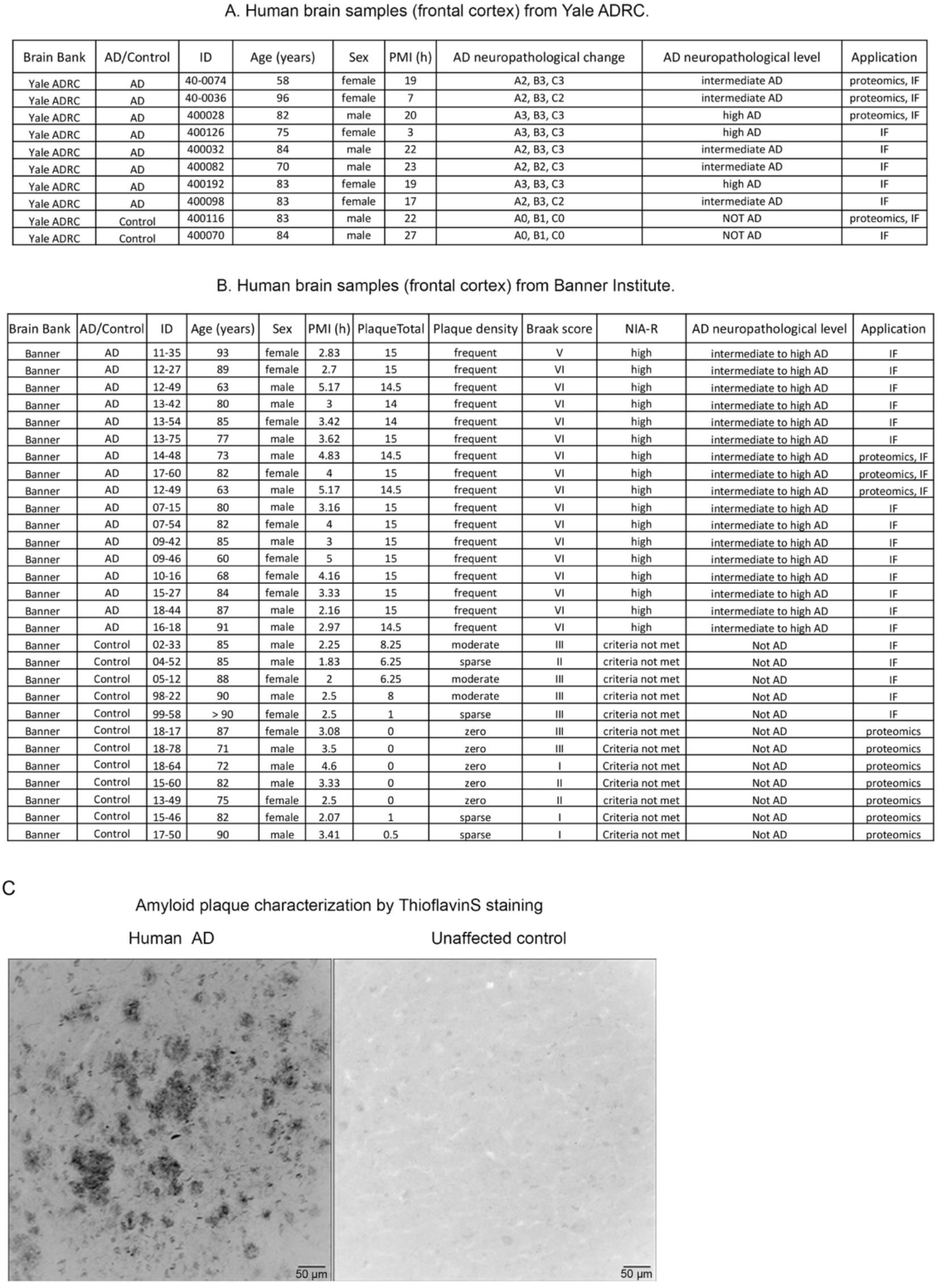
Metadata associated with the postmortem human brains. **A-B.** Specimens obtained from: **(A)** the Yale Alzheimer’s Research Center and (**B**) the Banner Sun Health Research Institute. IF = immunofluorescence validation. **C.** Representative images of a large field of view showing amyloid plaques (dark, ThioflavinS) in postmortem brains of AD humans and unaffected controls. Scale bar 50 μm.

**Figure S2.**
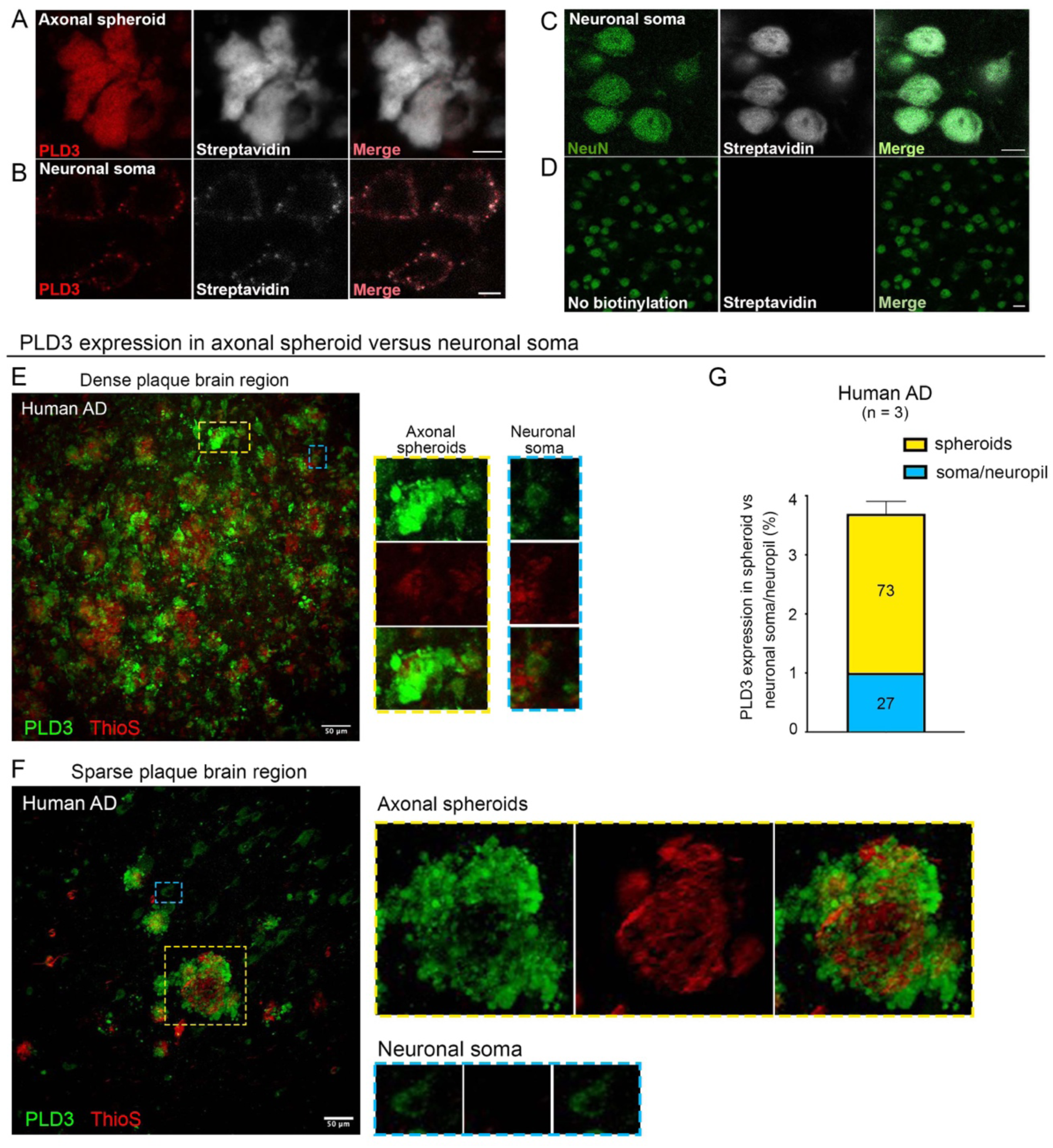
PLD3 expression and proximity labeling of axonal spheroids versus neuronal somata. **A-B.** Proximity labeling of PLD3 (red) in (**A**) axonal spheroids versus (**B**) neuronal somata. C. Proximity labeling of NeuN (green, a neuronal soma marker). (**A-C**) Scale bar 5 μm. **D.** No biotinylation reaction control shows that the streptavidin signal was eliminated. Scale bar 50 μm. **E-G.** Comparison of PLD3 expression in axonal spheroids versus neuronal soma. Representative images of PLD3 immunofluorescence staining in AD postmortem brains with (**E**) dense amyloid plaques (red, ThioflavinS stained) and (**F**) sparse amyloid plaques. Inserts show axonal spheroid halos around amyloid plaques (yellow squares) or neuronal soma (blue squares). Scale bar 50 μm. **G.** Quantification of PLD3 expression in axonal spheroids versus neuronal soma/neuropil in AD human postmortem brains (n = 3 brains).

**Figure S3.**
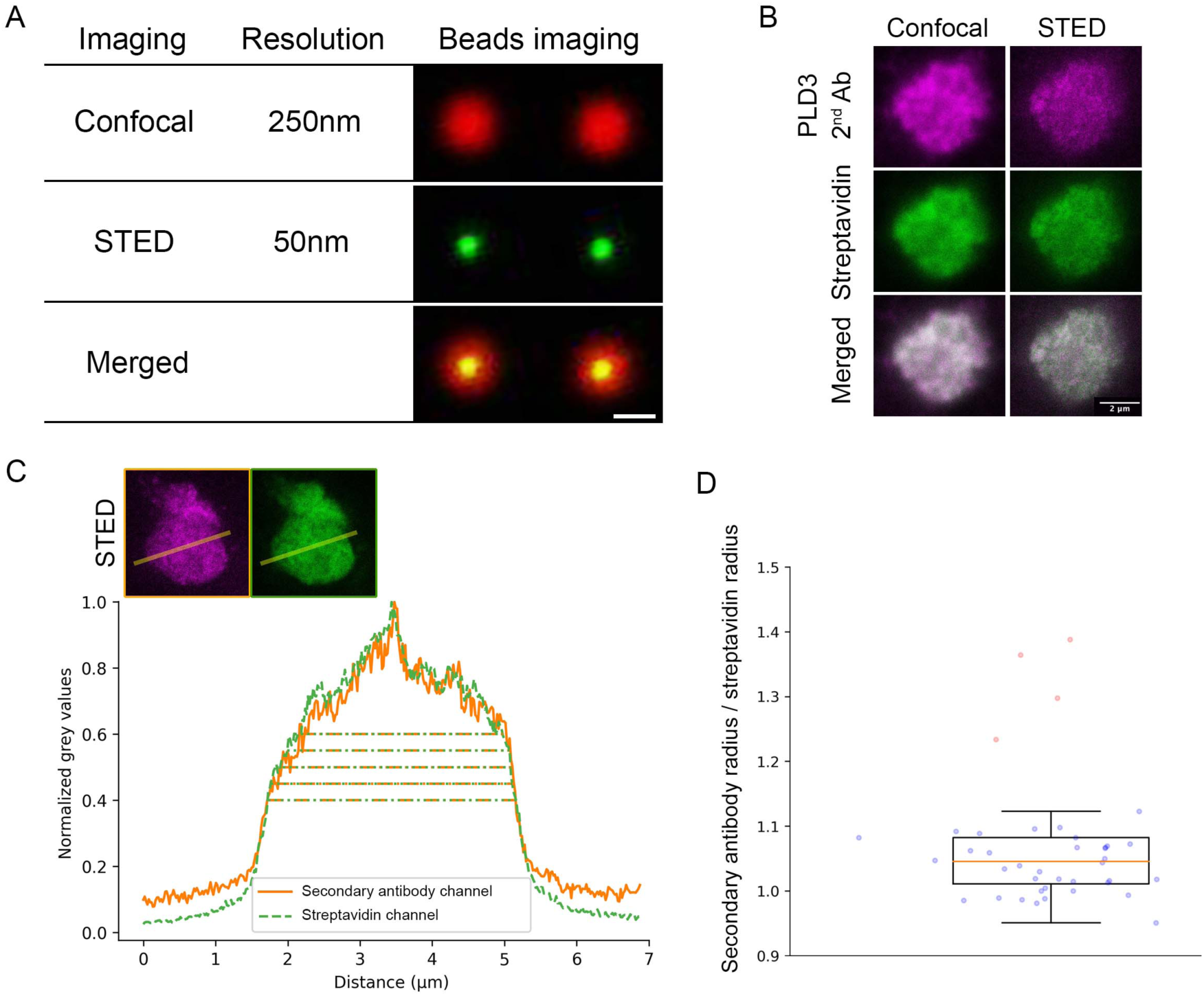
Super-resolution STED imaging reveals high spatial precision of proximity labeling in AD human brain. **A.** Imaging of beads illustrates the resolution contrast between confocal microscopy (250 nm) and STED microscopy (50 nm). Scale bar = 250 nm. **B-C.** Representative confocal and STED images showing proximity labeling of axonal spheroids (magenta, anti-PLD3 labeled) in AD human postmortem brains. Biotinylated proteins were labeled by streptavidin (green). Scale bar = 2 μm. **B.** A line plot representative of the radius measurements illustrates the signals from both the secondary antibody channel (magenta) and the streptavidin channel (green). **D.** Dot plot depicting the radius ratio between the secondary antibody channel (magenta) and the streptavidin channel (green). Average radius ratio = 141.0 nm, average ratio = 1.04, standard deviation = 0.04. The median value is represented by the orange line, while outliers are denoted by pink circles. Any value surpassing 1.5 times the interquartile range (IQR) above Q3 or below Q1 is classified as an outlier and is subsequently excluded [115].

**Figure S4.**
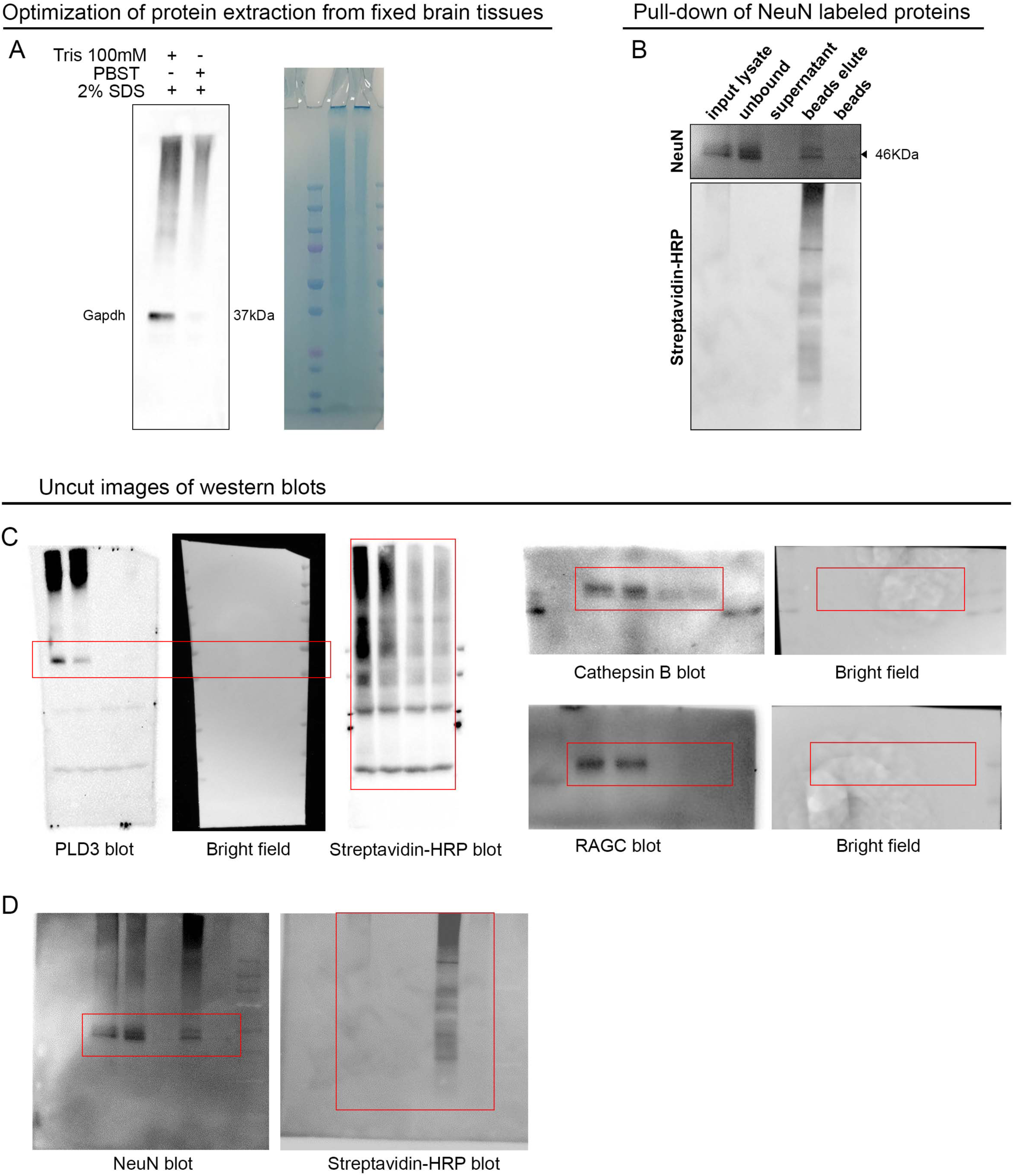
Optimization of protein extraction and western blots. **A.** Optimization of protein lysis protocol for fixed postmortem brains. The protein lysis buffer (100 mM Tris-HCL plus 2% SDS) extracted proteins much more efficiently compared to the previously published method (PBST plus 2% SDS), as indicated by a blot of housekeeping protein GAPDH and the Coomassie blue gel of the same membrane. **B.** Western blot showing biotinylated proteins pulled down from an anti-NeuN antibody proximity labeled sample. Biotinylated proteins, including the protein bait NeuN, were detected in the input brain lysates and streptavidin bead elutes but not in the supernatant from bead washes nor in the beads themselves after elution. Although the bait proteins were also detected in the unbound protein fractions, no streptavidin signal was detected from the unbound proteins, indicating that there was a portion of endogenous protein baits that was not biotinylated, yet most of the biotinylated proteins were pulled down efficiently. **C-D.** Uncropped western blot membranes of **(C)** PLD3, Cathepsin B and RAGC pulled down, related to Figure 1I, and **(D)** NeuN pulled down, related to **Figure S4B**.

**Figure S5.**
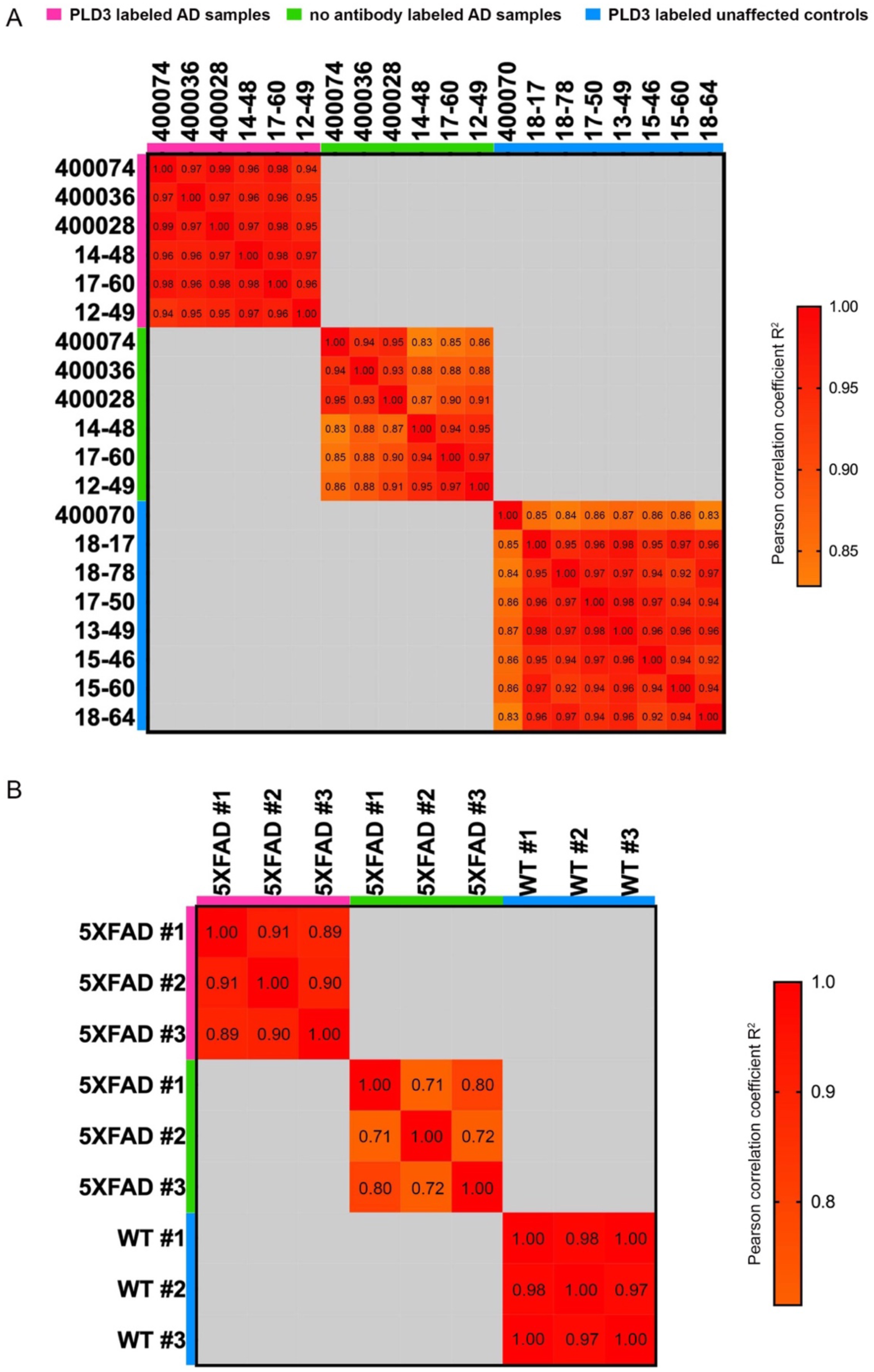
Correlation analysis of proteomics samples in humans and mice. Correlation analysis among biological replicates of PLD3-labeled and no antibody-labeled proteomic samples in (**A**) humans and (**B**) mice. Pearson correlation coefficient R2 of each comparison is listed in each box.

**Figure S6.**
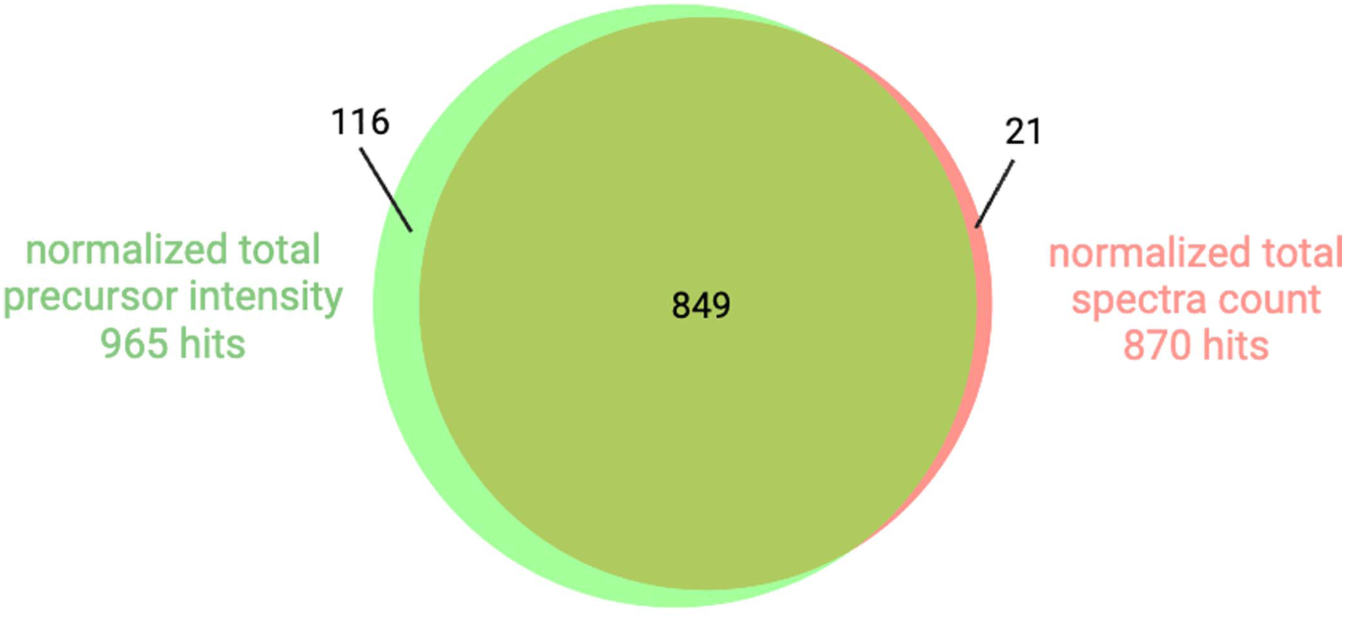
Quantification approach comparison in human PAAS proteomics. Human PAAS proteomics analysis was performed using either normalized total precursor intensity approach or normalized total spectra count approach. Venn diagram showed the comparison of 965 proteomics hits and 870 proteomic hits identified by these two approaches respectively.

**Figure S7.**
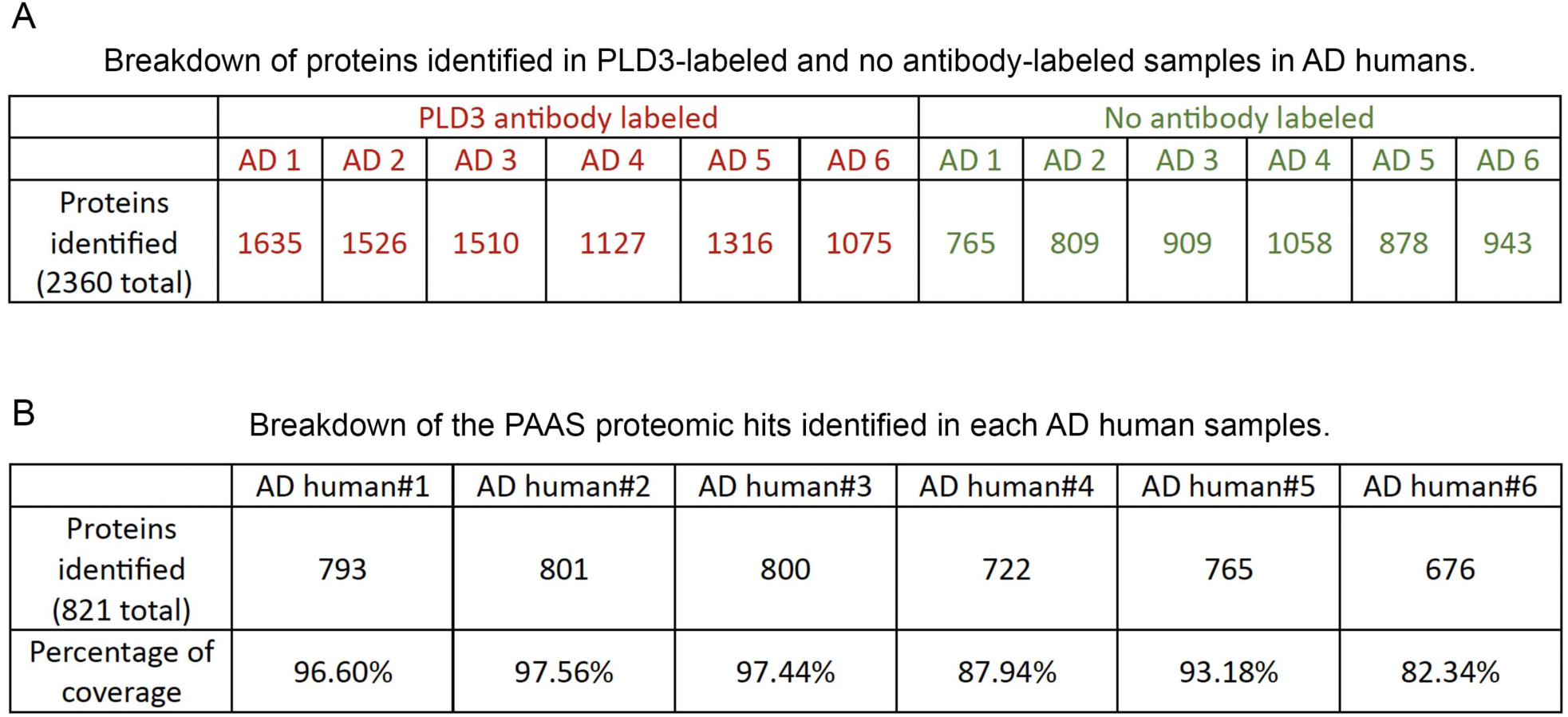
Breakdown of protein numbers identified in each human sample. **A.** Table shows all the proteins identified in the PLD3-labeled and no antibody-labeled samples in each AD humans. **B.** Table shows the identified PAAS proteomic hits number in each AD humans.

**Figure S8.**
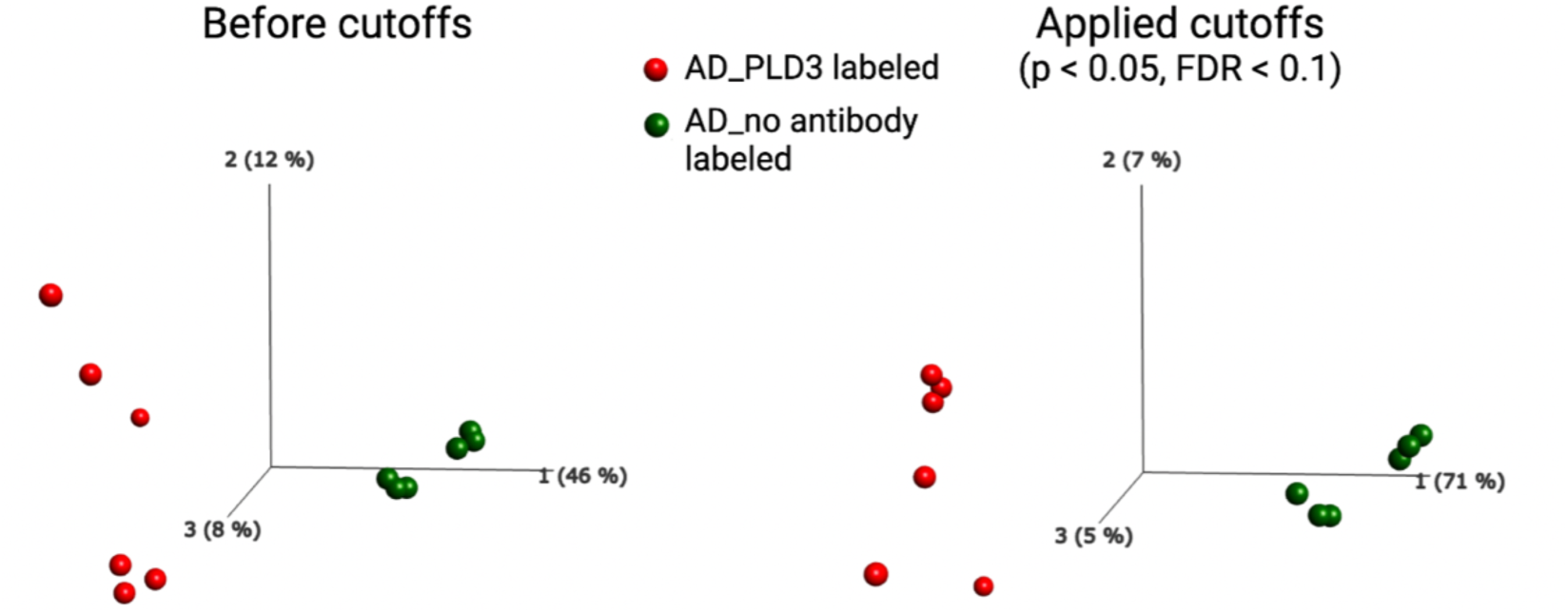
Principal component analysis (PCA) of PLD3-labeled versus no antibody-labeled proteomes in AD human postmortem brains. PCA plots showing identification of proteins and their quantities were found to be dependent on the presence of the antibody. Even without applying any cutoffs, the PLD3 antibody-labeled samples exhibited clear differentiation from the no antibody-labeled samples. Furthermore, applying cutoffs (p < 0.05 and FDR < 0.1) resulted in an increase in PC1 from 46% to 71%.

**Figure S9.**
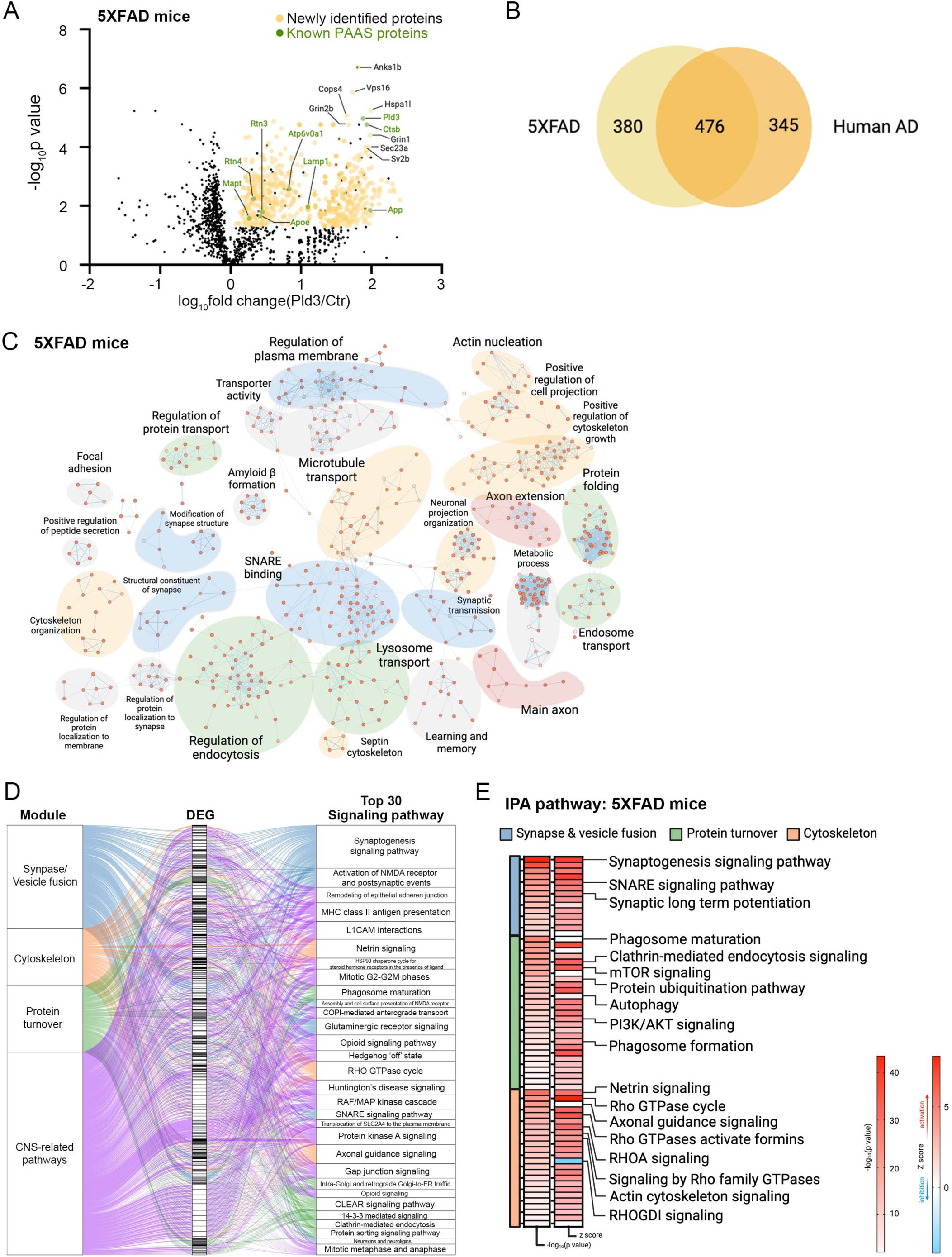
Proteomics analysis of PLD3-labeled PAAS proteomes in 5XFAD mice. **A.** Volcano plot show proteins (represented by their gene names) that passed the statistical cutoffs (yellow dots) in 5XFAD mice. The top 10 proteomic hits with the lowest p-value and highest fold changes are indicated by their gene names in black. The selected known PAAS proteins are labeled as green dots and their gene names in green. The black dots among the yellow ones represent proteins being filtered by the statistical cutoff as shown in Figure 2B. (See **Table S1** for full list of proteomic hits). **B.** Venn diagram shows shared proteomics hits between AD humans and mice. **C.** Pathway enrichment analysis of PAAS proteome in 5XFAD mice. The Enrichment Map represents a network of pathways where edges connect pathways with many shared genes. Node color reflects the FDR of each pathway. The theme labels were curated based on the main pathways of each subnetwork. Subnetworks with a minimum of four pathways connected by edges are shown. **D**. IPA pathway analysis of the PAAS proteome in 5XFAD mice. Top 30 CNS-related signaling pathways are shown. The signaling pathways are summarized as 4 modules. The alluvium plot shows different modules connect to the differentially expressed genes (DEGs) and the DEGs connects to the pathways that they are involved. **E.** IPA pathways related to the three modules with a p-value less than 0.01 are listed. Heatmaps indicate either the -log10 (p-value) or the z score of each signaling pathway (pathways with a z score in red are predicted to be activated while blue ones are predicted to be inhibited).

**Figure S10.**
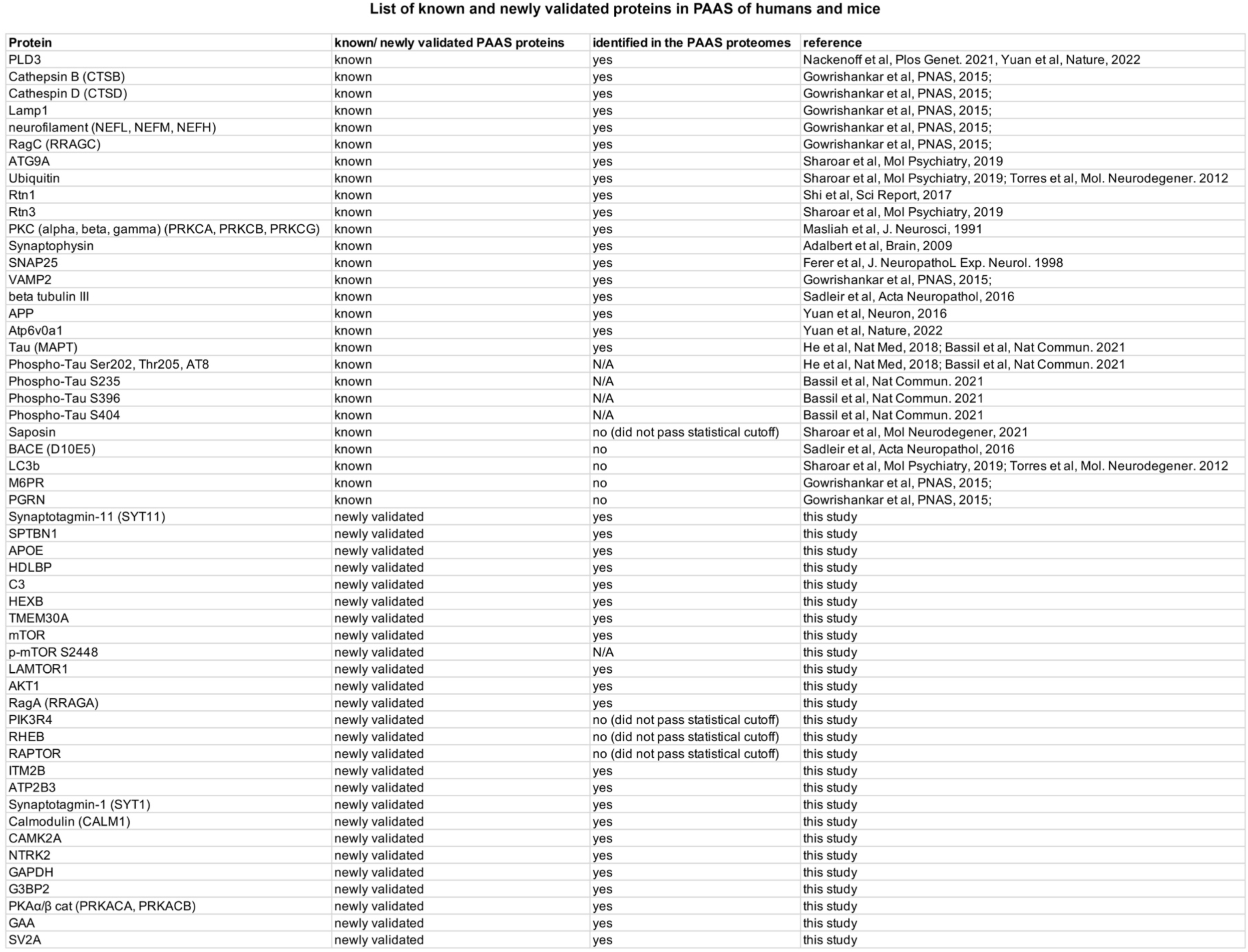
List of known PAAS proteins and newly validated PAAS proteomic hits in human and mice. Full information of this table including the antibodies, immunofluorescence staining information and quantification can be found in **Table S2**.

**Figure S11.**
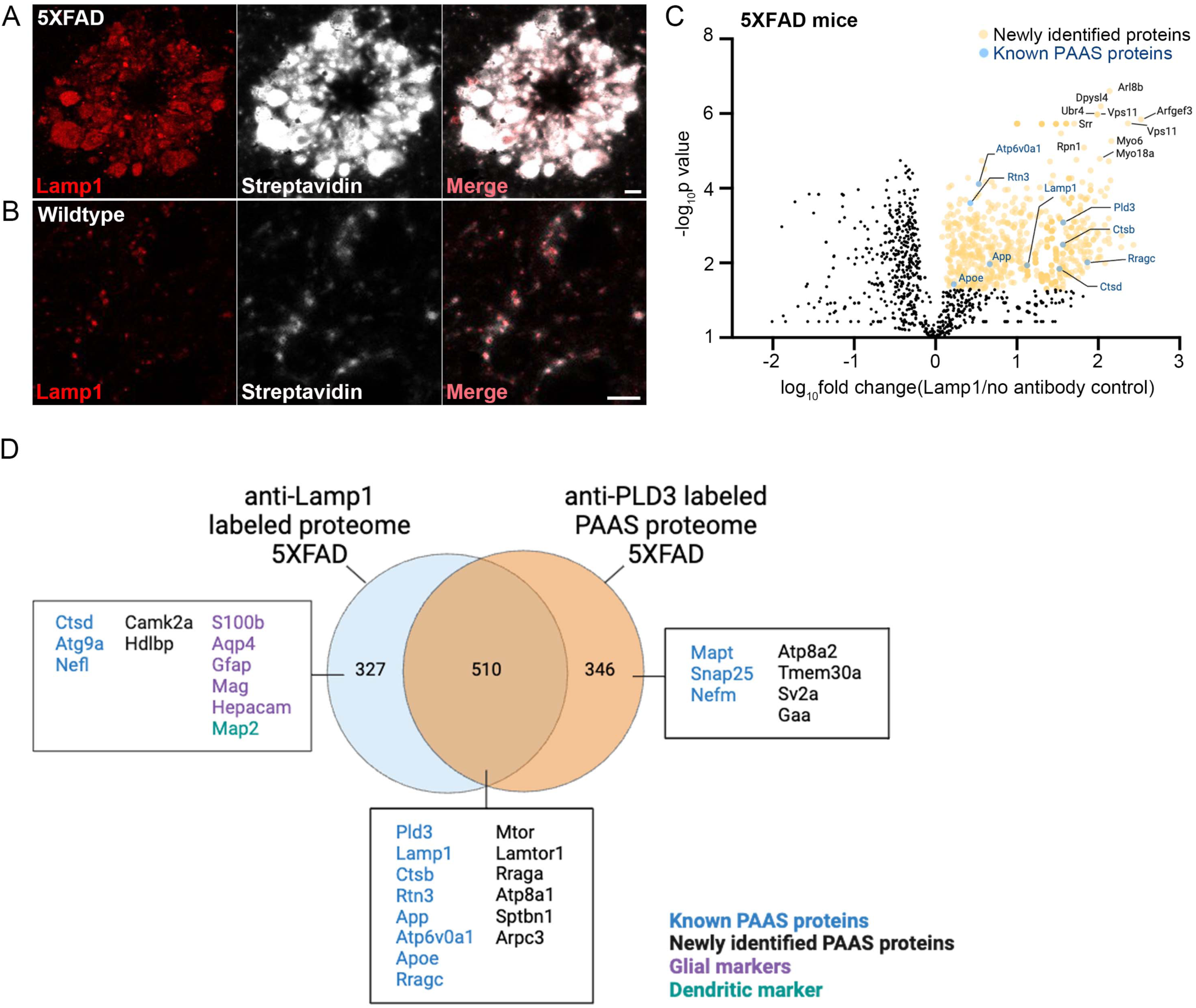
Comparison between anti-Lamp1 antibody labeled proteome with the anti-PLD3 antibody labeled PAAS proteomes in 5XFAD mice. **A-B.** Proximity labeling of Lamp1 (red) in (**A**) 5XFAD and (**B**) wild type mice. Lamp1 (red) labeled (**A**) axonal spheroid halo in 5XFAD mice and (**B**) lysosomes in neuronal cell bodies. Biotinylated proteins were labeled by streptavidin. Scale bar 5 μm. **C.** Volcano plot shows proteins that passed the statistical cutoffs (yellow dots) in 5XFAD mice. The gene names of the top 10 proteomic hits with the lowest p-value and highest fold changes are shown in black. The known PAAS proteins are shown as dark blue dots with their gene names labeled in blue. (See **Table S1** for full list of proteomic hits). **D.** Venn diagram showing comparison of the anti-Lamp1 antibody labeled proteomes with the anti-PLD3 antibody labeled PAAS proteomes in 5XFAD mice. Selected known PAAS proteins are shown in blue, newly identified proteins are shown in black, glial marker proteins are shown in purple and dendritic marker protein is shown in green.

**Figure S12.**
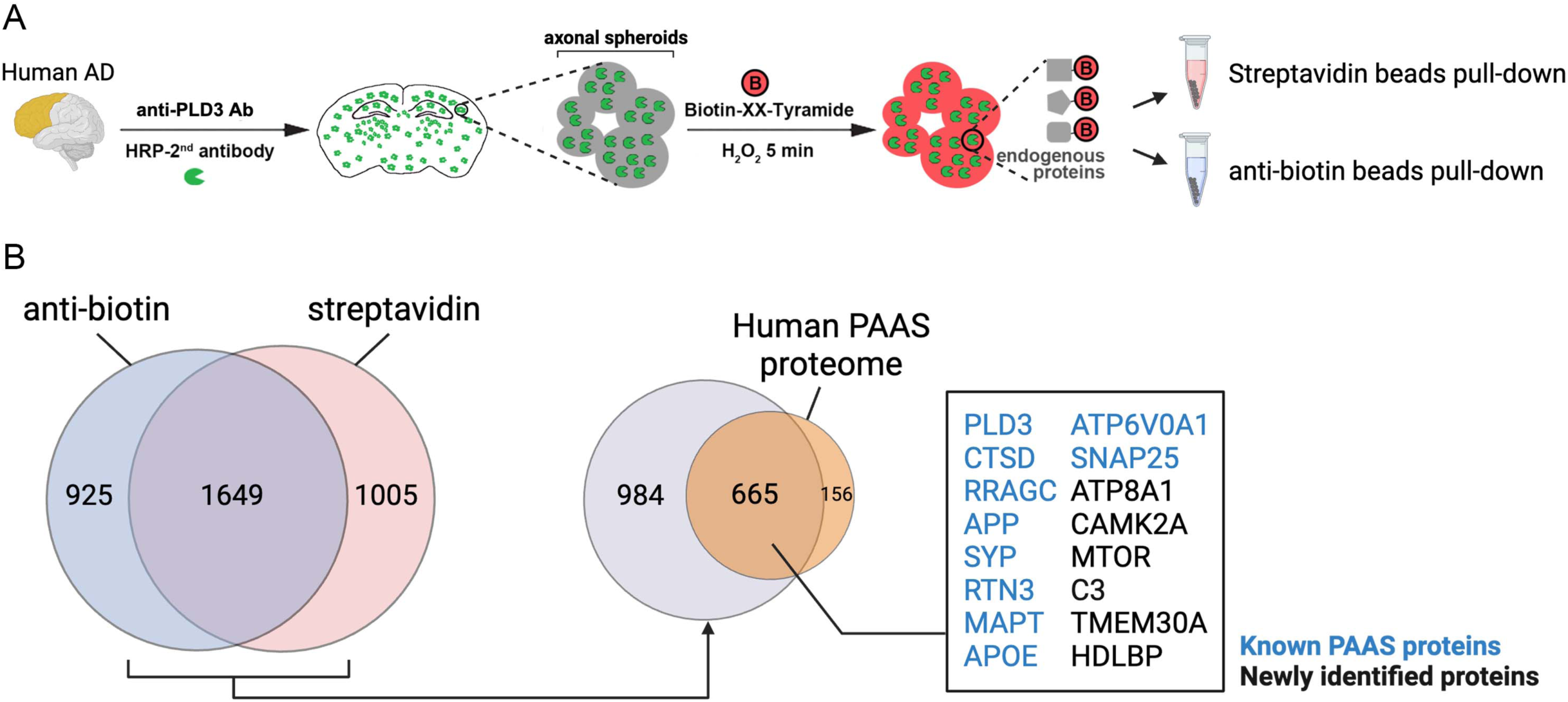
Comparison between anti-biotin beads pulldown with streptavidin beads pulldown of anti-PLD3 antibody labeled proteomes in AD human brains. **A.** Schematic showing pipelines for using streptavidin beads or anti-biotin beads to pulldown anti-PLD3 antibody labeled proteomes in AD human brains. For streptavidin beads, tissue sections were lysed and then incubated with streptavidin beads. For anti-biotin beads, protein lysate was digested prior to anti-biotin beads pull down. **B.** Comparison of the proteomes captured using anti-biotin beads and those captured with streptavidin beads. Among the total 821proteomic hits identified in the human PAAS proteome, 665 hits were pulldown by both the streptavidin beads method and the anti-biotin beads method. Selected known PAAS proteins (blue) and newly identified proteins (black) are shown in the box.

**Figure S13.**
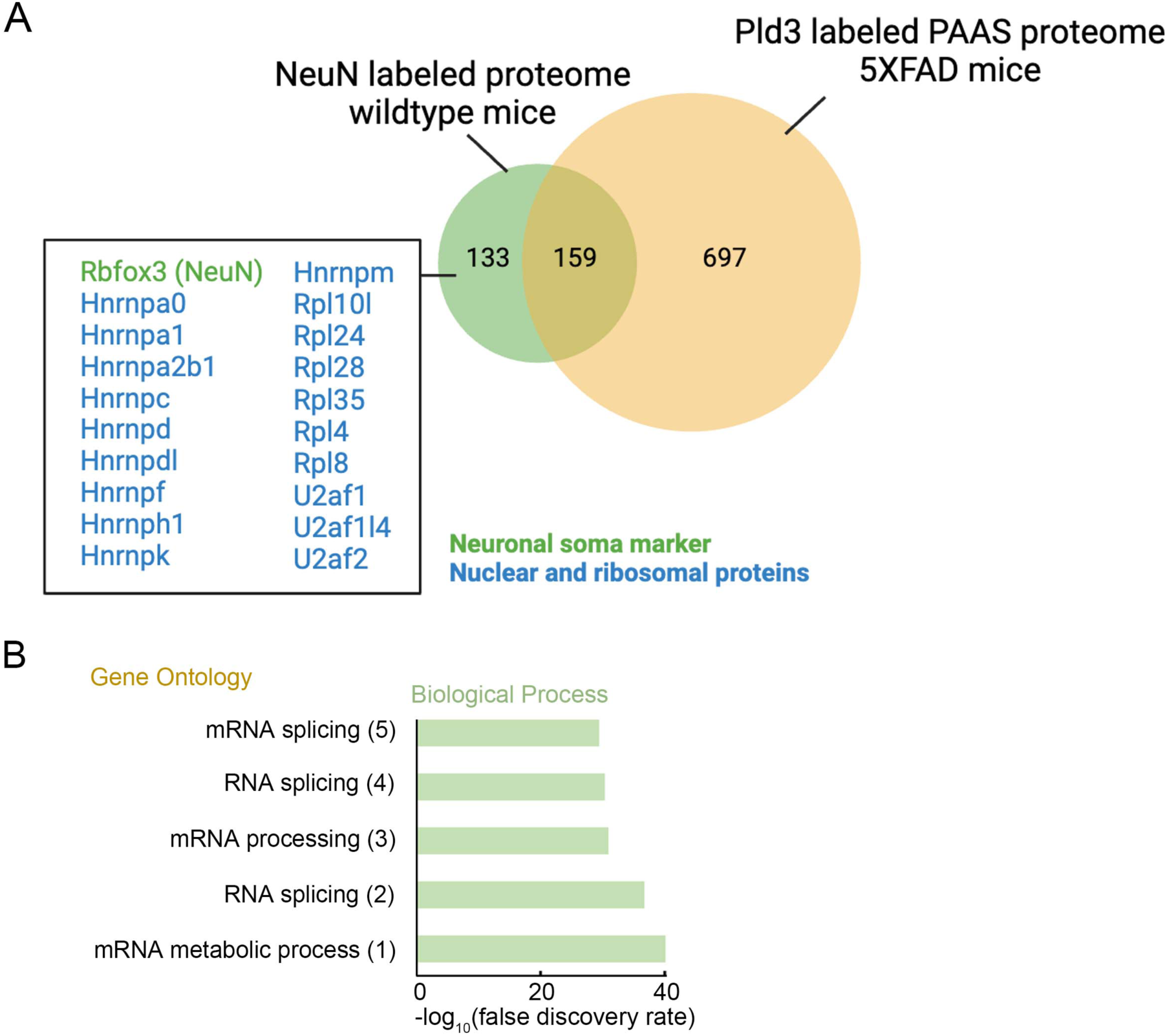
Proteomics analysis of NeuN-labeled neuronal nuclei and perinuclear cytoplasm proteomes in mice. **A.** The NeuN-labeled neuronal nuclei and perinuclear cytoplasm proteomes in wildtype C57BL/6J mice contain 292 proteomic hits. Comparison between the NeuN-labeled proteome and the PLD3-labeled PAAS proteome in mice showed 159 proteins are shared. Among the 133 unique protein hits in the anti-NeuN-labeled proteome, the protein bait and neuronal soma marker NeuN was detected, along with many nuclear and ribosomal proteins. **B** Gene Ontology analysis shows the top ranked biological process terms of the anti-NeuN-labeled proteomic dataset.

**Figure S14.**
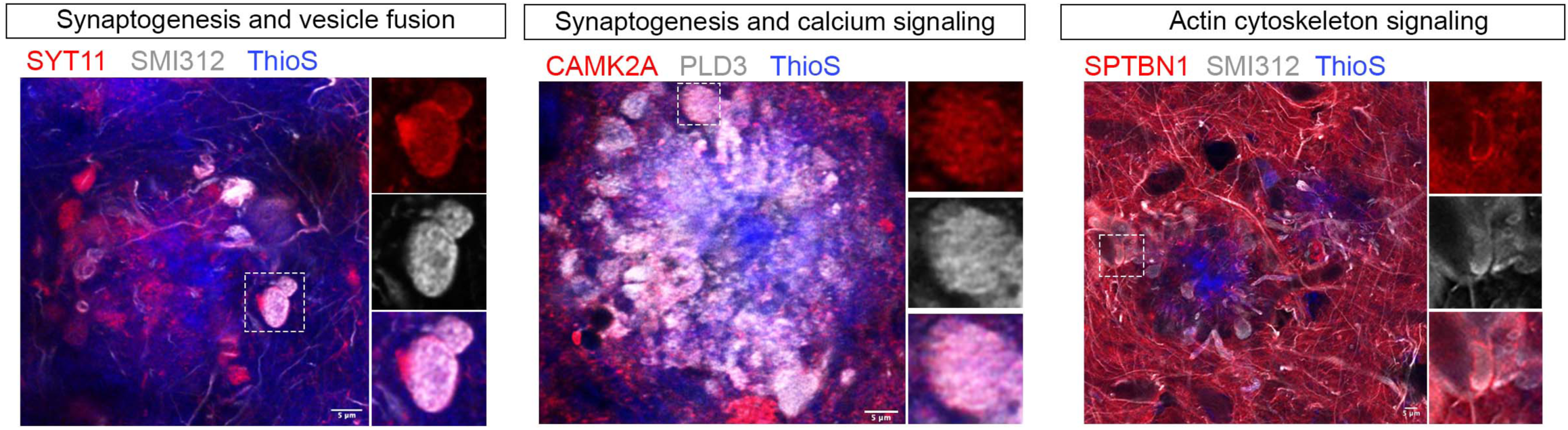
Immunofluorescence confocal imaging validation of new proteins related to synaptogenesis, vesicle fusion, calcium signaling and actin cytoskeleton in axonal spheroid in AD human postmortem brains. Representative zoom-out immunofluorescence confocal images of newly identified proteins expressed in axonal spheroids in AD human postmortem brains (inserts were shown in Figure 3E). Scale bar = 5 μm. Quantification was performed in n = 10 AD human brains. Protein expression quantifications can be found in **Table S2**.

**Figure S15.**
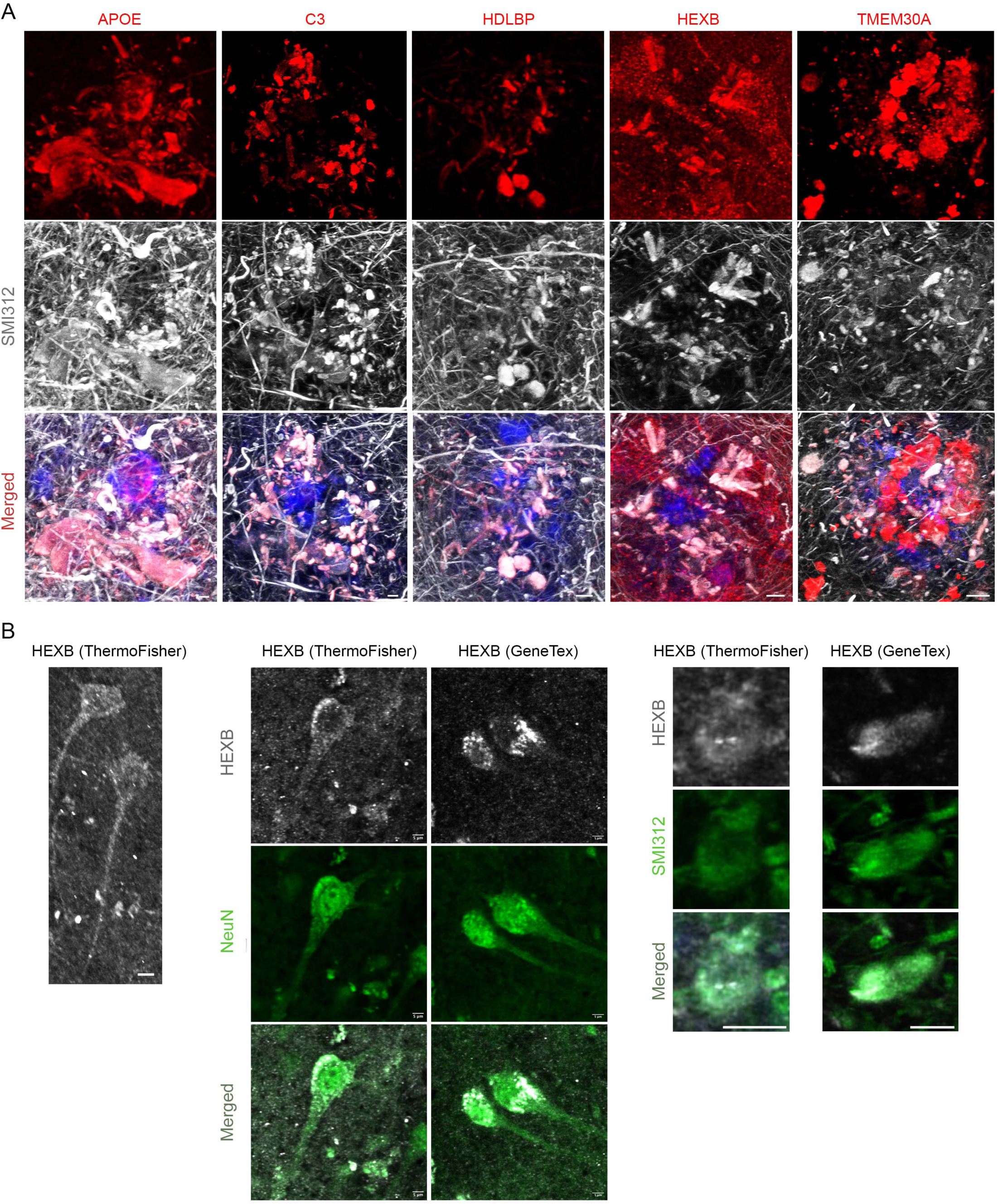
Lipid transport-related proteins are expressed in axonal spheroids around amyloid plaques in AD human postmortem brains. **A.** Representative immunofluorescence confocal images of the top-ranked lipid-related proteomic hits, including APOE, HDLBP, C3, HEXB and TMEM30A in AD humans. Lipid transport-related molecules are shown in red. Neurofilament marker SMI312 (grey) indicates the neuronal branches, and axonal spheroid structures around amyloid plaques (ThioflavinS, blue). Scale bar = 5 μm. Quantification was performed in n = 3 AD human brains. Protein expression quantifications can be found in **Table S2**. **B.** Immunofluorescence confocal image shows HEXB (grey) expressed in neuronal cell bodies (NeuN, green) and axonal spheroids (SMI312, green) in AD human postmortem brain (n = 3). Two different HEXB antibodies were used for validation. Scale bar = 5 μm.

**Figure S16.**
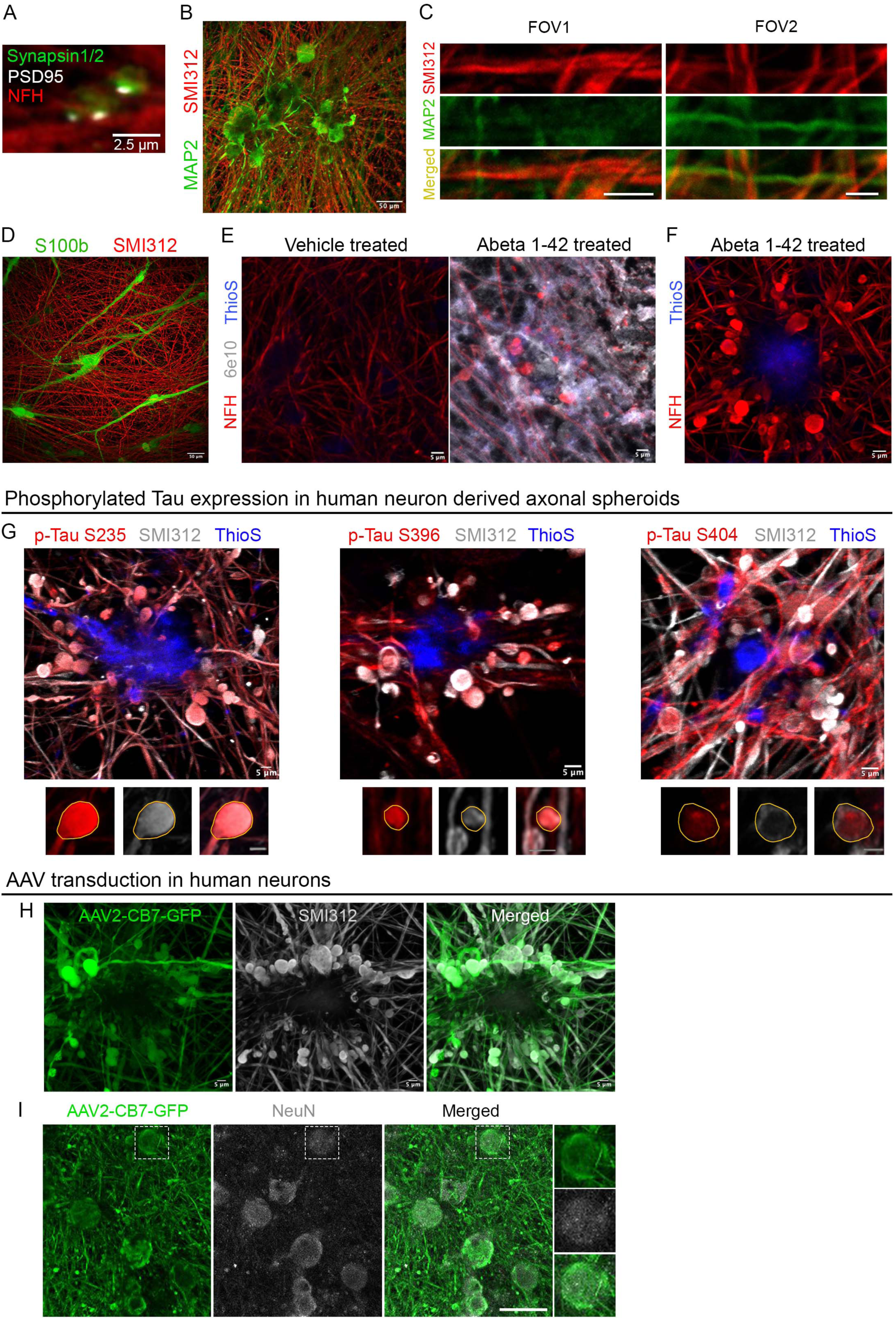
Characterization of the human iPSC-derived neuron-astrocyte coculture AD model. **A.** Immunofluorescence confocal deconvolved image shows iPSC-derived human neurons (red, neurofilament H) robustly expressing pre- and post-synaptic markers (green, Synapsin1/2; grey, PSD95) at day 150 of coculture. Scale bar 2.5 μm. **B-C.** Immunofluorescence confocal image shows neuronal cell bodies and dendritic processes (green, MAP2), as well as axonal processes (red, SMI312) of iPSC-derived human neurons. (**B**) A low-zoom field of view (FOV). Scale bar 50 μm. (**C**) Two high-zoom FOVs showing dendritic and axonal processes. Scale bar 5 μm. **D.** Immunofluorescence confocal image shows the presence of both neurons (grey, SMI312) and astrocytes (red, S100b) in the coculture. Scale bar 50 μm. **E.** Immunofluorescence confocal images show 6e10 positive (grey) and ThioflavinS positive (blue) amyloid beta deposits formed in human iPSC-derived AD model following treatment with amyloid beta 1-42 peptides. Axonal processes were labeled with neurofilament (NFH, red). Scale bar 5 μm. **F.** Immunofluorescence confocal image shows axonal processes formed spheroids (NFH, red) around amyloid plaque deposit (ThioflavinS, blue). Scale bar 5 μm. **G.** Immunofluorescence confocal deconvolved images show phosphorylated Tau S235, S396 and S404 (red) expression in PAAS derived from human neurons (grey, SMI312). Zoom out images were maximum projected, while the zoom in images show a single plane. Scale bar 5 μm. **H-I.** Immunofluorescence confocal deconvolved images of AAV2-CB7-GFP infected (**H**) human neurons with abundant axonal spheroids (green, anti-GFP staining), co-stained with the axonal spheroid marker SMI312 (grey); **(I**) cell bodies of human neurons, as revealed by both anti-GFP (green) and anti-NeuN (grey) staining, related to Figure 6O. Scale bar (**H**) 5 μm and (**I**) 50 μm.

**Figure S17.**
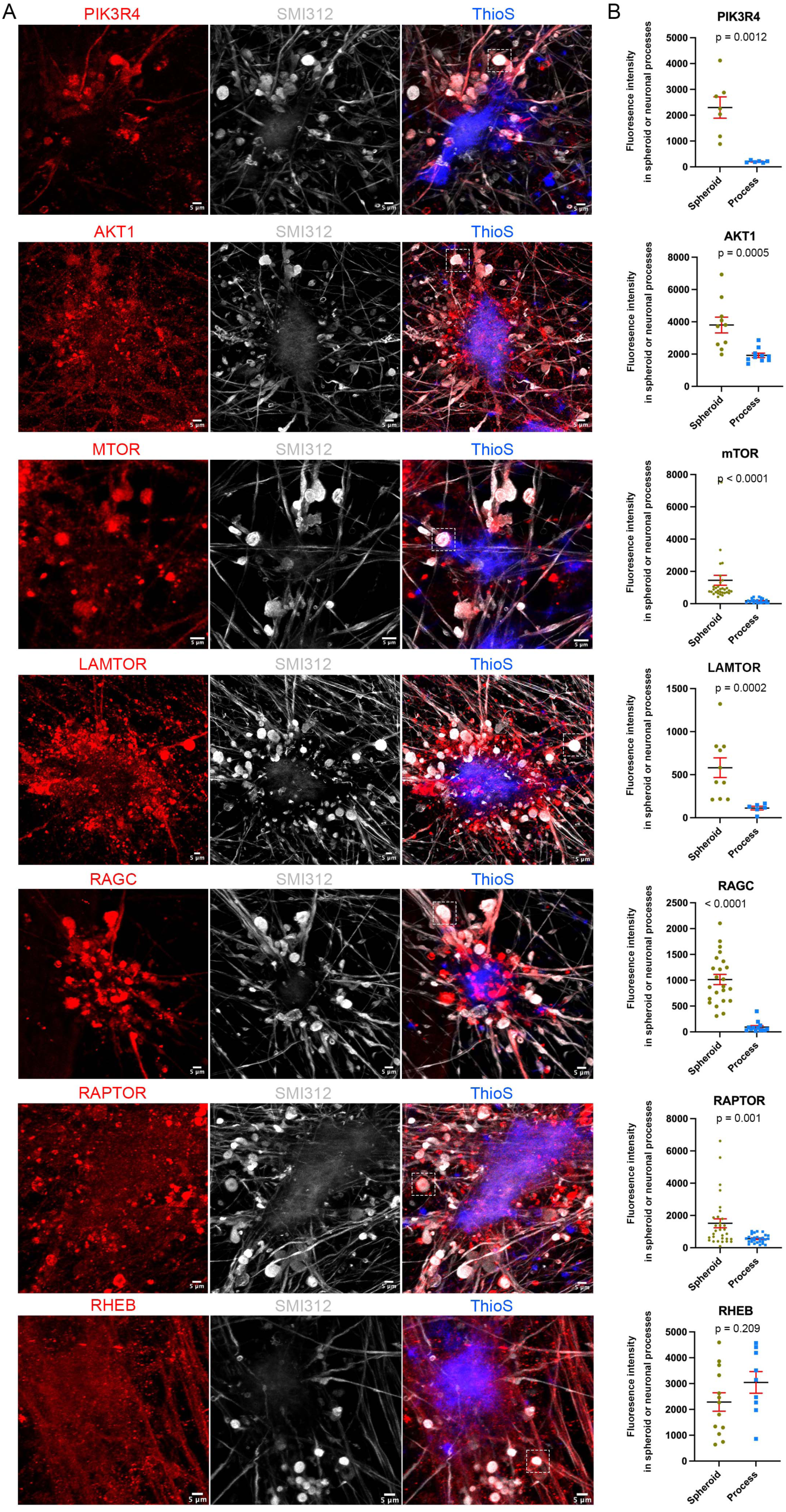
Immunofluorescence confocal imaging of the PI3K/AKT/mTOR signaling in human iPSC-derived neuron-astrocyte coculture AD model. **A.** PI3K/AKT/mTOR signaling molecules (red) are expressed in hiPSC-derived axonal spheroids (grey, SMI312) around amyloid plaque (blue, ThioflavinS). Spheroids in the white boxes in each large field-of-view image are shown as cropped images in Figure 6I. Scale bar 5 μm. **B.** Quantification of fluorescence intensity of protein hits in axonal spheroids versus neuronal processes. Each dot represents an ROI from three independent experiments. Unpaired non-parametric t test (Mann-Whitney) was performed. Error bars indicate SEM.

**Figure S18.**
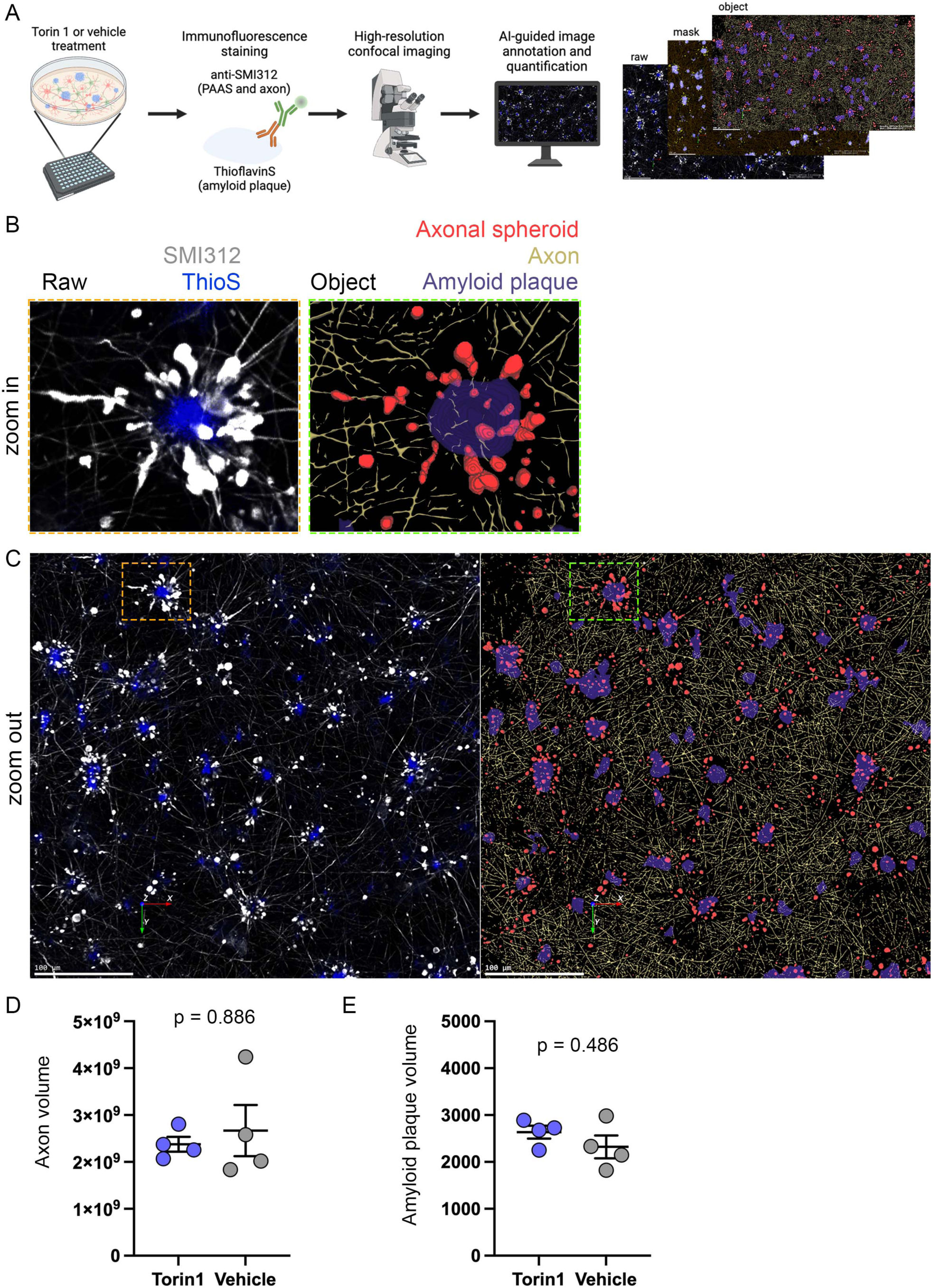
High throughput automated quantification of axonal spheroids, axons and amyloid plaques in human iPSC-derived AD model. **A.** Schematic showing the workflow of immunofluorescence labeling of axonal spheroids, axons and amyloid plaques in human iPSC-derived AD model, followed by confocal imaging and machine learning-based image analysis and quantification. **B-C.** Zoom in (**B**) and zoom out (**C**) images of immunofluorescence confocal imaging showing SMI312 antibody labeled axonal spheroids and axons (white), ThioflavinS labeled amyloid plaque (blue). Objects of axonal spheroids (red), axons (yellow) and amyloid plaques (purple) were generated according to the raw images after image annotation and analysis. Scale bars = 100 μm. **D-E.** Quantification showing (**D**) axon and (**E**) amyloid plaque volume (related to the experiment in Figures 6Q and **6R)**. Unpaired t test non-parametric two-tailed Mann Whitney test was used for all the statistical analysis.

**Figure S19.**
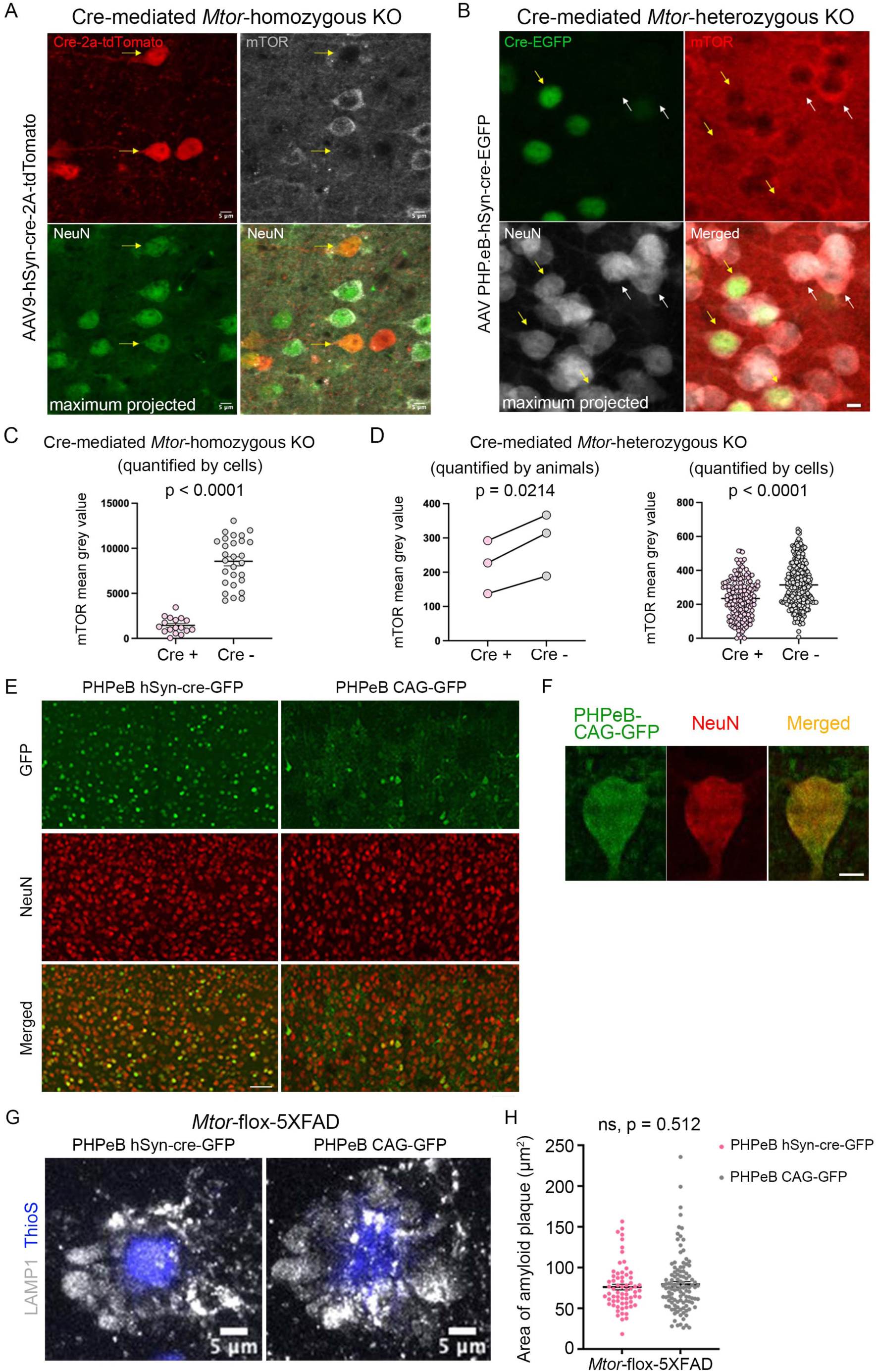
Genetic manipulation of mTOR signaling in mice. **A.** Uncropped immunofluorescence confocal images (shown in Figures 7B and **C**) show AAV9-hSyn-cre-2A-tdTomato-mediated *Mtor* complete knockout in homozygous *Mtor*-flox-5XFAD mice. mTOR expression (grey) was absent in Cre expressing neurons (red), compared to other adjacent neurons (green, NeuN) without Cre expression. **B.** AAV PHPeB-hSyn-cre-EGFP-mediated mTOR heterozygous knockout in heterozygous mTOR-flox-5XFAD mice. mTOR expression (red) was reduced in Cre expressing neurons (green), compared to other adjacent neurons (grey, NeuN) without Cre expression. **(A-B)** Scale bar = 5 μm. **C-D.** Quantification of mTOR fluorescence intensity (mean grey value) showed both (C) homozygous and (D) heterozygous mTOR knockout significantly reduced mTOR expression. Cre-virus positive (knockout) cells were compared to the Cre-virus negative (not knockout) cells in the same field of view. Cell bodies were outlined by NeuN staining. *Mtor*-floxed homozygous mouse (n=1), *Mtor*-floxed heterozygous mice (n=3). **E.** Immunofluorescence confocal images show AAV PHPeB virus (green) infection efficiency in the somatosensory cortex of 5XFAD mice. Anti-NeuN staining (red) indicate neuronal cell bodies. Scale bar = 50 μm. **F.** Immunofluorescence confocal image showing NeuN (red) positive neuronal soma (green, AAV PHPeB-CAG-EGFP infected) used for soma size quantification in Figure 7K. Scale bar = 5 μm. **G.** Representative images of axonal spheroid halos in heterozygous mTOR-flox-5XFAD mice under PHPeB-hSyn-cre-GFP (knockout) or PHPeB-CAG-GFP (not knockout) virus transduction. Spheroid halos were stained by LAMP1 (grey) and amyloid plaques were stained by ThioflavinS (blue). **H.** Quantification of the size of individual amyloid plaques within each ROI. Each dot represents a single amyloid plaque. *Mtor* knockout group n = 66 plaques, control group n = 109 plaques from 3 mice in each group. Unpaired t-test, two-tailed, p = 0.512.

**Figure S20.**
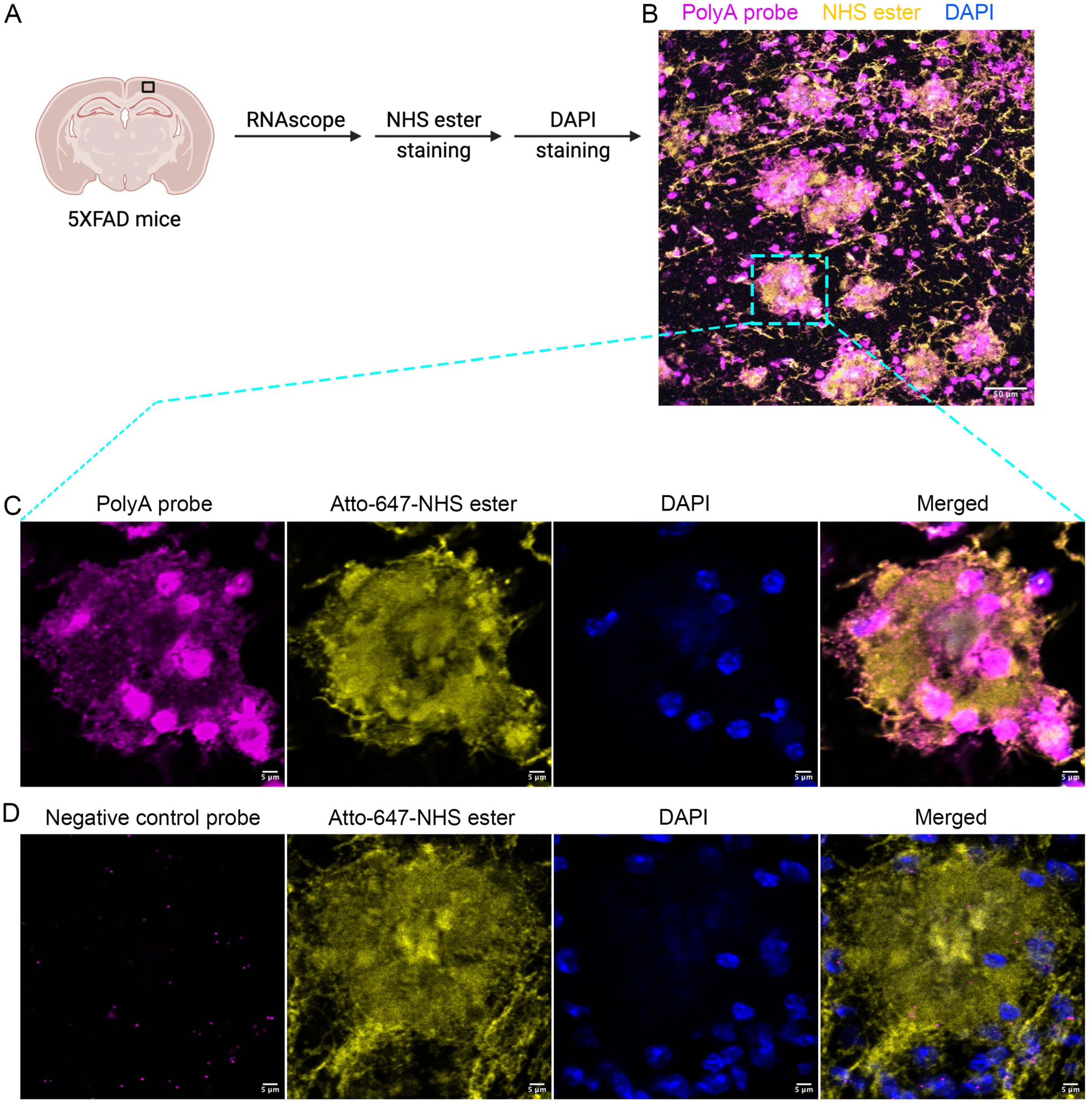
RNAscope revealed mRNA localization in axonal spheroids in 5XFAD mice. **A.** Schematic showing the RNAscope and staining pipeline in 5XFAD mice brain sections. **B-C.** (**B**) Low zoom and (**C**) high zoom (insert, blue box) RNAscope images showing mRNA species (PolyA probe labeled, magenta) are highly expressed in cell bodies (DAPI, blue) and are expressed within axonal spheroid halos (NHS ester labeled, yellow). NHS ester (yellow) labels axonal spheroid halo (including amyloid plaques), blood vessels and neuropil in 5XFAD mice cortex. **D.** High zoom image of negative control probe RNAscope in 5XFAD mice cortex. The RNA probe signal was mostly abrupted in both cell bodies (DAPI, blue) and axonal spheroid halos (NHS ester, yellow). Scale bar (**B**) 50 μm, (**C-D**) 5 μm.

**Figure S21.**
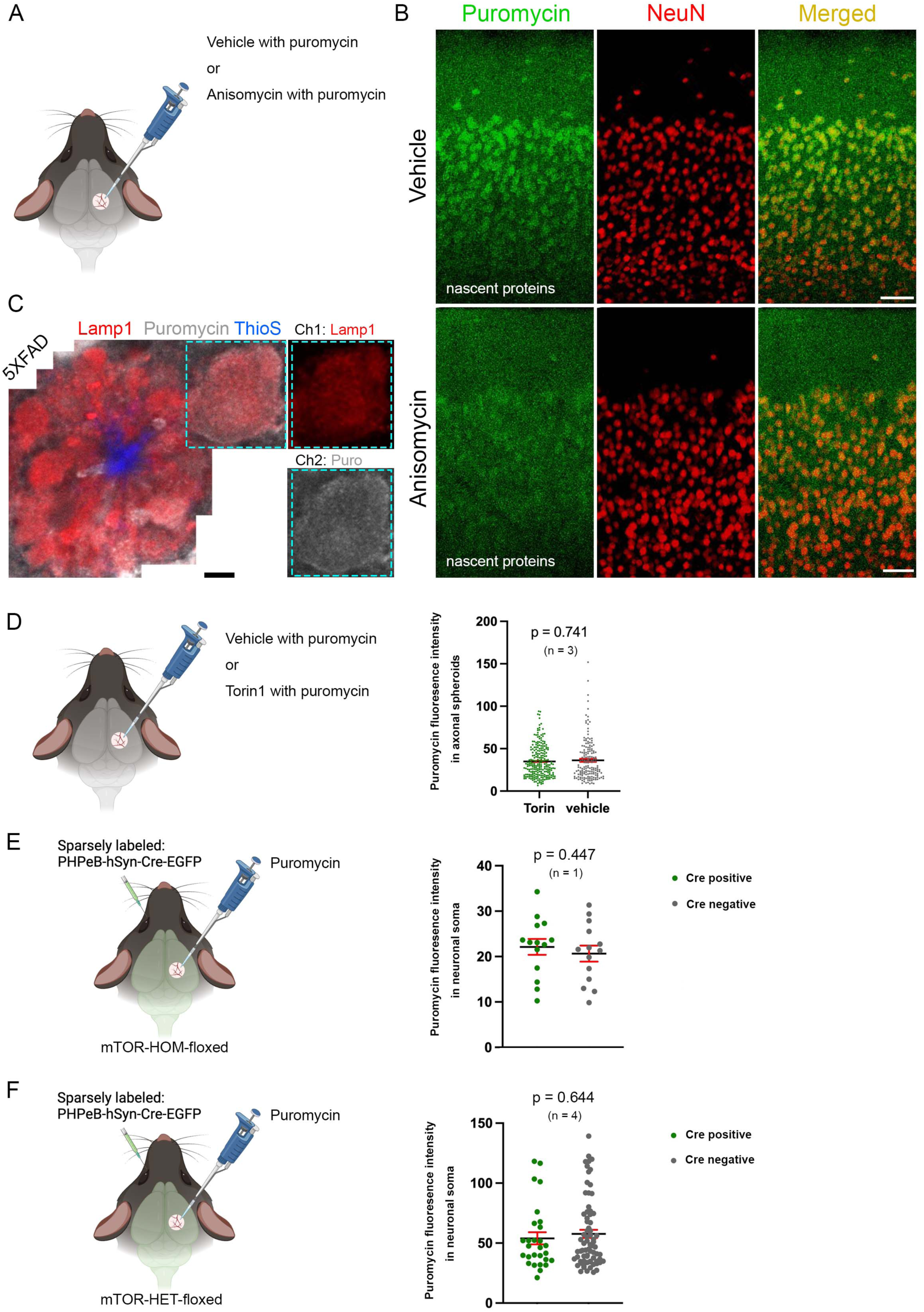
Puromycylation revealed nascent peptide in axonal spheroids in 5XFAD mice. **A.** Schematic showing puromycylation or anisomycin inhibition in mouse cortex. **B.** Immunofluorescence confocal images showing nascent proteins (puromycin labeled, green) in mouse cortex. Neuronal soma were labeled by NeuN (red). Anisomycin inhibition reduced the degree of puromycin labeling. **C.** Representative images showing nascent proteins (puromycin labeled, grey) in axonal spheroids (Lamp1, labeled) around amyloid plaque (ThioflavinS labeled, blue) in 5XFAD mice. Scale bar (**B-C**) 5 μm. **D.** Quantification showing 700 μM Torin treatment for two hours did not have an impact on puromycin labeling in axonal spheroids of 5XFAD mice cortices. Each dot represents an axonal spheroid, from n = 3 mice. **E-F.** Schematics showing (**E**) homozygous knockout or (**F**) heterozygous knockout of mTOR, by transducing PHP.eB-hSyn-Cre-GFP viruses in (**E**) mTOR-heterozygous-floxed or (**F**) -homozygous-floxed mice cortices. Quantification showing (**E**) homozygous knockout or (**F**) heterozygous knockout of mTOR in neurons did not have an impact on puromycin labeling in neuronal soma. Each dot represents a Cre positive, or Cre negative neuronal cell body (NeuN labeled) from (**E**) mTOR-HOM-floxed mouse (n = 1), and (**F**) mTOR-HET-floxed mice (n = 4). Unpaired t-test, two-tailed was performed.

### SUPPLEMENTARY TABLES

**Table S1:** Raw and analyzed proteomics results of PAAS, Lamp1-labeling proximity labeling proteomics and neuronal soma in humans and mice.

**Table S2:** Lists of newly identified and known PAAS proteins, as well as details and instructions of antibodies and immunofluorescence staining.

**Table S3:** Proteomic sample information (also available at Proteomexchange).

**Table S4:** Pathway enrichment analysis of Gene Ontology of PAAS proteomes in AD humans and mice.

**Table S5:** IPA signaling pathways of PAAS proteomes in AD humans and mice.

**Table S6:** GSEA analysis of PLD3-labeled proteomes in AD humans v.s. unaffected control humans.

### SUPPLEMENTARY MOVIE CAPTION

**Movie S1:** A 3D video captured using FIB-SEM from a 12-month-old 5XFAD mouse shows spheroids (magenta) filled with a large number of vesicles, located around amyloid plaques (cyan).

